# Taxonomic Revision and Identification Keys for the Giant Honey Bees

**DOI:** 10.1101/2024.04.03.587895

**Authors:** Nyaton Kitnya, Axel Brockmann, Gard W. Otis

## Abstract

The taxonomy and phylogeny of the giant honey bees (*Apis*; subgenus *Megapis*) are still controversial and unresolved. The species boundaries within the subgenus are unclear and some species that are recognized on the basis of genetic differences lack supporting morphological characters. Two species are now well accepted: *Apis dorsata* F. of tropical regions of Asia and *Apis laboriosa* Smith that inhabits the foothills of Himalaya and neighboring mountain ranges. In addition, researchers have suggested that the two allopatric populations of giant honey bees that inhabit Sulawesi, Indonesia (*A. binghami* Cockerell) and the oceanic Philippine islands (*A. breviligula* Maa) as well as the South Indian form also deserve species status. We conducted a taxonomic study based on morphological characters of *Megapis* from throughout Asia. Our study confirms that *Apis laboriosa* is a distinct species based on numerous morphological characters. Moreover, *A. dorsata* of mainland Asia differs from the two island taxa based on coloration, ocellus size, and the spacing of compound eyes and ocelli. We found no evidence that *breviligula* of the Philippines has a distinctively short tongue, and report only one minor character (the shape of sternum 5) that differed statistically between bees from Sulawesi and the Philippines. We conclude that the bees from these islands represent a single morphological species, *A. binghami*, with two subspecies, *A. b. binghami* and *A. b. breviligula*. *Apis dorsata* from the Andaman Islands are smaller than but conspecific with those of mainland Asia. We found no morphological autapomorphies in the giant honey bees of southern India known to differ in mtDNA from *A. dorsata* from elsewhere in mainland Asia. We provide a taxonomic keys to workers and drones within the subgenus *Megapis*.

## 1 Introduction

The taxonomy and phylogeny of the giant honey bees (*Apis* L.; subgenus *Megapis* Ashmead) are still controversial and poorly resolved, partly due to a paucity of studies. In his seminal review of the honey bees, Maa (1953) recognized four species of giant honey bees within the genus *Megapis*, viz *dorsata* Fabricius 1793, *laboriosa* Smith 1871, *binghami* Cockerell 1906, and *breviligula* Maa 1953. However, his classification was criticized by many in the research community for excessive splitting of the genus *Apis* into three genera and 24 species (including 12 species within the taxon currently recognized as *A. mellifera*) (Ruttner, 1988; Engel, 1999). Rather than rejecting the systematic revision by Maa (1953), Sakagami et al. (1980) examined in detail the morphological differences between workers of the two mainland Asian forms, *dorsata* and *laboriosa.* Examining more than a hundred morphological features, they found several discrete characters that distinguish *laboriosa* from *dorsata*. Unfortunately, they had only 2 specimens of *breviligula* and none of *binghami* to examine and therefore they did not comment on their status.

In 1991, Alexander had the opportunity to examine both workers and drones of all four giant honey bee taxa. He examined 18 morphological and two behavioral characters to determine the cladistic relationships of the species within *Apis*. Unfortunately, none of the characters he assessed resolved the phylogenetic relationships among the giant honey bees. Therefore, he treated *dorsata, laboriosa*, *binghami* and *breviligula* as the “*dorsata* group”.

Despite Sakagami et al. (1980) recognizing several diagnostic characters that differed strikingly between *dorsata* and *laboriosa*, until now most taxonomists who have studied bee taxonomy (e.g., Ruttner, 1988; Alexander, 1991; Engel, 1999) remained unconvinced that *laboriosa* deserved species status and adopted a conservative view that there is a single giant honey bee species, namely *Apis dorsata*. They considered that *laboriosa*, *binghami* and *breviligula* are distinctive populations within *A. dorsata* that occur in circumscript geographical areas. Specifically, *dorsata* is broadly distributed throughout Southeast Asia and the Indian subcontinent, from as far west as Afghanistan to Timor and possibly the Kei Islands of Indonesia in the east (Maa, 1953). *laboriosa* prevails across relatively high elevations of the pan-Himalaya region and smaller mountain ranges extending eastward into China, Laos, and Vietnam, southward into central Myanmar, and westward into Pakistan (Kitnya et al., 2020; Otis et al., submitted). *binghami* is found only on the island of Sulawesi and nearby islands of Indonesia and *breviligula* occurs on the islands of the Philippines, excluding Palawan (Smith, 2021; Bhatta et al., in prep.).

The phylogenetic relationships resulting from genetic studies are suggestive of four, possibly five, species of the giant honey bees, although tree topologies differ among the studies. Most phylogenetic relationships inferred from molecular analyses have reported *laboriosa* as sister to the other taxa (Arias and Sheppard, 2005; Raffiudin and Crozier, 2007; Wang et al., 2018; Bhatta et al., in prep.). However, Lo et al. (2010), based on analyses of both mitochondrial and nuclear DNA sequences, reported that *breviligula* of the Philippines diverged prior to splits among the other taxa.

Their results showed *laboriosa* (with low support value) closer to *dorsata* than *breviligula,* while *binghami* was not clearly resolved either from *dorsata* or *laboriosa*. In contrast, the phylogenetic inferences of Raffiudiun and Crozier (2007) and Smith (2021) showed *dorsata* and *binghami* as highly resolved clades with high support values. These topological differences within phylogenetic trees result from different analytical methods and portions of the genome examined.

Lo et al. (2010) and Bhatta et al. (in prep.) are the only researchers who have included all of the giant honeybee taxa in their analyses, unlike others who omitted one or more taxa. Lo et al. reported that *dorsata* from Tamil Nadu and Karnataka in India and Sri Lanka had diverged from the common ancestor of *dorsata* of Southeast Asia and formed a distinct clade within their phylogenies, suggesting that the giant honey bees of South India and Sri Lanka may deserve species status, a result first proposed by Smith (1991) and echoed in the results of Arias and Sheppard (2005). Our recent study (Kitnya et al., 2022) also showed two distinct clades of *dorsata* from Karnataka (South India) and Arunachal Pradesh (North East India) based on COI mtDNA. In the most complete analysis of mtDNA sequences to date, Bhatta et al. (in prep.) provide support for *laboriosa* as well as four species within the *dorsata* complex: *dorsata*, *binghami*, *breviligula*, and the South Indian form.

Morphological reanalysis of specimens of *Megapis* taxa throughout their ranges will yield significant information to help resolve the taxonomy of the giant honey bees. Some of the species recognized on the basis of genetic differences (Smith, 1991; 2021; Arias and Sheppard, 2005; Raffidiun and Crozier, 2007; Lo et al., 2010; Cao et al., 2012; Bhatta et al., in prep.) lack data for previously stated morphological characters (Maa, 1953; Sakagami et al., 1980; Trung et al., 1996). Without that supporting evidence, elevating taxa to species status is somewhat controversial on the basis of genetic differences alone, especially when the phylogenies based on those genetic analyses are not congruent.

To resolve the species boundaries within *Megapis* bees, we have examined specimens from across the ranges of all the taxa (Fig. 1) and quantified numerous characters. Here we describe those characters and create taxonomic keys for the identification of both workers and drones. Hereafter, for the sake of convenience, while describing each taxon within *Megapis*, Maa’s (1953) names—*dorsata, laboriosa, binghami* and *breviligula*—will be used, though the study will reassess the validity of these names based on present research. Fortunately, we had specimens of both workers and drones from many localities, unlike the limited series of worker specimens only on which Maa (1953) and Sakagami et al. (1980) based their taxonomic evaluations.

**Figure 1.**
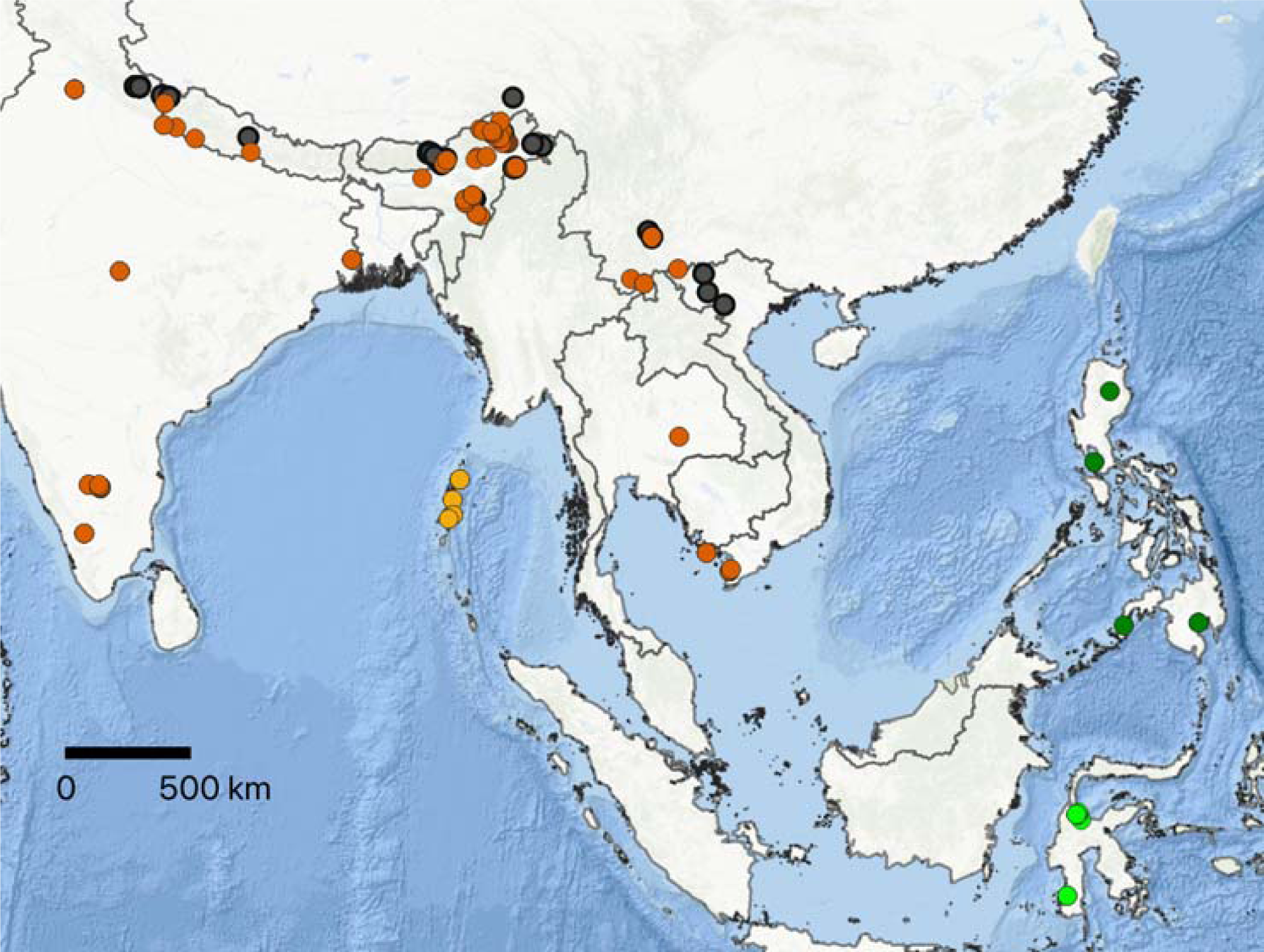
Map showing collection localities of *Megapis* examined. Neon green circles: *binghami*; Dark green circles: *breviligula*; Orange circles: *dorsata* (mainland); Mustard yellow circles: *dorsata* (Andaman Island); Black circles: *laboriosa*. Color codes are consistent throughout the manuscript.

We hypothesized that the four *Megapis* species recognized by Maa would be upheld. The striking and distinctive morphological characters of *laboriosa* reported in previously studies (Maa, 1953; Sakagami et al., 1980; Trung et al., 1996; Kitnya et al., 2022) suggested that it deserved distinct species status from *dorsata* of mainland Asia. The genetic and morphometric analyses of sympatric *laboriosa* and *dorsata* of Kitnya et al. (2022) further demonstrated the species status of *laboriosa*. The giant honey bees, *binghami* and *breviligula*, from the islands of Sulawesi, Indonesia and the Philippines, respectively, are similar in appearance to each other. As described by Maa (1953), these black island giant honey bees superficially resemble *laboriosa* with which they were earlier sometimes confused (e.g., Cockerell, 1914). Based on the distinctive short tongue and other distinctive characters of *breviligula* reported by Maa (1953), we hypothesized that the black island taxa represent two separate species. We were unsure what our results might reveal about the South Indian population, given that *dorsata* of South India and Southeast Asia are very similar in external appearance and behavior. However, as reviewed above, in molecular phylogenies they form well resolved clades (Smith, 1991, 2021; Lo et al., 2010; Kitnya et al., 2022).

## 2 Materials and Methods

### 2.1 Collection and processing of specimens

Collections of the giant honey bees were carried out over four years (2017 to 2021) by NK and her colleagues in different parts of India: Andaman Islands, Arunachal Pradesh, Delhi, Karnataka, Madhya Pradesh, Manipur, Nagaland, Punjab, Tamil Nadu, Telangana, Uttarakhand, and West Bengal. Additionally, we examined specimens from China, Nepal, Thailand, Vietnam, Indonesia, and the Philippines. See Supplementary File 1 for locations of the specimens we examined and the names of collectors.

During field trips, bees were collected with an aerial insect net while they foraged on flowers. Generally, they were killed in a tube containing few drops of ethyl acetate on a ball of cotton; however, some bees killed by being placed directly in 70% ethanol were also included. Dead bees with locality and date information were transferred to packets made from butter paper (transparent paper) for temporary storage in the field. All bees collected were pinned with number 2 insect pins and labelled with collection details. These specimens were then dried in a warm oven at 35° C for 72 hours and deposited in an insect cabinet in the NCBS-TIFR Research Collection Facility for further study. Photographs of each specimen along with label details were taken to create a digital database, available at: https://www.ncbs.res.in/research-facilities/museum-fieldstations).

The specimens studied include bees of all four named taxa as well as the South Indian form. These include *laboriosa* from India (Arunachal Pradesh, Nagaland and Uttarakhand), Nepal, China, and Vietnam; *dorsata* from India (Andaman Islands, Arunachal Pradesh, Delhi, Karnataka, Madhya Pradesh, Manipur, Nagaland, Punjab, Tamil Nadu, Telangana, Uttarakhand, West Bengal), Nepal, Thailand, China, and Vietnam; *binghami* from Central and South Sulawesi in Indonesia; and *breviligula* from Luzon and Mindanao islands in the Philippines (refer to Fig. 1 and Supplementary file 1 for collection details). The numbers of specimens examined and their provenance are given in Table 1.

**Table 1:**
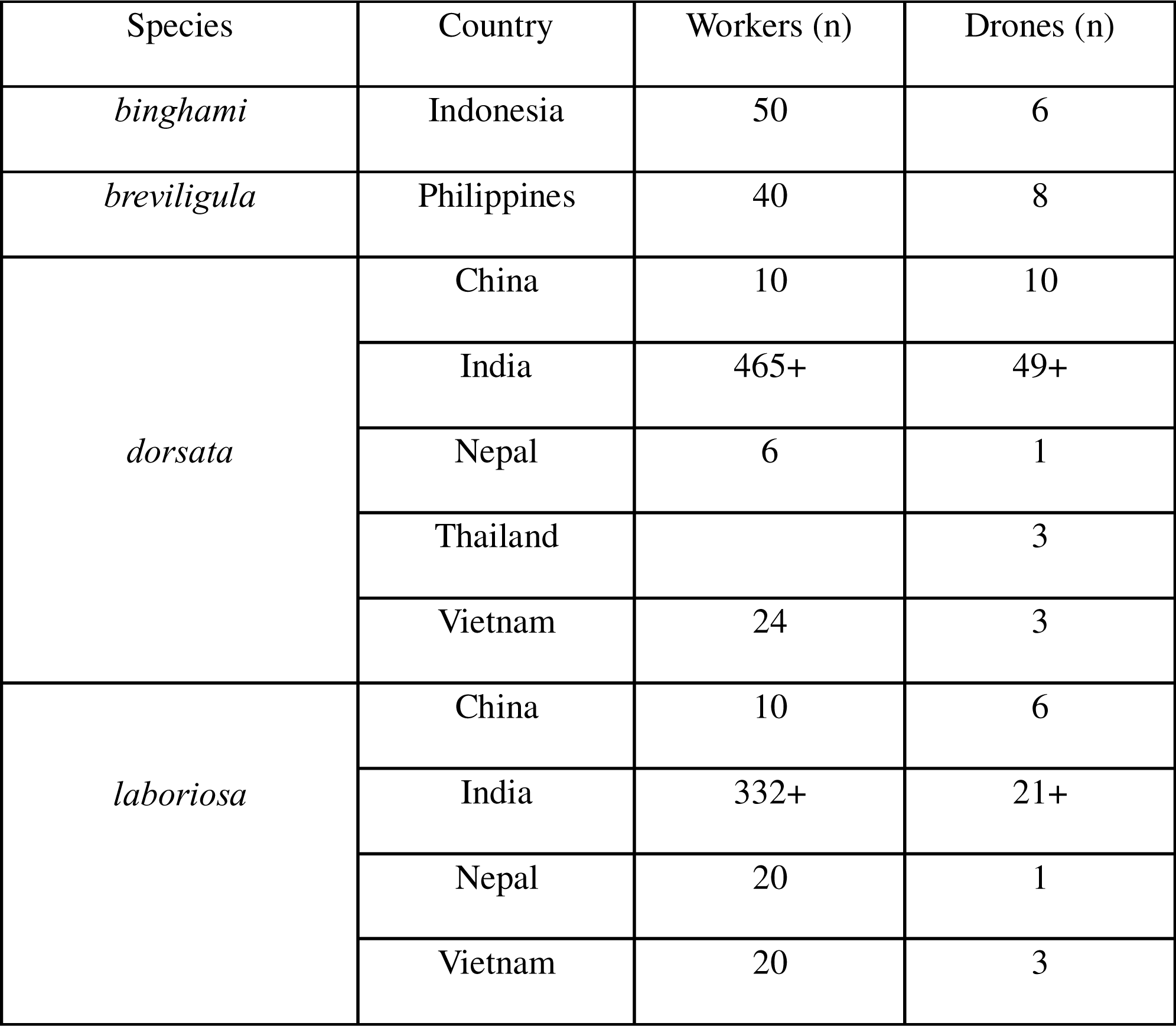
Sample sizes of *Megapis* specimens examined.

### 2.2 Dissection of specimens

For assessment of characters that required dissection, dried specimens were placed in a humidification chamber for 24–48 hours or boiled in 10% KOH for 4–5 mins. The mouthparts were removed with forceps. The metasomal terga were separated from the metasomal sterna. Soft tissues were removed with the help of a soft brush and then the sterna were carefully pulled apart. After the removal of soft tissues, the sterna and the 7^th^ hemitergite (spiracular plate) were washed with distilled water. The dissected metasomal sterna 2–6, spiracular plate (7^th^ hemitergite), and mouthparts were placed on a microscope slide with few drops of glycerine and examined under a dissecting microscope at various magnifications (sterna 2–5 at 1.6x; 7^th^ hemitergite at 4x, and mouthparts at 1.6x). Simultaneously, images were captured using a Leica M125 C stereomicroscope, with a dedicated Leica MC 190 HD camera and LAS V4.12 software to process images. While taking the images, another slide was placed over the body part so that it remained in a horizontal plane and properly spread. The body parts were stored in glycerine along with the respective voucher code in the NCBS-TIFR Research Collection Facility. The specimens of bees we dissected were a small subset of the total specimens available for study.

### 2.3 Terminology used in the study

The morphological terminology used in this study mostly follows Michener (1994, 2007) and Maa (1953). The abbreviations used are as follows: MOD = median ocellus diameter; IOD = interocellar distance; OOD = ocellocular distance (OOD); UOD = upper ocular distance; HW = head width; ML = malar length; MW = malar width; *i*-V = *indica vein*; L_1_ = tongue length (prementum + glossa); S_5_L = sternum 5 length; S_5_W = sternum 5 width; S_n_ = metasomal sternum number n; T_n_ = metasoamal tergum number n. All measurements, in mm, were taken using imageJ (http://imagej.nih.gov/ij).

## 3 Results

### 3.1 Species accounts

The general habitūs of *Megapis* workers and drones of four taxa are displayed in Figure 2. The sample sizes along with countries from where specimens were collected are given in Table 1. The means, SD and ranges of all character measurements are given in Table 2.

**Figure 2:**
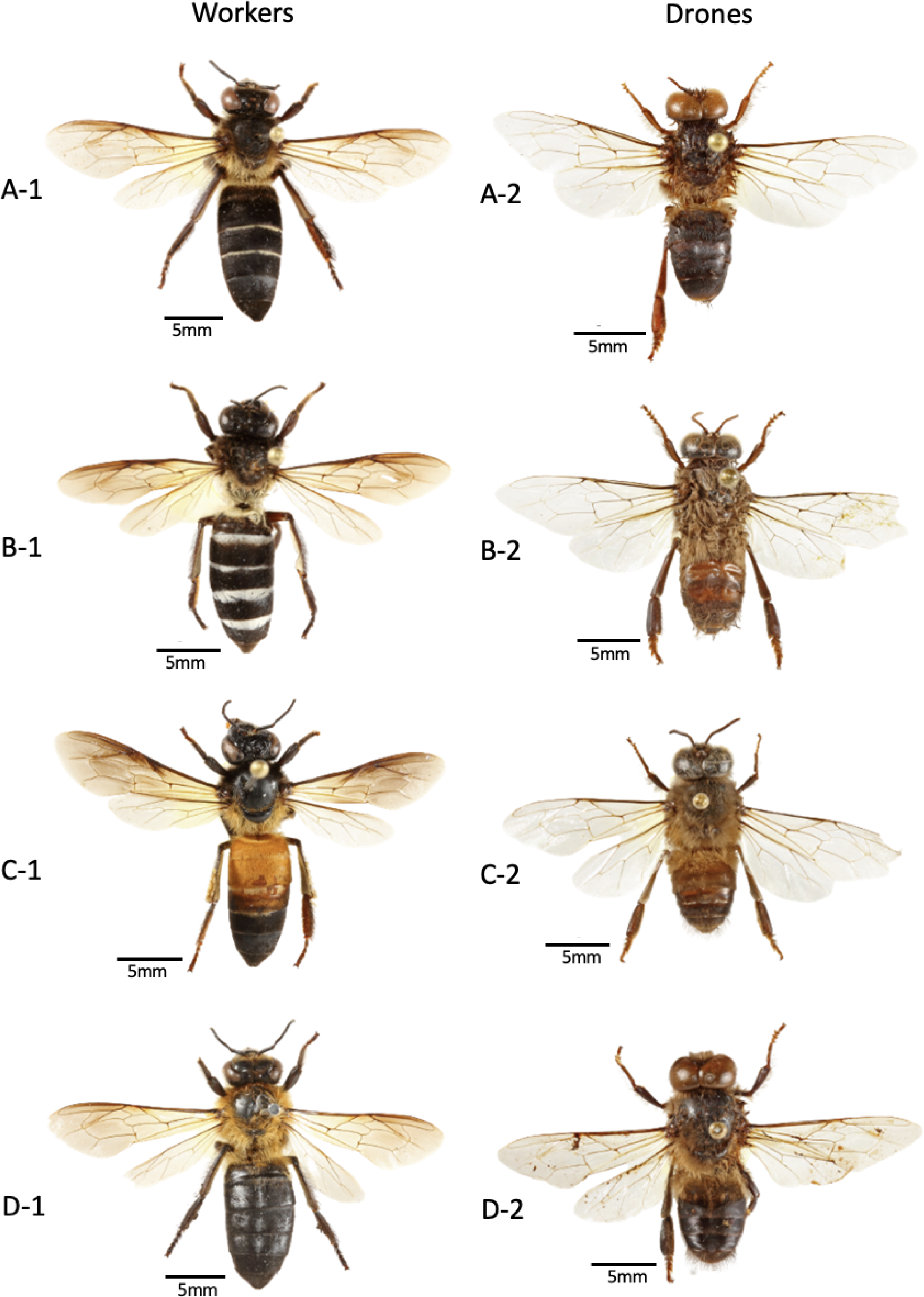
Dorsal habitus of *Megapis* (Ashmead, 1904). A. *binghami* (A-1: NRC-AA-3057, Palolo Valley, Rahmat, Sigi Regency, Central Sulawesi, Indonesia, 19ix1998, collector: G. W. Otis; A-2: NRC-AA-3070, Forestry Centre, Tabo Tabo, South Sulawesi, Indonesia, 3vi1989, collector: G. W. Otis); B. *breviligula* (B-1: NRC-AA-7668, Bulano ERTC, Philippines, July 2008, collector: Y.C. Su; B-2: NRC-AA-7665, Alfonso, Province of Cavite, Luzon, Philippines, Date 12-20vii2008, collector: Y. C. Su); C. *dorsata* (C-1: NRC-AA-7260, NCBS Campus, Bangalore, Karnataka, India, 27iii2023, collector: N. Kitnya; C-2: NRC-AA-7256, NCBS Campus, Bangalore, Karnataka, India, 28iii2023, collector: N. Kitnya); D. *laboriosa* (D-1: NRC-AA-2862, Xinping County, Sa Town, Yuxi City, Yunnan, China, 29iv2019, collector: Q. Lifeu; D-2: NRC-AA-2918, Kaski, Gandaki, Nepal, 8v1984, collector: B. A. Underwood).

**Table 2:** Means (mm), standard deviations, and ranges of characters measured. Median Ocellus Diameter (MOD); Interocellar Distance (IOD); Ocellocular Distance (OOD: Ratio of IOD and OOD); Ratio of Upper Ocular Distance (UOD) and Head Width (HW); Malar Length (ML); Malar Width (MW); *indica* Vein (*i*-V); Tongue Length (L_1_ = prementum + glossa); Sternum 5 Width (S_5_W); Ratio of S_5_L and S_5_W.

| Character | *binghami* |  |  | *breviligula* |  |  | *dorsata* |  |  | *laboriosa* |  |  |
| --- | --- | --- | --- | --- | --- | --- | --- | --- | --- | --- | --- | --- |
|  | Mean | SD | Range | Mean | SD | Range | Mean | SD | Range | Mean | SD | Range |
| MOD | 0.48 | 0.01<br>(n=30) | 0.46–0.52 | 0.47 | 0.01<br>(n=20) | 0.45–0.48 | 0.41 | 0.01<br>(n=105) | 0.40–<br>0.44 | 0.36 | 0.01<br>(n=60) | 0.34–0.39 |
| IOD | 0.46 | 0.04<br>(n=25) | 0.36–0.53 | 0.42 | 0.03<br>(n=20) | 0.38–0.48 | 0.43 | 0.03<br>(n=95) | 0.37–<br>0.51 | 0.38 | 0.03<br>(n=55) | 0.30–0.47 |
| OOD | 0.31 | 0.02<br>(n=25) | 0.27–0.35 | 0.31 | 0.02<br>(n=20) | 0.27–0.34 | 0.45 | 0.03<br>(n=95) | 0.38–<br>0.51 | 0.62 | 0.04<br>(n=55) | 0.55–0.74 |
| IOD/OOD | 1.49 | 0.16<br>(n=25) | 1.28–1.96 | 1.38 | 0.11<br>(n=20) | 1.19–1.52 | 0.96 | 0.08<br>(n=95) | 0.81–<br>1.16 | 0.60 | 0.05<br>(n=55) | 0.50–0.71 |
| UOD/<br>HW | 0.38 | 0.01<br>(n=29) | 0.36–0.41 | 0.39 | 0.01<br>(n=21) | 0.38–0.41 | 0.44 | 0.01<br>(n=105) | 0.42–<br>0.46 | 0.50 | 0.01<br>(n=55) | 0.48–0.52 |
| ML | 0.55 | 0.03<br>(n=27) | 0.50–0.61 | 0.60 | 0.04<br>(n=18) | 0.52–0.66 | 0.61 | 0.03<br>(n=112) | 0.52–<br>0.66 | 0.85 | 0.04<br>(n=48) | 0.75–0.96 |
| MW | 0.69 | 0.03<br>(n=27) | 0.59–0.75 | 0.68 | 0.02<br>(n=18) | 0.64–0.72 | 0.68 | 0.03<br>(n=112) | 0.59–<br>0.80 | 0.74 | 0.03<br>(n=48) | 0.65–0.80 |
| *i*-V | 0.49 | 0.1<br>(n=20) | 0.32–0.61 | 0.43 | 0.11<br>(n=16) | 0.33–0.62 | 0.52 | 0.12<br>(n=60) | 0.26–<br>0.73 | 0.93 | 0.23<br>(n=44) | 0.53–1.28 |
| $L_1$ | 6.34 | 0.12<br>(n=12) | 6.16–6.49 | 6.26 | 0.17<br>(n=13) | 5.93–6.46 | | | | | | |
| $S_5L$ | 1.77 | 0.32<br>(n=16) | 1.43–2.22 | 1.74 | 0.31<br>(n=16) | 1.35–2.12 | | | | | | |
| $S_5W$ | 3.09 | 0.59<br>(n=16) | 2.57–3.92 | 3.20 | 0.61<br>(n=16) | 2.51–3.92 | | | | | | |
| $S_5L/ S_5W$ | 0.57 | 0.02<br>(n=16) | 0.56–0.60 | 0.54 | 0.02<br>(n=16) | 0.59–0.57 | | | | | | |

#### 3.1.1 *Apis binghami* Cockerell, 1906

*Apis zonata* Smith, 1859: 8 (junior homonym) [*binghami* Cockerell]

*Megapis zonata* (Smith); Ashmead 1904: 121 [*binghami* Cockerell]

*Apis dorsata binghami* Cockerell 1906: 166. Replacement name for *Apis zonata* Smith 1859 [*binghami* Cockerell]

*Megapis binghami* (Cockerell); Maa 1953: 564 [*binghami* Cockerell]

*Apis dorsata binghami* (Cockerell); Engel 1999: 185 [*binghami* Cockerell]

Description of worker *A. binghami* (Fig. 2: A-1)

a. Integument: Integument generally black, except edge of labrum, mandibles, antennae, and legs that are iridescent brown.
b. Pubescence: Pubescence on head predominantly black except for short, whitish decumbent hairs on the paraocular area, and mixture of yellow-brown hairs on the genal area. Black-brown hairs on mesosoma (scutum and scutellum) except pale yellow hair on pronotum, lateral mesosoma, metanotum and propodeum. Distinct patch of brown hairs on mesopleuron. Pale-yellow hairs on the propodeum continues to the base of Tergum 1 (T1). However, black or sooty brown hairs covering a major portion of T1 continue to the end of the metasomal terga. Yellowish brown hairs on sternum 1 (S1), coxae, trochanters, femurs and tibiae. Visible band of short, decumbent whitish hairs at the anterior bases of T3, T4, and T5. There is some slight variation of color pattern among the individuals as described here. For example, in some specimens, such as those collected in ethanol (or with compressed metasoma after being killed), the bands of white hairs are not visible. This applies to the descriptions of other taxa as well.
c. Structures:
  I. Head: Ocelli arise from a prominently raised pedestal. Lateral ocelli much bulged and tilted toward compound eyes (S2: A). Median ocellus, 0.460–0.517 mm in diameter (Fig. 3A). Interocellar distance (IOD) is wider than ocellocular distance (OOD) (IOD = 0.359–0.531 mm; OOD = 0.269–0.351 mm). Ratio of IOD/OOD, 1.280–1.959 (Fig. 3B). Malar length (0.503–0.611 mm) slightly shorter than malar width (0.588–0.751 mm). Ratio of upper ocular distance (UOD) and head width (HW), 0.358–0.408 (Fig. 3C). Tongue length (prementum + glossa), 6.156–6.491 mm (Fig. 5A and Table 2).
  II. Metasomal abdomen: Anteglandulus of sternum 2 medially angulated, not arcuate (Fig. 4: A-1). Glandulus at the midpoint of sterna 3–5 curved posteriorly (see sternum 3 at Fig. 4: A-2). Ratio of length and width of sternum 5, 0.571–0.600. On 7^th^ hemitergite, posterior laminal spiracularis not invaginated (Fig. 4: A-3). The number of pairs of barbs on the sting, 10–11.
  III. Wing: *indica* vein on hind wing short, 0.324–0.613 mm. Callow individuals with grey integument and pale whitish hairs.

Description of drone *A. binghami* (Fig. 2: A-2)

**Figure 3:**
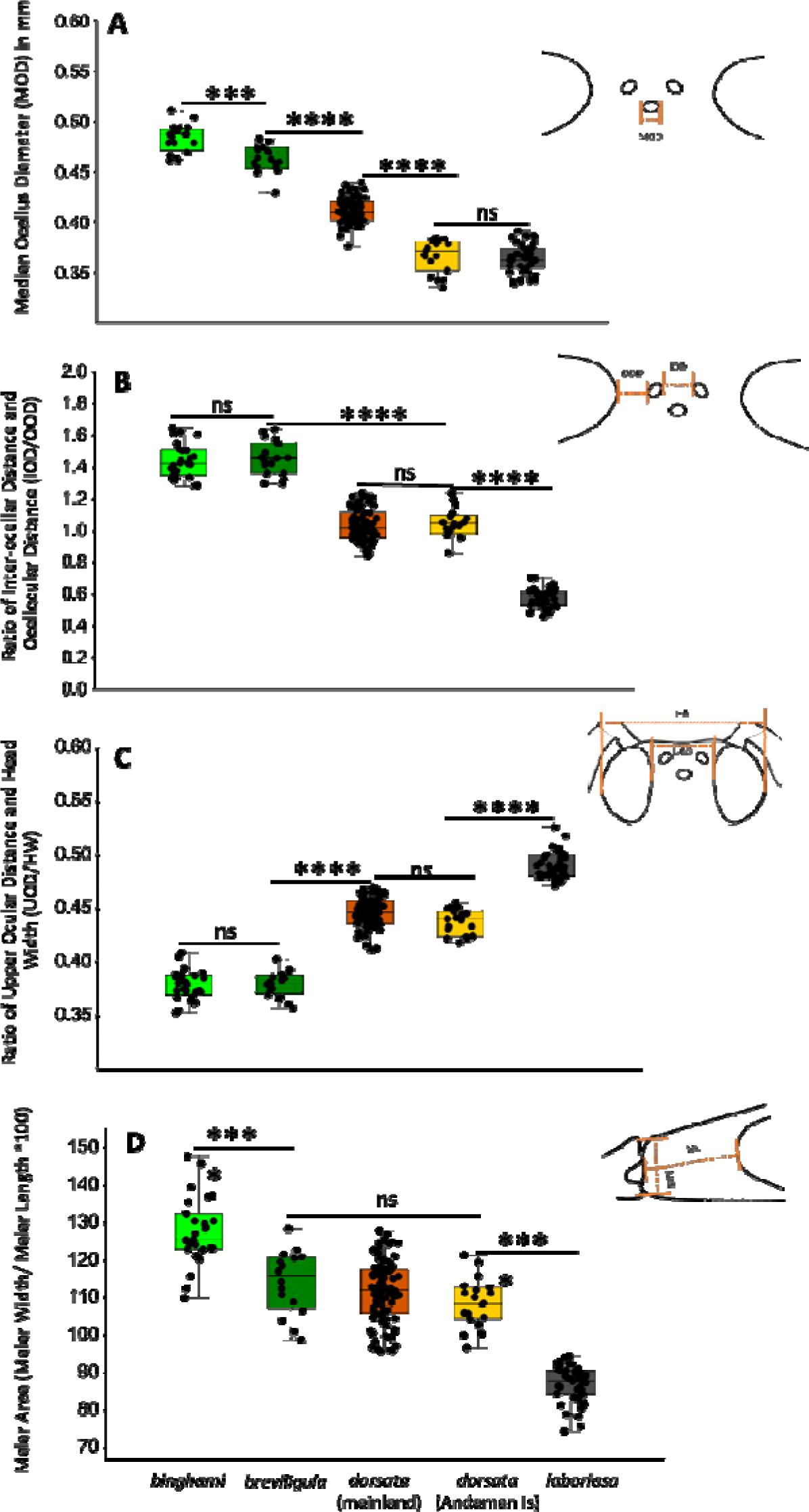
Head morphology of *Megapis* workers. A: Median Ocellus Diameter (MOD) in mm; B. Ratio of Interocellar Distance (IOD) and Ocellocular Distance (OOD); C. Ratio of Upper Ocular Distance (UOD) and Head Width (HW); D. Malar area: Malar Width (MW)/ Malar Length (ML) x 100. Significance of Tukey’s pairwise test: *, p ≤ 0.05; ***, p ≤ 001; ****, p ≤ 0.0001. Refer to supplementary file 1, Figures S2: A, S2: B and S2: C for comparative images of the traits mentioned here.

**Figure 4:**
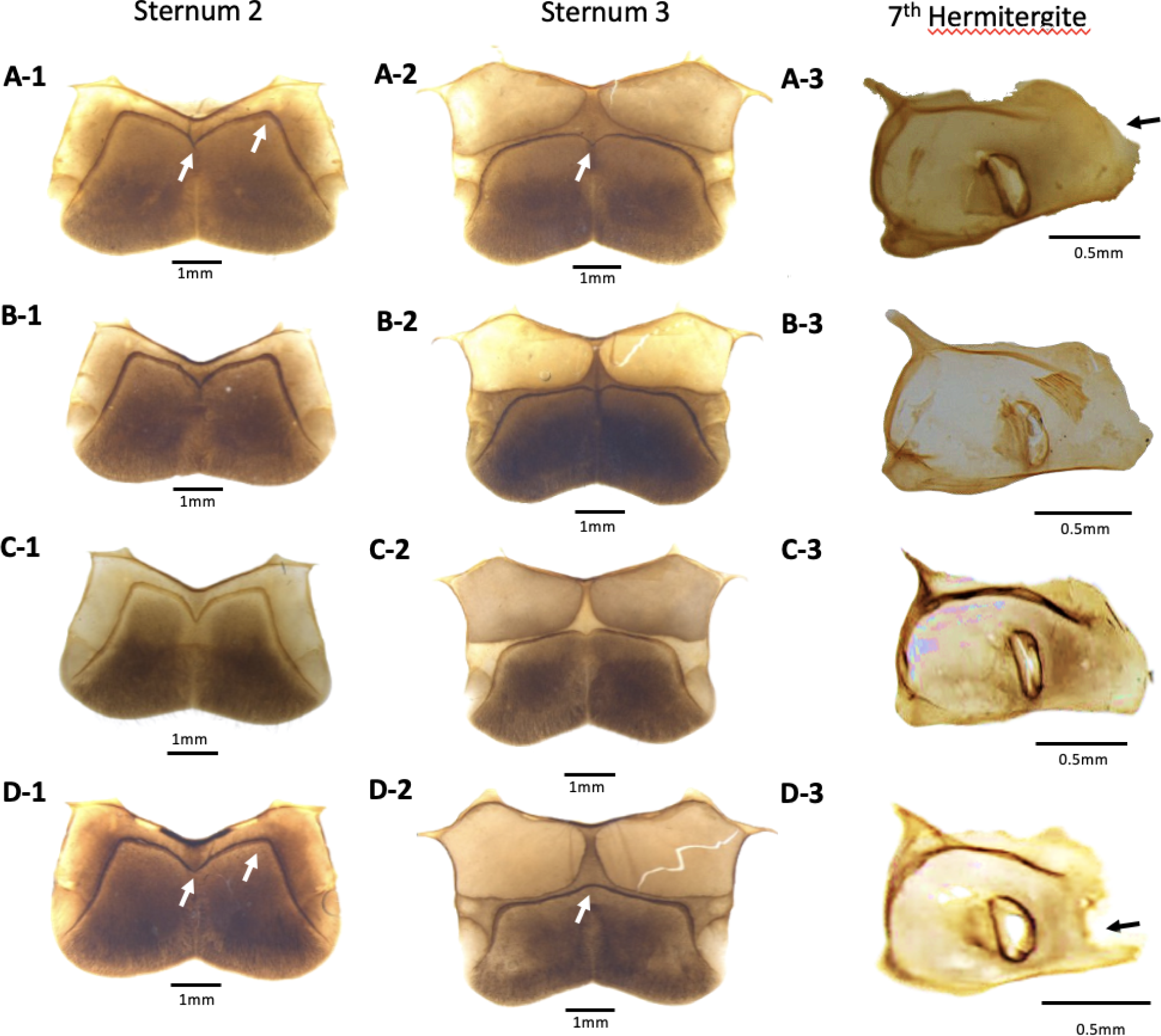
Abdominal morphology of *Megapis* workers. A: *binghami*; B: *breviligula*; C: *dorsata*; D: *laboriosa*. Arrows on sternum 2 indicate locations of differences in anteglandular structure; Arrows on sternum 3 indicate differences at mid-point of glandulus; Arrow on 7^th^ hemitergite indicates invagination of spiracular lamella at posterior edge in *laboriosa* that is absent in the other taxa.

**Figure 5:**
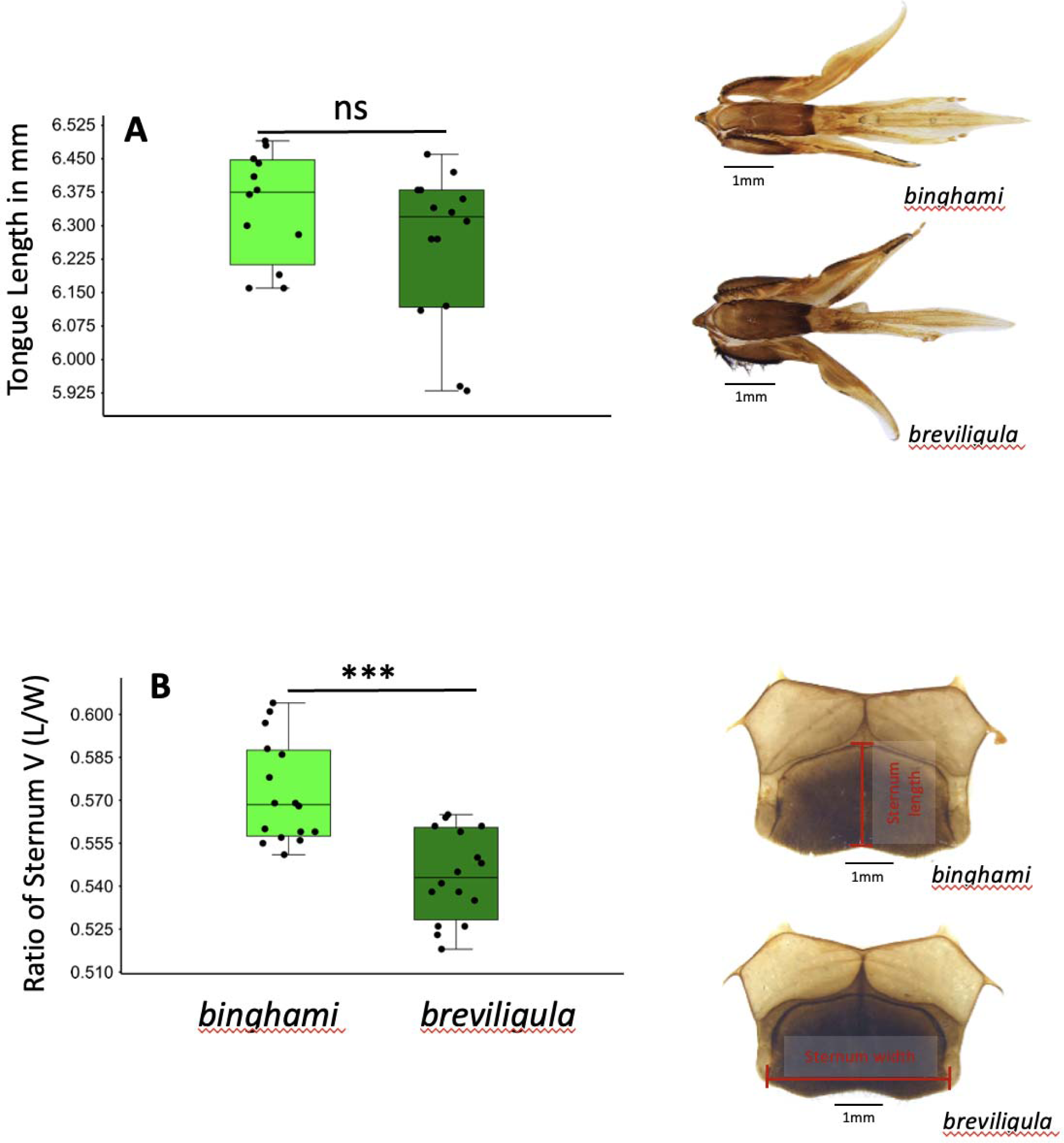
Tongue length and proportions of Sternum 5 of *binghami* and *breviligul*a workers. A. Tongue lengths (mm). B. Ratio of sternum 5 length to width. Significance of t-tests: ns = non-significant (p > 0.05); ***, p ≤ 0.001.

a. Integument: Integument of head, antennae, mesosoma and metasoma iridescent brown; antennae and legs brown.
b. Pubescence: Pale yellow hairs on face, eyes, gena, scutum, scutellum, lateral mesosoma, metasomal tergum 1, ventral mesosoma, sterna and legs. Brown hairs on metasomal tergum 2 to apex of metasoma; long, stiff, brown hairs on metasoma from tergum 5 onward to apex of metasoma. Clumps of light brown hairs on basitarsi III.
c. Structures: Ocellar triangle prominently raised (Fig. 6: A-1 & A-2); in fact, it is the most prominently raised among the giant honey bee taxa. Lateral ocelli close together; with almost no space between lateral ocelli and compound eyes (Fig. 6: A-2). Ocelli protrude strongly; lateral ocelli angled towards compound eyes. In dorsal view, ocelli extend prominently beyond the plane of compound eyes (Fig. 6: A-3).

**Figure 6:**
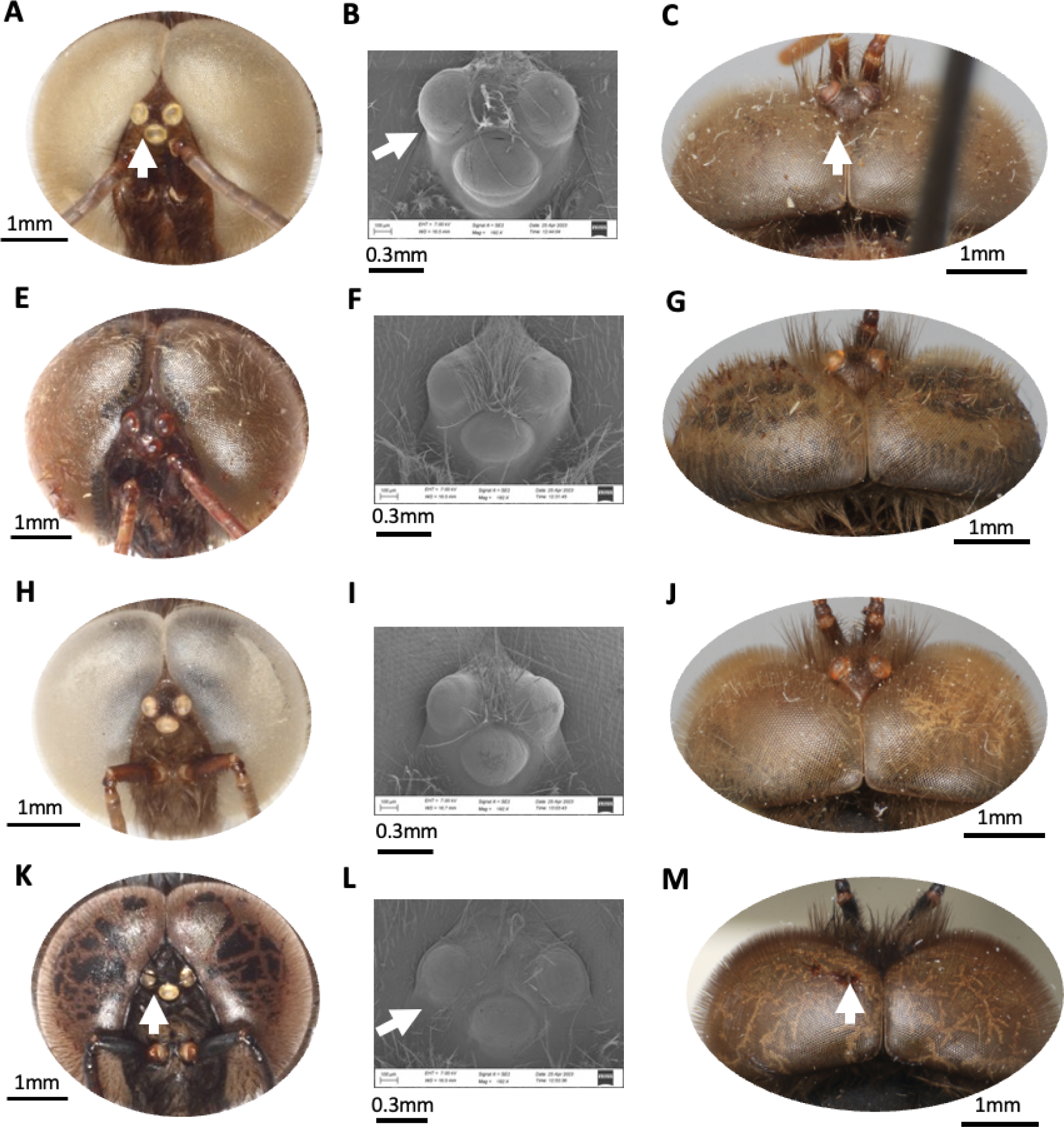
Head morphology of *Megapis* drones. A: *bingham;* B: *breviligula*; C: *dorsata*; D: *laboriosa*. Column 1 and 2: Position of ocellar triangle, the ocellar platform and sizes of ocelli. Column 3: extension of the ocelli beyond the compound eyes.

##### Queen: Unknown

Specimens examined: INDONESIA: Forestry Center, Tabo Tabo, South Sulawesi, 3 VI 1989 (GWO), 5 workers, and 4 drones, 6 VI 1989 (GWO), 6 workers; Kamarora, Palolo Valley, Sigi Regency, Central Sulawesi, 9 IX 1989 (GWO), 5 workers; 9 XI 1995 (GWO), 1 worker; 22 IX 1996 (GWO), 1 worker; Rahmat, Palolo Valley, Sigi Regency, Central Sulawesi, 19 IX 1998 (GWO), 13 workers; Wuasa, Kab. Poso, Kech. Lore Utara, Central Sulawesi, 9 XI 1996 (GWO) 1 worker.

##### Distribution

Indonesia: Sulawesi and nearby islands (e.g., Buton Is., Kabaena Is., Taliabu Is.) of Indonesia (Smith, 2021; iNaturalist, 2024).

##### Remarks

This taxon of giant honey bee has a strongly raised ocellar triangle and the ocelli are relatively larger than in the other taxa in both workers and drones, possibly suggesting that it is more nocturnal in its foraging than other *Megapis* species Fig. 3, 6 and S2: A). The *indica* vein on the worker hind wing is shorter than in *laboriosa* but does not differ from *dorsata* or *breviligula*. On average, sternum 5 proportionately longer (i.e., ratio of length/width is greater) than that of *breviligula* worker, similar to report by Maa (1953) (Fig. 3.9).

## 3.1.2 *Apis breviligula* (Maa), 1953

*Megapis breviligula* Maa 1953: 563 [*breviligula* Maa]

*Apis dorsata breviligula* (Maa); Engel 1999: 186 [*breviligula* Maa]

Description of worker *A. breviligula* (Fig. 2: B-1)

a. Integument and Pubescence: Integument of adult worker black over entire body. Bronzy iridescence on the edge of the labrum, mandibles, and legs. Pattern of pubescence coloration not different from *binghami.* Callow individuals pale grey, as in *binghami*.
b. Structures:
  I. Head: Ocellar triangle raised (S2: A). Ocelli domed. Lateral ocelli angled toward compound eyes. Diameter of median ocellus, 0.451–0.483 mm. Interocellar distance (IOD) 0.380–0.479 mm; ocellocular distance (OOD), 0.268–0.338 mm; ratio of IOD/OOD, 1.193–1.520 (Fig. 3A). Malar length and width, 0.522–0.663 mm and 0.640–0.722 mm, respectively. Ratio of upper ocular distance and head width (UOD/HW), 0.377–0.408. Tongue, 5.933–6.456 mm long (Fig. 5A).
  II. Metasomal abdomen: Anteglandulus of sternum 2 angulated at the midpoint (Fig. 4: B-1). Glandulus on sterna 3–5 curved posteriorly at the midpoint (Fig. 4: B-2, showing for sternum 3). Ratio of length to width of sternum 5, 0.535–0.563. Posterior laminal spiracularis on 7^th^ hemitergite not invaginated. On sting, the number of pairs of barbs ranges from 10–11.
  III. Wing: *indica* vein on hind wing 0.327–0.617 mm long.

Description of drone *A. breviligula* (Fig. 2: B-2)

a. Integument: Integument of head, antennae, mesosoma, legs and metasoma iridescent brown.
b. Pubescence: Brown hairs on head, mesosoma and metasoma with pale yellow to brown hairs from ventral view. Mesosoma fully covered with long, pale yellow to brown hairs. Clumps of brown hairs on basitarsus III.
c. Structure: Ocellar triangle prominently raised. Ocelli protrude strongly, angled towards compound eyes (Fig. 6: B-1). Lateral ocelli seem to meet compound eyes with no space between them (Fig. 6: B-2). In dorsal view, ocelli extend beyond the plane of compound eyes (Fig. 6: B-3).

### Queen: Unknown

Specimens examined: PHILIPPINES: Bulano Pasan, Zamboanga, Mindanao, (YCS) 4 workers; Calinan, Davao City, Davao del Sur, Mindanao, 30 IX 1998 (YCS), 16 workers, 3 drones; College Villa, Los Baños, Luzon, 2008 (DRS), 2 workers.

### Distribution

Throughout the Philippine Islands, excluding Palawan (Smith, 2021).

### Remarks

Workers of *breviligula* are very similar to *binghami* in their overall morphological appearance (Fig. 2). Maa (1953) described *breviligula* as a species based in part on its short tongue length. However, in the fully extended tongues measured in the present study, this was not true: tongue lengths overlapped broadly and mean length did not differ statistically between *binghami* and *breviligula* (Fig. 5A). This study confirmed the relatively less prominently raised ocellar triangle of *breviligula* compared to *binghami* (Maa, 1953), and its proportionately shorter sternum 5 (Fig. 5B). Furthermore, the lateral ocelli bulge outwards toward compound eyes but to a lesser extent than in *binghami* (Fig. 3B; S2: A). Diameter of the median ocellus is similar to *binghami* but larger than in *dorsata* and *laboriosa* (Fig. 3A). Values for the ratio of upper ocular distance to head width (UOD/HW) are very similar between *breviligula* and *binghami* but smaller than those of *dorsata* and *laboriosa* (Fig. 3C). Malar length and width are approximately the same in *binghami* and *dorsata* (Fig. 3D), but shorter in length than in *laboriosa.* No distinct differences between *binghami* and *breviligula* drones were observed, however, they both have prominently raised ocelli that are larger than in *dorsata* and very unlike *laboriosa* that has relatively flat ocelli and ocellar triangle.

## 3.1.3 *Apis dorsata* Fabricius, 1793

*Apis dorsata* Fabricius 1793: 328 [*dorsata* Fabricius]

*Apis nigripennis* Latreille 1804: 170 [*dorsata* Fabricius]

*Apis bicolor* Klug 1807: 264. Preoccupied (*nec* Fabricius 1781, Villers 1789) [*dorsata* Fabricius]

*Apis testacea* Smith 1858: 49 [*dorsata* Fabricius]

*Apis testaccea* Smith 1871: 396 *Lapsus calami* [*dorsata* Fabricius]

*Megapis dorsata* (Smith); Ashmead 1904: 121 [*dorsata* Fabricius]

*Apis darsata* Baldensperger 1928: 173 *Lapsus calami* [*dorsata* Fabricius]

*Megapis dorsata* (Fabricius), Maa 1953: 566 [*dorsata* Fabricius]

*Apis dorsatao* Ruttner 1988: 118 *Lapsus calami* [*dorsata* Fabricius]

*Apis dorsata dorsata* (Fabricius), Engel 1999: 186 [*dorsata* Fabricius]

Description of worker *A. dorsata* (Fig. 2: C-1)

a. Integument: Integument of head and antennae shiny black except the apex of the mandibles and labrum that are reddish brown. Legs black, excluding the margins and ventral surface of basitarsi III that are reddish brown. Mesosoma integument black. Metasomal terga 1–3 and sterna 1–2 yellow to brown; other terga and sterna black to apex of metasoma (Fig. 2: C-1). Extent of bicolorism varies, with yellow integument extending to tergum 4 or 5 in some individuals. Young callow individuals with pale yellowish integument. Specimens collected from southern India and Sri Lanka not noticeably different from those of Southeast Asia.
b. Pubescence:

I. Head: Yellowish and brown hairs on labrum; short, decumbent white hairs on clypeus, paraocular area, and adjacent antennal suture; stiff, medium to long black and brown hairs on frons, ocellar triangle and vertex of head; pale yellow and brown hairs on compound eyes; short black hairs on antennal scape, pedicel and first two segments of flagellum, followed by short decumbent pale yellow hairs on the remaining flagellar segments; pale yellow hairs on genal area.
II. Mesosoma: Long black hairs cover pronotum and scutum; long brown to yellow hairs on scutellum, metanotum and propodeum that continue onto metasomal abdominal segments. Brown hairs on ventral mesosoma. Patch of brown hairs present at the mesopleuron. Long, dense, yellow hairs on coxae and trochanters; long, scattered, yellow hairs on femurs; and stiff brown hairs on basitarsi and tibiae.
III. Metasoma: Yellow hairs on terga 1–2 (T1–T2); mixture of pale yellow and black hairs laterally on T3 and T4; yellow hairs on the anterior margin of T5 and T6. Black hairs cover majority of T5 and T6 (Fig. 2: C-1). From ventral view, yellow to brown hairs on sterna 1 and 2. Transverse band of short, decumbent, whitish hairs on anterior bases of sterna 3–5. The extent of whitish hairs visible varies depends on how bees died and the extension of the metasoma of the dead bees, e.g. the whitish or yellow hairs seem to be absent or dull in specimens killed in ethanol.
c. Structures:

I. Head: Ocelli arise from a raised pedestal (S2: A). Lateral ocelli much bulged and tilted toward compound eyes. Median ocellus diameter, 0.399–0.439 mm (Fig. 3A). Interocellar distance (IOD) approximately equal to the distance between lateral ocelli and compound eyes (OOD) (IOD = 0.370– 0.510 mm; OOD = 0.380–510 mm). Ratio of IOD/OOD, 0.813–1.159 (Fig. 3B). Malar length (0.523–0.662 mm) equal to or slightly shorter than malar width (0.593–0.797 mm). Ratio of upper ocular distance (UOD) to head width (HW), 0.420–0.459 (Fig. 3C).
II. Metasomal abdomen: Anteglandulus medially angulated, not arcuate (Fig. 4: C-1). Glandulus at the midpoint of sterna 3–5 curved posteriorly. Absence of invaginated posterior laminal spiracularis on 7^th^ hemitergite. 10–11 pairs of barbs on sting.
III. Wing: *indica* vein on hind wing, 0.264–0.728 mm. *Apis dorsata* from across its range have similar morphological appearance. The South India and North India groups, as indicated by molecular data (Smith, 1991; 2021; Lo et al., 2010, Kitnya et al., 2022), are not differentiated by morphological characters. However, worker specimens from Andaman Islands are smaller in overall size (example, wing length and width, the traits that serve as proxies for body size) though coloration pattern and other morphological features do not distinguish them from mainland *dorsata*.

Description of drone, *A. dorsata* (Fig. 2: C-2)

a. Integument: Integument of head, scape and pedicel brown; flagellum orange brown. Mesosoma black. Metasomal terga 2 and 3 orange-brown (tergum 1 cannot be viewed, fully covered with dense hairs); posterior portion of terga 2 and 3 light brown. Terga 3–7 chestnut brown. Legs brown, with last tarsi of hind legs orange.
b. Pubescence: Brown hairs on face, eyes and gena; long, dense, brown hairs fully covering the mesosoma and continuing onto metasomal tergum 1 and sternum 1. Short brown hairs on terga 2 and 3; long, stiff, brown hairs on terga 4–7. Pale yellow hairs on sterna 2–7. Long, brown hairs on coxae, trochanters, and femurs; nearly bare tibiae; clumps of pale-yellow hairs on ventral basitarsi III.
c. Structures: Ocellar triangle visibly raised. Ocelli protruded, strongly domed, and tilted towards compound eyes (Fig. 6: C-1 & C-2); extend beyond compound eyes in dorsal view (Fig. 6: C-3). We were unable to examine drones from Andaman Island due to lack of specimens.

### Queen: Unknown

Specimens examined: CHINA: Mandian Village, Dragon fruit land, Xishuangbanna, Yunnan, 30 IV 2019 (QL), 3 workers; Xishuangbanna Botanical Garden-National Museum, Xishuangbanna, Yunnan, 1 V 2019 (QL), 6 workers, 10 drones; INDIA: Amtung, West Siang, Arunachal Pradesh, 21 IV 2018 (NK, MG), 6 workers; Bambooflat, South Andaman Island, Andaman Island, 19 I 2018 (MVP), 2 workers; Canara Bank Layout, near NCBS Campus, Bangalore, Karnataka, 18 IV 2021 (NK), 3 drones; Hainikal, Peren, Nagaland, 1 V 2018 (NK), 4 workers, 9 drones; ICTS Campus, Bangalore, Karnataka, 16 II 2018 (GGT), 2 workers; Kaying, West Siang, Arunachal Pradesh (NK), 3 workers; Modi, Siang, Arunachal Pradesh, 28 III 2019 (NK, MG, GWO), 2 workers; Nag Mandir, West Kameng, Arunachal Pradesh, 28 × 2017 (MVP), 2 workers; NCBS Campus, Bangalore, Karnataka (NK), 4 workers; Palace ground, Bangalore, Karnataka, 9 II 2014 (Dudde), 3 drones; Pangkang, Upper Siang, Arunachal Pradesh, 28 III 2019 (NK, MG, GWO), 22 workers; Punjab University Campus, Chandigarh, Punjab (AB), 3 workers; Rono Hills, Papum Pare, Arunachal Pradesh, 4 XII 2017 (NK), 1 worker; Sahakara Nagar, Bangalore, Karnataka, 6 II 2020 (NK), 2 drones; Thungjang, Tirap, Arunachal Pradesh, 12 XI 2017 (NK), 7 workers; Tumbin, West Siang, Arunachal Pradesh 16 V 2019 (NK), 8 workers, 7 drones; Tutnyu, Tirap, Arunachal Pradesh, 13 × 2017 (NK), 7 workers; THAILAND: Phuang subdist, Khao Khitchakut, Chanthaburi, 6 II 1999 (NW), 3 drones; VIETNAM: Kien Giang, Mekong Delta, Phu Quoc, 21 II 2018 (BR), 10 workers; Moung Te, Lai Chau, 17 V 2007 (PHT), 3 workers; U Minh, Ca Mau, 8 VIII 2001 (Thai and Se), 6 workers, 1 drone.

### Distribution

Afghanistan, Pakistan, India, Nepal, Bhutan, Bangladesh, China, Myanmar, Thailand, Cambodia, Laos, Vietnam, Malaysia, Singapore, Indonesia (excluding Sulawesi and neighboring islands), Philippines (Palawan only), and Sri Lanka (Smith 2021).

### Remarks

This is the most widely distributed taxon of the giant honey bees. In workers, the mesosoma has brown to black hairs except the posterior and lateral edges of the scutum with pale yellow hairs. The metasomal abdomen is bicolored in foragers (Fig. 2: C-1) and uniformly yellow in callow workers. The legs are paler than in the other *Megapis* taxa. The lateral marginal area of sterna 2–5 is smallest in *dorsata*. From measurements in the present study, *dorsata* has the smallest wax-plates, in contrast to Maa’s (1953) report that the wax-plates of *binghami* are the smallest. Ratio of upper ocular distance to head width is smaller than in *laboriosa* but larger than in *binghami* and *breviligula* (Fig. 3C). Also, median ocellus diameter is intermediate between *laboriosa* and *binghami/breviligula* (Fig. 3A).

## 3.1.4 *Apis laboriosa* Smith, 1871

*Apis laboriosa* Smith in Moore et al. 1871: 249 [*laboriosa* Smith] *Apis himalayana* Maa, 1944: 4 (*nomen nudum*) [*laboriosa* Smith] *Megapis laboriosa* (Smith), Maa 1953: 570 [*laboriosa* Smith]

*Apis labortiosa* Willis, Winston, and Honda, 1992: 169 *Lapsus calami* [*laboriosa* Smith]

*Apis dorsata laboriosa* (Smith), Engel 1999: 186 [*laboriosa* Smith]

Description of worker *A. laboriosa* (Fig. 2: D-1)

a. Integument: Uniformly black over entire body, except the apical edge of mandibles, labrum, antennae, and legs are reddish brown.
b. Pubescence:
  I. Head: Mixture of short decumbent pale yellow, brown and black hairs on mandibles, labrum, clypeus, paraocular area; slightly longer pale yellow hairs around antennal suture; yellow and brown hairs on frons; long, dense, brown and golden hairs cover the vertex; long, yellow, and brown hairs on compound eyes; short, decumbent, black hairs on scape, pedicel, and first two flagellomeres of antennae while short, decumbent pale yellow hairs cover the flagellum from segment 3 onward to tip of antennae; long, dense golden-yellow hairs on genal area.
  II. Mesosoma: Long, dense, golden yellow hairs on pronotum; long, dense, brown hairs cover center of scutum, while dense, long, golden yellow hairs cover rest of the mesosoma, a distinctive character of *A. laboriosa* (Fig. 2: D-1). Unlike other taxa of giant honey bees, patch of brown hairs at the center of mesopleuron absent. Long, golden-yellow hairs on coxae, trochanters and femurs; stiff, brown to black hairs completely cover tibiae and basitarsi.
  III. Metasoma: Dorsal view: golden-yellow hairs completely cover tergum 1 (T1) and continue to the basal margin of T2 (Fig. 2: D-1); short black hairs on terga 2–6; thin bands of white hairs along anterior bases of T1–T5 (some of the bands of white hairs may become removed during aging of the bee, collection, or storage; therefore, in some specimens bands are absent); no band of white hairs at anterior basal margin of T6. Ventral view: short, decumbent black hairs with strips of white hairs at anterior bases of sterna 3–5. However, the presence of white hairs on terga varies between bees; also, they are not clearly visible if the specimens were collected in ethanol or if the bee died with compressed metasoma.

c) Structures:

I. Head: Region where ocelli occur is flat or only slightly raised (S2: A). Lateral ocelli do not tilt toward compound eyes (S2: B). Median ocellus width, 0.338–0.391 mm (Fig. 3A). Distance between lateral ocelli and compound eyes distinctly wide as compared to other three taxa: OOD = 0.550–0.740 mm, interocellar distance: IOD = 0.300–0.470 mm (Table 2). Ratio of interocellar distance to ocellocular distance (IOD/OOD), 0.500–0.712 (Fig. 3B; Table 2). Malar area, distinctly long, ML = 0.753–0.957 mm (Fig. 3D; S2: C). Ratio of upper ocular distance (UOD) to head width (HW), 0.480–0.519 mm.
II. Metasomal abdomen: Anteglandulus of sternum 2 medially arcuate (Fig. 4: D-1: sternum 2). Lateral marginal area of sterna 2–5 widest as compared to other three taxa. On sterna 3–5, glandulus at midpoint curved towards base (anteriorly) (Fig. 4: D-2: sternum 3). Posterior laminal spiracularis distinctly invaginated (7^th^ hemitergite) (Fig. 4: D-3). 13–14 pairs of barbs on sting.
III. Wing: *indica* vein on hind wing relatively long, 0.534–1.282 mm.

Description of drone *A. laboriosa* (Fig. 2: D-2)

a. Integument: Integument of head, antennae, mesosoma, legs and metasomal abdomen are uniformly black.
b. Pubescence: Brown to black hairs on head; black hairs on mesosoma; golden yellow hairs at the posterior scutum and scutellum and lateral mesosoma that continue onto metasomal tergum 1. Black branched hairs on metasomal tergum 2–4; black, long, unbranched, and stiff hairs on terga 5–6. Long yellow hairs on coxae, trochanters and femurs. Stiff, black hairs on baristarsi. Clumps of coppery brown hairs on ventral basitarsi III.
c. Structures: Ocellar triangle nearly flat (e.g., only slightly raised). Space between lateral ocelli is distinctly wide compare to all other *Megapis* taxa (Fig. 6: D-1 & D-2). Ocelli not visible beyond the plane of compound eyes from dorsal view (Fig. 6: D-3).

### Queen: Unknown

Specimens examined: CHINA: Xinping County-Sa, Yuxi City, Yunnan, 29 IV 2019 (QL), 6 workers; Xinping County-Street Town, Yuxi City, Yunnan, 14 IV 2019 (QL), 5 workers, 6 drones; INDIA: Benrue, Peren, Nagaland, 6 IV 2018 (NK), 3 workers; Kaakad, Rudraprayag, Uttarakhand, 2 V 2019 (MVP), 3 workers; Kala Pahar, Tirap, Arunachal Pradesh, 12 × 2017 (NK), 1 worker; Kanchyoti, Uttarakhand, (MVP), 8 workers; Modi, Siang, Arunachal Pradesh, 28 III 2019 (NK, KM, GWO), 2 workers; Nag Mandir, West Kameng, Arunachal Pradesh, 28 × 2017 (NK), 7 workers; New Kaspi, West Kameng, Arunachal Pradesh, 7 IX 2017 (NK), 1 worker; Photin, West Kameng, Arunachal Pradesh, 4 IV 2019 (NK, KM, GWO), 1 worker; Tutnyu, Tirap, Arunachal Pradesh, 12 × 2017, 28 IV 2018 (NK), 10 workers, 4 drones; Yembo, East Siang, Arunachal Pradesh, 26 III 2019 (NK, KM, GWO), 2 workers; NEPAL: Kaski, Gandaki Province, 8 V 1984 (BAU), 1 drone; VIETNAM: Mộc Châu, Sơn La, VII 2001 (PHT), 4 workers, 2 drones; Sapa, Lao Cai, 12 IV 2019 (THP), 3 workers, 1 drone; Mộc Châu Sơn La (SL), Moung La, 30 XI 2019, 1 worker (THP)

### Distribution

Pakistan (Azad Jammu and Kashmir), India (Uttarakhand, Sikkim, West Bengal, and states comprising North East India), Nepal, Bhutan, southern China, Myanmar, northern Thailand, northern Laos and northern Vietnam (Kitnya et al., 2020; Smith, 2021; Otis et al., submitted).

### Remarks

Among *Megapis*, this taxon is the most distinctive. *A. laboriosa* workers can be easily distinguished from other taxa by their long, golden-yellow thoracic hair (Fig. 2: D-1), flat ocellar triangle, barely projecting ocelli (S2: A), much wider distance between lateral ocelli and compound eye (OOD), distinctly longer malar area (Fig. 3D), wider lateral marginal area on S2–5, larger wax-plates, uniquely arcuate midpoint of glandulus of sternum 2 (Fig. 4: D-1), distinctly anteriorly curved midpoint of glandulus of sternum 3, shape of the posterior end of the lamella on the 7^th^ hermitergite (Fig. 4: D-3), and greater number of barbs on the sting. Drone ocelli are distinctly flat and do not extend beyond the plane of compound eyes from dorsal view (Fig. 6:D-2 & D-3). The drone metatarsus is narrower and longer in *laboriosa* than that of the other three giant honey bee taxa (Fig. 7).

**Figure 7:**
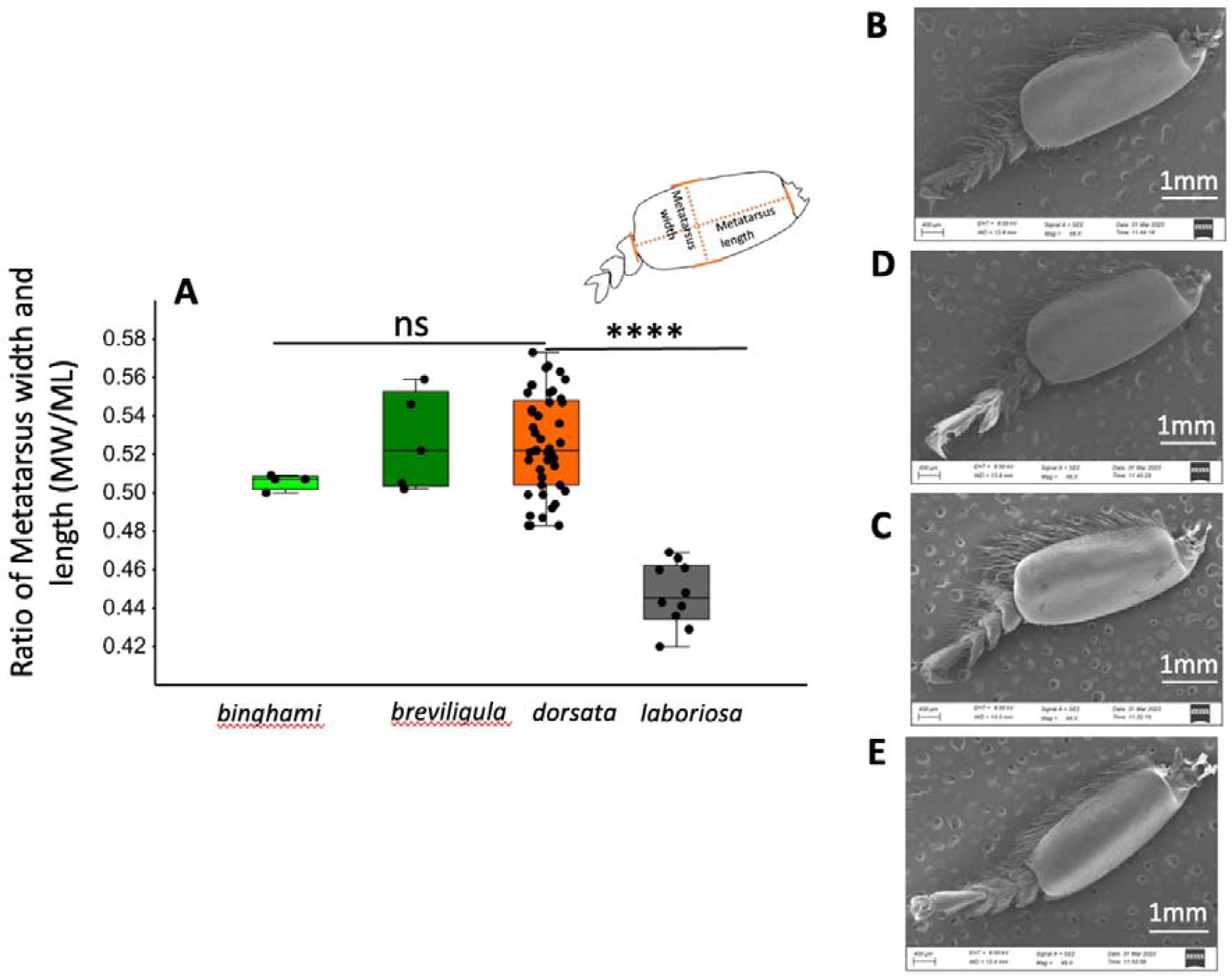
Leg morphology of *Megapis* drones. A. Ratio of metatarsus width to length. B–E: photographs of drone metatarsi (B. *binghami*; C. *breviligula*; D. *dorsata*; and E. *laboriosa*). Metatarsus of *laboriosa* is narrower.

After its original description by Smith in 1871, most honey bee researchers ignored the large, dark honey bees of Himalaya. In 1944, Maa referred to it as *Apis himalayana*. Later, in 1953, he described it as a distinct species, as confirmed by Sakagami et al. (1980). However, later researchers considered it to be a high elevation form of *A. dorsata* (Ruttner, 1988; Engel, 1999). Given the large number of autapomorphies of both drone and worker *A. laboriosa*, is remarkable that it was not widely accepted as a species long ago.

#### 3.2 Taxonomic Keys

## 3.2.1 Taxonomic key for *Megapis* workers

1 Ocelli slightly domed, positioned on nearly flat platform; IOD (<0.48 mm) shorter than OOD (>0.54 mm); median ocellus is <0.395 mm wide; UOD/HW is large (>0.47); malar space longer than broad, with ML >0.70 mm; glandulus of sternum 2 weekly curved at the mid and antero-lateral area; antegradular area of sternum 2 very wide; glandulus of sternum 3 curved anteriorly at the midpoint; posterior laminal spiracularis distinctly invaginated; 13–14 pairs of sting barbs; mesopleural patch at lateral mesosoma absent; dorsal mesosoma covered with long golden-yellow hairs.

… *A. laboriosa*

-Ocelli strongly domed, positioned on a prominent pedestal; IOD wider or equal to OOD; ratio of UOD/HW <0.47; malar length <0.70 mm and shorter or equal to malar width; median ocellus >0.395 mm wide; glandulus of sternum 2 sharply curved at the mid and antero-lateral area; antegradular area of sternum 2 not wide; glandulus at sternum 3 curved posteriorly at the midpoint; posterior laminal spiracularis distinctly not invaginated; 10–11 pairs of sting barbs; mesopleura patch of hairs at the lateral mesosoma present; black or brown hairs on dorsal mesosoma.

……………………. 2

2 IOD approximately equal to OOD; UOD/HW is >0.410; median ocellus <0.440 mm wide; integument of abdominal tergites is bicolored with anterior terga yellow or orange; lateral mesosoma hairs with distinct black patch at the centre.

… *A. dorsata*

-IOD distinctly greater than OOD; UOD/HW <0.410; median ocellus is >0.440 mm; integument of abdominal tergites entirely black; lateral mesosoma hairs with very small or no black patch at the centre.

… *A. binghami*

### 3.2.2 Taxonomic key for *Megapis* drones

1. Ocelli small and slightly convex, and enveloped by long hairs; lateral ocelli are widely separated, do not overlap compound eyes; the ocellar triangle is nearly flat; ocelli do not protrude beyond the plane of compound eyes from dorsal view; integument of head, mesosoma, metasoma, and legs black; scutellum and metasomal tergum 1 covered with long, golden-yellow hairs which continue to the anterior base of metasomal tergum 2; metasomal terga 2–4 covered with short black hairs; tergum 5 to apex of metasoma covered with very long, stiff, unbranched, black hairs.

… *A. laboriosa*

-Ocelli large, strongly convex (domed); lateral ocelli touch the compound eyes; the ocelli are positioned on a raised platform; the distance between lateral ocelli is narrow; the ocelli extend prominently beyond the plane of compound eyes from dorsal view; integument of head, antennae, mesosoma, metasoma range in color from orange to iridescent brown; scutellum and metasomal tergum 1 covered with long, pale yellow to brown hairs; metasomal terga 2–4 covered with short brown hairs; tergum 5 to apex of metasoma covered with very long, stiff, unbranched brown hairs.

..……………………. 2

2. Integument of metasomal terga 1–3 ranges from orange to light brown; metasomal tergum 4 onward with apical edge chestnut brown (bicolored metasoma); ocellar triangle visibly raised; callow drones have completely yellowish metasomal abdomen.

… *A. dorsata*

-Integument of all metasomal terga iridescent to dark brown (mono-colored); ocellar triangle prominently raised; callow drones have completely grey metasomal abdomen.

… *A. binghami*

## 4 Discussion

Our detailed examination of numerous morphological characters confirms the status of *A. laboriosa* as a distinct species, with several characters of both workers and drones separating it from the other taxa of giant honey bees. Several autapomorphies of *A. laboriosa* noted in the comparative study of *A. laboriosa* and *A. dorsata* from sites of co-occurrence (Kitnya et al., 2022) were consistent throughout their distributions. Proportions of head characters, median ocellus diameter, malar length, sternal glandulus shape, and spiracular plate (7^th^ hemitergite) shape are some of characters that allow one to easily distinguish *A. laboriosa* from the other taxa of giant honey bees, in addition to the very obvious differences in the color of thoracic hairs and abdominal tergites (for bees of mainland Asia) noted by Sakagami et al. (1980). The greater number of barbs on the lancets of the sting (13–14) is also diagnostic of *A. laboriosa*. Our results agree with and extend the previous reports of Maa (1953), Sakagami et al. (1980), and Trung et al. (1996).

The results also indicate that the widespread mainland *A. dorsata* is a species distinct from the giant honey bees of the islands of the oceanic Philippines and Sulawesi, Indonesia. Bicolored yellow/orange and black metasomal tergites, width of the median ocellus, and ratio of UOD/HW consistently separate *A. dorsata* from bees of those two island populations. Additionally, we detected no distinct morphological characters between the two genetic groups found in mainland India. This was surprising given the considerable genetic distance between the clades that inhabit southern portions of the Indian subcontinent and the rest of mainland Asia (Smith, 1991, 2021; Lo et al., 2010; Bhatta et al., in prep.). Additional sampling and analyses of both nuclear and mitochondrial DNA should resolve the extent of gene flow between the two groups and whether they represent cryptic species or simply divergent populations.

Worker bees from Andaman Island are generally smaller in size than mainland *A. dorsata*. For example, wing length and width, two traits which are proxies for overall body size, are notably smaller (mean and standard deviation of forewing length: mainland =13.11 mm; SD=0.098); Andaman Island = 12.08 mm; SD=0.073; mean and standard deviation of forewing width: mainland India =4.39 mm; SD=0.098; Andaman Island= 4.07 mm; SD=0.047). Worker bees from Andaman Island also have a significantly (p<0.0001) smaller median ocellus (Fig. 3A). However, the absence of non-size-related traits that differentiate these populations along with a lack of distinctive genetic differences (Bhatta et al., in prep.) suggest that the giant honey bees of the Andaman Islands are in the early stages of divergence from the population on the mainland. During the Last Glacial Maximum, 26.5–19.0 ka, the sea level was 120–130 m lower than it is today (Clark et al., 2009). At that time, the water gap between the Andaman Islands and mainland Asia was only ∼40–50 km wide, as inferred from the estimated shorelines when sea levels were lower (Voris, 2000; Ali, 2018). Swarms of bees may have flown between these land masses, allowing for gene exchange that would have slowed divergence. More detailed examination may lead to the discovery of morphological autapomorphies that distinguish these lineages.

We are the first to explore in detail the morphology of giant honey bees of the Philippines and Sulawesi, Indonesia. Maa indicated the giant honey bee of the Philippines, *Apis breviligula*—the “short-tongued honeybee”—is a species distinct from *binghami* of Sulawesi from which it “can be readily distinguished by its uniquely short glossa, longer malar areas, less strongly raised ocellar triangle and much shorter but broader abdominal sternum 5” (Maa, 1956, p. 564). In contrast to that statement, we have shown that tongue lengths do not differ significantly between these two geographic forms (Fig. 5A). The distributions of malar width/malar length of the two forms, although differing significantly, have overlapping distributions (Fig. 3D). Differences in the ocellar region and proportions of sternum 5 between *binghami* and *breviligula* are relatively minor and not diagnostic. These differences are similar in degree to differences between subspecies of *A. mellifera* and *A. cerana.* On the basis of morphological characters, we consider *breviligula* and *binghami* to be conspecific. As such, according to the rules of priority in taxonomic nomenclature, the giant honey bees of the Philippines would be recognized as a *Apis binghami breviligula* due to the earlier description of *A. binghami* from of Sulawesi.

Our morphological examination of *Megapis* bees firmly supports three species of giant honey bees: *A. laboriosa, A. dorsata,* and *A. binghami*. In contrast, phylogenic analyses of DNA sequences provide evidence for 4–5 species depending on the sequences examined: *A. laboriosa, A. dorsata, A. binghami*, *A. breviligula*, and the South Indian form (Arias and Sheppard 2005; Raffidiun and Crozier, 2007; Lo et al., 2010; Smith, 2021; Bhatta et al., in prep). A detailed comprehensive genetic analysis (e.g., based on ultra-conserved elements) along with more detailed morphological comparisons will help to resolve the phylogeny and taxonomy of *Megapis*.

## Data Availability

All the specimens examined are deposited in the Research Collection Facility, National Centre for Biological Sciences-TIFR, Bangalore. Specimen details are presented in the Supplementary Data (Suppl. File 1).

## Ethics Statement

Not applicable

## Authors’ Contribution

All authors conceptualized the project; NK examined the specimens and drafted the manuscript. All authors edited and approved the manuscript.

## Acknowledgements

The authors are thankful to the Research and Collection Facility, Imaging Facility of NCBS-TIFR, Bangalore, for the space and support to carry out the work. Authors also acknowledge various people who helped in collection or contributed the specimens used in the study:

Bharath Kumar (BK), Benjamin Rutschmann (BR); Chet Prasad Bhatta (CPB), Deborah R Smith (DRS), Geetha GT (GGT), Jaya Narah (JN), Karsing Megu (KM), Krishnaswamy A (KA), La Quang Trung (LQT), Natapot Warrit (NW), Pham Hong Thai (PHT), Pilot Dovih (PD), Prabhudev MV (PMV), Rajath S (RS), Savita Chib (SC), Stefan Reyes (SR), Tarun Karmakar (TK), Thomas Schmitt (TS), Viney Hedge (VH), YC Su (YCS), Yeshwant HM (YHM).

## SUPPLEMENTARY FILES

**Supplementary file1 (in excel spreadsheet)**

Details of specimens examined: collection locality, date, preservation method, collector, and voucher codes.

**Supplementary file2 (in powerpoint presentation)**

S2: A: Variation of ocellocular distance in *Megapis* worker. Arrow indicates the ocellocular distance (OOD) which is wider in *laboriosa* as compared to other three taxa: A. *binghami*; B. *breviligula*; C. *dorsata*; D. *laboriosa*.

S2: B: Variation in projection of compound eyes toward centre of head in *Megapis* workers. A. *binghami*; B. *breviligula*; C. *dorsata*; D. *laboriosa*. Arrow indicates the extension of compound eyes towards the center of the head. Note that in *binghami* and *breviligula*, the compound eyes seem to be largest (with the greatest extension inwards). The compound eyes of *laboriosa* project the least towards toward the centre of the head.

S2: C: Malar length in *Megapis* workers. A. *binghami*; B. *breviligula*; C. *dorsata*; D. *laboriosa*. The malar area is longer in *laboriosa* than in the other three taxa.

