## Supplementary file1 for "Taxonomic Revision and Identification Keys for the Giant Honey Bees": Supplimentary file1_Taxonomic Revision and Identification Keys for the Giant Honey Bees_submitted.pdf

Supplementary file 1: Taxonomic Revision and Identification Keys for the Giant Honey Bees_submitted

| Voucher code | Genu | Species | Caste | Collector | Collection year | month | day | Collection country | latitude (N, S) | - longitude (E, W) | Altitude (m) |
|---|---|---|---|---|---|---|---|---|---|---|---|
| NRC-AA-3038 | *Apis* | *binghami* | Female ( worker) | Gard W. Otis | 1989 | 6 | 3 | Indonesia | -4.734 | 119.68 | 420m |
| NRC-AA-3039 | *Apis* | *binghami* | Female ( worker) | Gard W. Otis | 1989 | 6 | 3 | Indonesia | -4.734 | 119.68 | 420m |
| NRC-AA-3040 | *Apis* | *binghami* | Female ( worker) | Gard W. Otis | 1989 | 6 | 3 | Indonesia | -4.734 | 119.68 | 420m |
| NRC-AA-3041 | *Apis* | *binghami* | Female ( worker) | Gard W. Otis | 1989 | 6 | 3 | Indonesia | -4.734 | 119.68 | 420m |
| NRC-AA-3042 | *Apis* | *binghami* | Female ( worker) | Gard W. Otis | 1989 | 6 | 3 | Indonesia | -4.734 | 119.68 | 420m |
| NRC-AA-3043 | *Apis* | *binghami* | Female ( worker) | Gard W. Otis | 1996 | 11 | 9 | Indonesia | -1.429 | 120.28 | 1100 m |
| NRC-AA-3044 | *Apis* | *binghami* | Female ( worker) | Gard W. Otis | 1989 | 6 | 6 | Indonesia | -4.736 | 119.68 | 420m |
| NRC-AA-3045 | *Apis* | *binghami* | Female ( worker) | Gard W. Otis | 1989 | 6 | 6 | Indonesia | -4.736 | 119.68 | 420m |
| NRC-AA-3046 | *Apis* | *binghami* | Female ( worker) | Gard W. Otis | 1989 | 6 | 6 | Indonesia | -4.736 | 119.68 | 420m |
| NRC-AA-3047 | *Apis* | *binghami* | Female ( worker) | Gard W. Otis | 1989 | 6 | 6 | Indonesia | -4.736 | 119.68 | 420m |
| NRC-AA-3048 | *Apis* | *binghami* | Female ( worker) | Gard W. Otis | 1995 | 11 | 9 | Indonesia | -1.212 | 120.15 | 600m |
| NRC-AA-3049 | *Apis* | *binghami* | Female ( worker) | Gard W. Otis | 1989 | 9 | 9 | Indonesia | -1.212 | 120.15 | 600m |
| NRC-AA-3050 | *Apis* | *binghami* | Female(worker) | Gard W. Otis | 1989 | 9 | 9 | Indonesia | -1.212 | 120.15 | 600m |
| NRC-AA-3051 | *Apis* | *binghami* | Female(worker) | Gard W. Otis | 1989 | 9 | 9 | Indonesia | -1.212 | 120.15 | 600m |
| NRC-AA-3052 | *Apis* | *binghami* | Female(worker) | Gard W. Otis | 1989 | 9 | 9 | Indonesia | -1.212 | 120.15 | 600m |
| NRC-AA-3053 | *Apis* | *binghami* | Female(worker) | Gard W. Otis | 1996 | 9 | 22 | Indonesia | -1.098 | 120.15 | 700m |
| NRC-AA-3054 | *Apis* | *binghami* | Female(worker) | Gard W. Otis | 1998 | 9 | 19 | Indonesia | -1.185 | 120.07 | 700m |
| NRC-AA-3055 | *Apis* | *binghami* | Female(worker) | Gard W. Otis | 1998 | 9 | 19 | Indonesia | -1.185 | 120.07 | 700m |
| NRC-AA-3056 | *Apis* | *binghami* | Female(worker) | Gard W. Otis | 1998 | 9 | 19 | Indonesia | -1.185 | 120.07 | 700m |
| NRC-AA-3057 | *Apis* | *binghami* | Female(worker) | Gard W. Otis | 1998 | 9 | 19 | Indonesia | -1.185 | 120.07 | 700m |
| NRC-AA-3058 | *Apis* | *binghami* | Female(worker) | Gard W. Otis | 1998 | 9 | 19 | Indonesia | -1.185 | 120.07 | 700m |
| NRC-AA-3059 | *Apis* | *binghami* | Female(worker) | Gard W. Otis | 1998 | 9 | 19 | Indonesia | -1.185 | 120.07 | 700m |
| NRC-AA-3060 | *Apis* | *binghami* | Female(worker) | Gard W. Otis | 1998 | 9 | 19 | Indonesia | -1.185 | 120.07 | 700m |
| NRC-AA-3061 | *Apis* | *binghami* | Female(worker) | Gard W. Otis | 1998 | 9 | 19 | Indonesia | -1.185 | 120.07 | 700m |
| NRC-AA-3062 | *Apis* | *binghami* | Female(worker) | Gard W. Otis | 1998 | 9 | 19 | Indonesia | -1.185 | 120.07 | 700m |
| NRC-AA-3063 | *Apis* | *binghami* | Female(worker) | Gard W. Otis | 1998 | 9 | 19 | Indonesia | -1.185 | 120.07 | 700m |
| NRC-AA-3064 | *Apis* | *binghami* | Female(worker) | Gard W. Otis | 1998 | 9 | 19 | Indonesia | -1.185 | 120.07 | 700m |
| NRC-AA-3065 | *Apis* | *binghami* | Female(worker) | Gard W. Otis | 1998 | 9 | 19 | Indonesia | -1.185 | 120.07 | 700m |
| NRC-AA-3066 | *Apis* | *binghami* | Female(worker) | Gard W. Otis | 1998 | 9 | 19 | Indonesia | -1.185 | 120.07 | 700m |
| NRC-AA-3067 | *Apis* | *binghami* | Male (drone) | Gard W. Otis | 1989 | 6 | 3 | Indonesia | -4.734 | 119.68 | 420m |
| NRC-AA-3068 | *Apis* | *binghami* | Male (drone) | Gard W. Otis | 1989 | 6 | 3 | Indonesia | -4.734 | 119.68 | 420m |
| NRC-AA-3069 | *Apis* | *binghami* | Male (drone) | Gard W. Otis | 1989 | 6 | 3 | Indonesia | -4.734 | 119.68 | 420m |
| NRC-AA-3070 | *Apis* | *binghami* | Male (drone) | Gard W. Otis | 1989 | 6 | 3 | Indonesia | -4.734 | 119.68 | 420m |
| NRC-AA-3084 | *Apis* | *binghami* | Female(worker) | Gard W. Otis | 1989 | 6 | 3 | Indonesia | -4.734 | 119.68 | 420m |
| NRC-AA-3085 | *Apis* | *binghami* | Female(worker) | Gard W. Otis | 1989 | 6 | 3 | Indonesia | -4.734 | 119.68 | 420m |
| NRC-AA-7097 | *Apis* | *binghami* | Female(worker) | Gard W. Otis | 1989 | 6 | 3 | Indonesia | -4.734 | 119.68 | 420m |
| NRC-AA-7098 | *Apis* | *binghami* | Female(worker) | Gard W. Otis | 1989 | 6 | 3 | Indonesia | -4.734 | 119.68 | 420m |
| NRC-AA-7099 | *Apis* | *binghami* | Female(worker) | Gard W. Otis | 1989 | 6 | 3 | Indonesia | -4.734 | 119.68 | 420m |
| NRC-AA-7100 | *Apis* | *binghami* | Female(worker) | Gard W. Otis | 1989 | 6 | 3 | Indonesia | -4.734 | 119.68 | 420m |
| NRC-AA-7101 | *Apis* | *binghami* | Female(worker) | Gard W. Otis | 1989 | 6 | 3 | Indonesia | -4.734 | 119.68 | 420m |
| NRC-AA-8080 | *Apis* | *binghami* | Female(worker) | Gard W Otis | 1989 | 6 | 3 | Indonesia | | | |
| NRC-AA-8081 | *Apis* | *binghami* | Female(worker) | Gard W Otis | 1989 | 6 | 3 | Indonesia | | | |
| NRC-AA-8082 | *Apis* | *binghami* | Female(worker) | Gard W Otis | 1989 | 6 | 3 | Indonesia | | | |
| NRC-AA-8083 | *Apis* | *binghami* | Female(worker) | Gard W Otis | 1989 | 6 | 3 | Indonesia | | | |
| NRC-AA-8084 | *Apis* | *binghami* | Female(worker) | Gard W Otis | 1989 | 6 | 3 | Indonesia | | | |
| NRC-AA-8085 | *Apis* | *binghami* | Female(worker) | Gard W Otis | 1989 | 6 | 3 | Indonesia | | | |
| NRC-AA-8086 | *Apis* | *binghami* | Female(worker) | Gard W Otis | 1989 | 6 | 3 | Indonesia | | | |
| NRC-AA-8087 | *Apis* | *binghami* | Female(worker) | Gard W Otis | 1989 | 6 | 3 | Indonesia | | | |
| NRC-AA-8088 | *Apis* | *binghami* | Female(worker) | Gard W Otis | 1989 | 6 | 3 | Indonesia | | | |
| NRC-AA-8089 | *Apis* | *binghami* | Female(worker) | Gard W Otis | 1989 | 6 | 3 | Indonesia | | | |
| NRC-AA-3020 | *Apis* | *binghami* | Female(worker) | Gard W. Otis | 1998 | 9 | 30 | Philippines | 7.14 | 125.39 | |
| NRC-AA-3021 | *Apis* | *binghami* | Female(worker) | Gard W. Otis | 1998 | 9 | 30 | Philippines | 7.14 | 125.39 | |

| | | | | | | | | | | |
|---|---|---|---|---|---|---|---|---|---|---|
| NRC-AA-3022 | *Apis binghami* | Female(worker) | Gard W. Otis | 1998 | 9 | 30 | Philippine | 7.14 | 125.39 | |
| NRC-AA-3023 | *Apis binghami* | Female(worker) | Gard W. Otis | 1998 | 9 | 30 | Philippine | 7.14 | 125.39 | |
| NRC-AA-3024 | *Apis binghami* | Female(worker) | Gard W. Otis | 1998 | 9 | 30 | Philippine | 7.14 | 125.39 | |
| NRC-AA-3025 | *Apis binghami* | Female(worker) | Gard W. Otis | 1998 | 9 | 30 | Philippine | 7.14 | 125.39 | |
| NRC-AA-3026 | *Apis binghami* | Female(worker) | Gard W. Otis | 1998 | 9 | 30 | Philippine | 7.14 | 125.39 | |
| NRC-AA-3027 | *Apis binghami* | Female(worker) | Gard W. Otis | 1998 | 9 | 30 | Philippine | 7.14 | 125.39 | |
| NRC-AA-3028 | *Apis binghami* | Female(worker) | Gard W. Otis | 1998 | 9 | 30 | Philippine | 7.14 | 125.39 | |
| NRC-AA-3029 | *Apis binghami* | Female(worker) | Gard W. Otis | 1998 | 9 | 30 | Philippine | 7.14 | 125.39 | |
| NRC-AA-3030 | *Apis binghami* | Female(worker) | Gard W. Otis | 1998 | 9 | 30 | Philippine | 7.14 | 125.39 | |
| NRC-AA-3031 | *Apis binghami* | Female(worker) | Gard W. Otis | 1998 | 9 | 30 | Philippine | 7.14 | 125.39 | |
| NRC-AA-3032 | *Apis binghami* | Female(worker) | Gard W. Otis | 1998 | 9 | 30 | Philippine | 7.14 | 125.39 | |
| NRC-AA-3033 | *Apis binghami* | Female(worker) | Gard W. Otis | 1998 | 9 | 30 | Philippine | 7.14 | 125.39 | |
| NRC-AA-3034 | *Apis binghami* | Female(worker) | Gard W. Otis | 1998 | 9 | 30 | Philippine | 7.14 | 125.39 | |
| NRC-AA-3035 | *Apis binghami* | Female(worker) | Gard W. Otis | 1998 | 9 | 30 | Philippine | 7.14 | 125.39 | |
| NRC-AA-3036 | *Apis binghami* | Female(worker) | Deborah Smith | 2008 | | | Philippine | 17.222 | 121.53 | |
| NRC-AA-3037 | *Apis binghami* | Female(worker) | Deborah Smith | 2008 | | | Philippine | 17.222 | 121.53 | |
| NRC-AA-3086 | *Apis binghami* | Male ( drone) | Gard W. Otis | 1998 | 9 | 30 | Philippine | 7.14 | 125.39 | |
| NRC-AA-7102 | *Apis binghami* | Female(worker) | Stefan Reyes | | | | Philippines | | | |
| NRC-AA-7103 | *Apis binghami* | Female(worker) | Stefan Reyes | | | | Philippines | | | |
| NRC-AA-7104 | *Apis binghami* | Female(worker) | Stefan Reyes | | | | Philippines | | | |
| NRC-AA-7105 | *Apis binghami* | Female(worker) | Stefan Reyes | | | | Philippines | | | |
| NRC-AA-7106 | *Apis binghami* | Female(worker) | Stefan Reyes | | | | Philippines | | | |
| NRC-AA-7107 | *Apis binghami* | Female(worker) | Stefan Reyes | | | | Philippines | | | |
| NRC-AA-7108 | *Apis binghami* | Female(worker) | Y.C Su | 2008 | 7 | 2 to 2 | Philippine | ° 1' 3.3" | ° 1' 44.5 | 781m |
| NRC-AA-7109 | *Apis binghami* | Female(worker) | Deborah Smith | | | | Philippines | | | |
| NRC-AA-7110 | *Apis binghami* | Female(worker) | Smith, and L V | 2004 | 3 | 4 | Philippine | 4°08′16″ | 0°51′19″E | |
| NRC-AA-7111 | *Apis binghami* | Female(worker) | Y.C Su | 2008 | 7 | 2 to 2 | Philippine | ° 1' 3.3" | ° 1' 44.5 | 781m |
| NRC-AA-7662 | *Apis binghami* | Female(worker) | Y.C Su | 2008 | 7 | 2 to 2 | Philippine | ° 1' 3.3" | ° 1' 44.5 | 781m |
| NRC-AA-7663 | *Apis binghami* | Male ( drone) | Smith, L Villafu | 2004 | 3 | 4 | Philippine | 4°08′16″ | 0°51′19″E | |
| NRC-AA-7664 | *Apis binghami* | Male ( drone) | Smith, L Villafu | 2004 | 3 | 4 | Philippine | 4°08′16″ | 0°51′19″E | |
| NRC-AA-7665 | *Apis binghami* | Male ( drone) | Smith, L Villafu | 2004 | 3 | 4 | Philippine | 4°08′16″ | 0°51′19″E | |
| NRC-AA-7666 | *Apis binghami* | Male ( drone) | Smith, L Villafu | 2004 | 3 | 4 | Philippine | 4°08′16″ | 0°51′19″E | |
| NRC-AA-7667 | *Apis binghami* | Male ( drone) | Smith, L Villafu | 2004 | 3 | 4 | Philippine | 4°08′16″ | 0°51′19″E | |
| NRC-AA-7668 | *Apis binghami* | Female(worker) | Y.C Su | 2008 | 7 | 2 to 2 | Philippine | ° 1' 3.3" | ° 1' 44.5 | 781m |
| NRC-AA-7669 | *Apis binghami* | Female(worker) | Y.C Su | 2008 | 7 | 2 to 2 | Philippine | ° 1' 3.3" | ° 1' 44.5 | 781m |
| NRC-AA-8090 | *Apis binghami* | Female(worker) | | | | | philippines | | | |
| NRC-AA-8091 | *Apis binghami* | Female(worker) | | | | | philippines | | | |
| NRC-AA-8092 | *Apis binghami* | Female(worker) | | | | | philippines | | | |
| NRC-AA-8093 | *Apis binghami* | Female(worker) | | | | | philippines | | | |
| NRC-AA-8094 | *Apis binghami* | Female(worker) | | | | | philippines | | | |
| NRC-AA-8095 | *Apis binghami* | Female(worker) | | | | | philippines | | | |
| NRC-AA-8096 | *Apis binghami* | Female(worker) | | | | | philippines | | | |
| NRC-AA-8097 | *Apis binghami* | Female(worker) | | | | | philippines | | | |
| NRC-AA-8098 | *Apis binghami* | Female(worker) | | | | | philippines | | | |
| NRC-AA-2922 | *Apis dorsata* | Female(worker) | Qiu Lifei | 2019 | 5 | 2 | China | 21.924 | 101.26 | |
| NRC-AA-2923 | *Apis dorsata* | Female(worker) | Qiu Lifei | 2019 | 5 | 2 | China | 21.924 | 101.26 | |
| NRC-AA-2924 | *Apis dorsata* | Female(worker) | Qiu Lifei | 2019 | 5 | 2 | China | 21.924 | 101.26 | |
| NRC-AA-2925 | *Apis dorsata* | Female(worker) | Qiu Lifei | 2019 | 5 | 2 | China | 21.924 | 101.26 | |
| NRC-AA-2926 | *Apis dorsata* | Female(worker) | Qiu Lifei | 2019 | 5 | 2 | China | 21.928 | 101.25 | |
| NRC-AA-2927 | *Apis dorsata* | Female(worker) | Zhou Xin | 2019 | 4 | 30 | China | 22.125 | 100.68 | |
| NRC-AA-2929 | *Apis dorsata* | Female(worker) | Zhou Xin | 2019 | 4 | 30 | China | 22.125 | 100.68 | |
| NRC-AA-2930 | *Apis dorsata* | Female(worker) | Nyaton Kitnya; | 2019 | 3 | 28 | India | 28.487 | 95.087 | 534m |
| NRC-AA-2931 | *Apis dorsata* | Female(worker) | Kitnya and Karsi | 2018 | 4 | 21 | India | 28.281 | 94.851 | 500m |
| NRC-AA-2932 | *Apis dorsata* | Female(worker) | Kitnya and Karsi | 2018 | 4 | 21 | India | 28.281 | 94.851 | 500m |
| NRC-AA-2933 | *Apis dorsata* | Female(worker) | Kitnya and Karsi | 2018 | 4 | 21 | India | 28.281 | 94.851 | 500m |
| NRC-AA-2934 | *Apis dorsata* | Female(worker) | Kitnya and Karsi | 2018 | 4 | 21 | India | 28.281 | 94.851 | 500m |
| NRC-AA-2935 | *Apis dorsata* | Female(worker) | Kitnya and Karsi | 2018 | 4 | 21 | India | 28.281 | 94.851 | 500m |
| NRC-AA-2936 | *Apis dorsata* | Female(worker) | Kitnya and Karsi | 2018 | 4 | 21 | India | 28.281 | 94.851 | 500m |
| NRC-AA-2937 | *Apis dorsata* | Female(worker) | Nyaton Kitnya | | | | India | 28.438 | 94.67 | 480m |
| NRC-AA-2938 | *Apis dorsata* | Female(worker) | Nyaton Kitnya | | | | India | 28.438 | 94.67 | 480m |
| NRC-AA-2939 | *Apis dorsata* | Female(worker) | Nyaton Kitnya | | | | India | 28.438 | 94.67 | 480m |
| NRC-AA-2940 | *Apis dorsata* | Female(worker) | Nyaton Kitnya | 2019 | 5 | 16 | India | 28.456 | 94.684 | 356m |
| NRC-AA-2941 | *Apis dorsata* | Female(worker) | Nyaton Kitnya | 2019 | 5 | 16 | India | 28.456 | 94.684 | 356m |
| NRC-AA-2942 | *Apis dorsata* | Female(worker) | Nyaton Kitnya | 2019 | 5 | 16 | India | 28.456 | 94.684 | 356m |

| | | | | | | | | | | | |
|---|---|---|---|---|---|---|---|---|---|---|---|
| NRC-AA-2943 | *Apis* | *dorsata* | Female(worker) | Nyaton Kitnya; | 2019 | 3 | 28 | India | 28.497 | 95.088 | 558m |
| NRC-AA-2944 | *Apis* | *dorsata* | Female(worker) | Nyaton Kitnya; | 2019 | 3 | 28 | India | 28.497 | 95.088 | 558m |
| NRC-AA-2945 | *Apis* | *dorsata* | Female(worker) | Nyaton Kitnya; | 2019 | 3 | 28 | India | 28.497 | 95.088 | 558m |
| NRC-AA-2946 | *Apis* | *dorsata* | Female(worker) | Nyaton Kitnya; | 2019 | 3 | 28 | India | 28.497 | 95.088 | 558m |
| NRC-AA-2947 | *Apis* | *dorsata* | Female(worker) | Nyaton Kitnya; | 2019 | 3 | 28 | India | 28.497 | 95.088 | 558m |
| NRC-AA-2948 | *Apis* | *dorsata* | Female(worker) | Nyaton Kitnya; | 2019 | 3 | 28 | India | 28.497 | 95.088 | 558m |
| NRC-AA-2949 | *Apis* | *dorsata* | Female(worker) | Nyaton Kitnya; | 2019 | 3 | 28 | India | 28.497 | 95.088 | 558m |
| NRC-AA-2950 | *Apis* | *dorsata* | Female(worker) | Nyaton Kitnya; | 2019 | 3 | 28 | India | 28.497 | 95.088 | 558m |
| NRC-AA-2951 | *Apis* | *dorsata* | Female(worker) | Nyaton Kitnya; | 2019 | 3 | 28 | India | 28.497 | 95.088 | 558m |
| NRC-AA-2952 | *Apis* | *dorsata* | Female(worker) | Nyaton Kitnya; | 2019 | 3 | 28 | India | 28.497 | 95.088 | 558m |
| NRC-AA-2953 | *Apis* | *dorsata* | Female(worker) | Nyaton Kitnya; | 2019 | 3 | 28 | India | 28.497 | 95.088 | 558m |
| NRC-AA-2954 | *Apis* | *dorsata* | Female(worker) | Nyaton Kitnya; | 2019 | 3 | 28 | India | 28.497 | 95.088 | 558m |
| NRC-AA-2955 | *Apis* | *dorsata* | Female(worker) | Nyaton Kitnya; | 2019 | 3 | 28 | India | 28.497 | 95.088 | 558m |
| NRC-AA-2956 | *Apis* | *dorsata* | Female(worker) | Nyaton Kitnya; | 2019 | 3 | 28 | India | 28.497 | 95.088 | 558m |
| NRC-AA-2957 | *Apis* | *dorsata* | Female(worker) | Nyaton Kitnya; | 2019 | 3 | 28 | India | 28.497 | 95.088 | 558m |
| NRC-AA-2959 | *Apis* | *dorsata* | Female(worker) | Nyaton Kitnya; | 2019 | 3 | 28 | India | 28.497 | 95.088 | 558m |
| NRC-AA-2960 | *Apis* | *dorsata* | Female(worker) | Nyaton Kitnya; | 2019 | 3 | 28 | India | 28.497 | 95.088 | 558m |
| NRC-AA-2961 | *Apis* | *dorsata* | Female(worker) | Axel Brockmann | | | | India | 30.359 | 76.45 | |
| NRC-AA-2962 | *Apis* | *dorsata* | Female(worker) | Axel Brockmann | | | | India | 30.359 | 76.45 | |
| NRC-AA-2963 | *Apis* | *dorsata* | Female(worker) | Axel Brockmann | | | | India | 30.359 | 76.45 | |
| NRC-AA-2964 | *Apis* | *dorsata* | Female(worker) | Qiu Lifei | 2019 | 5 | 2 | China | 21.928 | 101.25 | |
| NRC-AA-2965 | *Apis* | *dorsata* | Female(worker) | Nyaton Kitnya | | | | India | 13.072 | 77.58 | |
| NRC-AA-2966 | *Apis* | *dorsata* | Female(worker) | Nyaton Kitnya | 2018 | 5 | 1 | India | 25.643 | 93.753 | 460m |
| NRC-AA-2967 | *Apis* | *dorsata* | Female(worker) | Nyaton Kitnya | 2018 | 5 | 1 | India | 25.643 | 93.753 | 460m |
| NRC-AA-2968 | *Apis* | *dorsata* | Female(worker) | Nyaton Kitnya | 2018 | 5 | 1 | India | 25.643 | 93.753 | 460m |
| NRC-AA-2969 | *Apis* | *dorsata* | Female(worker) | Nyaton Kitnya | 2018 | 5 | 1 | India | 25.643 | 93.753 | 460m |
| NRC-AA-2970 | *Apis* | *dorsata* | Female(worker) | Thai & Se | 2001 | 8 | 8 | Vietnam | 9.377 | 104.98 | |
| NRC-AA-2971 | *Apis* | *dorsata* | Female(worker) | Thai & Se | 2001 | 8 | 8 | Vietnam | 9.377 | 104.98 | |
| NRC-AA-2972 | *Apis* | *dorsata* | Female(worker) | Thai & Se | 2001 | 8 | 8 | Vietnam | 9.377 | 104.98 | |
| NRC-AA-2973 | *Apis* | *dorsata* | Female(worker) | Thai & Se | 2001 | 8 | 8 | Vietnam | 9.377 | 104.98 | |
| NRC-AA-2974 | *Apis* | *dorsata* | Female(worker) | Thai & Se | 2001 | 8 | 8 | Vietnam | 9.377 | 104.98 | |
| NRC-AA-2975 | *Apis* | *dorsata* | Female(worker) | Thai & Se | 2001 | 8 | 8 | Vietnam | 9.377 | 104.98 | |
| NRC-AA-2976 | *Apis* | *dorsata* | Female(worker) | Pham Hong Tha | 2007 | 6 | 17 | Vietnam | 22.557 | 102.73 | |
| NRC-AA-2977 | *Apis* | *dorsata* | Female(worker) | Pham Hong Tha | 2007 | 6 | 17 | Vietnam | 22.557 | 102.73 | |
| NRC-AA-2978 | *Apis* | *dorsata* | Female(worker) | Pham Hong Tha | 2007 | 6 | 17 | Vietnam | 22.557 | 102.73 | |
| NRC-AA-2979 | *Apis* | *dorsata* | Female(worker) | Nyaton Kitnya; | 2019 | 3 | 28 | India | 28.497 | 95.088 | 558m |
| NRC-AA-2980 | *Apis* | *dorsata* | Female(worker) | Nyaton Kitnya; | 2019 | 3 | 28 | India | 28.497 | 95.088 | 558m |
| NRC-AA-2981 | *Apis* | *dorsata* | Female(worker) | Nyaton Kitnya; | 2019 | 3 | 28 | India | 28.497 | 95.088 | 558m |
| NRC-AA-2982 | *Apis* | *dorsata* | Female(worker) | Nyaton Kitnya; | 2019 | 3 | 28 | India | 28.497 | 95.088 | 558m |
| NRC-AA-2983 | *Apis* | *dorsata* | Male ( drone) | Qiu Lifei | 2019 | 5 | 1 | China | 21.924 | 101.26 | |
| NRC-AA-2984 | *Apis* | *dorsata* | Male ( drone) | Qiu Lifei | 2019 | 5 | 1 | China | 21.924 | 101.26 | |
| NRC-AA-2985 | *Apis* | *dorsata* | Male ( drone) | Qiu Lifei | 2019 | 5 | 1 | China | 21.924 | 101.26 | |
| NRC-AA-2986 | *Apis* | *dorsata* | Male ( drone) | Qiu Lifei | 2019 | 5 | 1 | China | 21.924 | 101.26 | |
| NRC-AA-2987 | *Apis* | *dorsata* | Male ( drone) | Qiu Lifei | 2019 | 5 | 1 | China | 21.924 | 101.26 | |
| NRC-AA-2988 | *Apis* | *dorsata* | Male ( drone) | Qiu Lifei | 2019 | 5 | 1 | China | 21.924 | 101.26 | |
| NRC-AA-2989 | *Apis* | *dorsata* | Male ( drone) | Qiu Lifei | 2019 | 5 | 1 | China | 21.924 | 101.26 | |
| NRC-AA-2990 | *Apis* | *dorsata* | Male ( drone) | Qiu Lifei | 2019 | 5 | 1 | China | 21.924 | 101.26 | |
| NRC-AA-2991 | *Apis* | *dorsata* | Male ( drone) | Qiu Lifei | 2019 | 5 | 1 | China | 21.924 | 101.26 | |
| NRC-AA-2992 | *Apis* | *dorsata* | Male ( drone) | Qiu Lifei | 2019 | 5 | 1 | China | 21.924 | 101.26 | |
| NRC-AA-2993 | *Apis* | *dorsata* | Male ( drone) | Nyaton Kitnya | 2019 | 5 | 16 | India | 28.456 | 94.684 | 356m |
| NRC-AA-2995 | *Apis* | *dorsata* | Male ( drone) | Nyaton Kitnya | 2018 | 5 | 1 | India | 25.643 | 93.753 | 532m |
| NRC-AA-2996 | *Apis* | *dorsata* | Male ( drone) | Nyaton Kitnya | 2018 | 5 | 1 | India | 25.643 | 93.753 | 532m |
| NRC-AA-2997 | *Apis* | *dorsata* | Male ( drone) | Nyaton Kitnya | 2018 | 5 | 1 | India | 25.643 | 93.753 | 532m |
| NRC-AA-2998 | *Apis* | *dorsata* | Male ( drone) | Nyaton Kitnya | 2018 | 5 | 1 | India | 25.643 | 93.753 | 532m |
| NRC-AA-2999 | *Apis* | *dorsata* | Male ( drone) | Nyaton Kitnya | 2018 | 5 | 1 | India | 25.643 | 93.753 | 532m |
| NRC-AA-3000 | *Apis* | *dorsata* | Male ( drone) | Dudde, USA | 2014 | 2 | 9 | India | 12.998 | 77.592 | |
| NRC-AA-3001 | *Apis* | *dorsata* | Male ( drone) | Dudde, USA | 2014 | 2 | 9 | India | 12.998 | 77.592 | |
| NRC-AA-3002 | *Apis* | *dorsata* | Male ( drone) | Dudde, USA | 2014 | 2 | 9 | India | 12.998 | 77.592 | |
| NRC-AA-3003 | *Apis* | *dorsata* | Male ( drone) | Nyaton Kitnya | 2020 | 2 | 6 | India | 13.065 | 77.59 | |
| NRC-AA-3004 | *Apis* | *dorsata* | Male ( drone) | Nyaton Kitnya | 2020 | 2 | 6 | India | 13.065 | 77.59 | |
| NRC-AA-3005 | *Apis* | *dorsata* | Male ( drone) | Nyaton Kitnya | 2021 | 4 | 18 | India | 13.07 | 77.575 | |

| | | | | | | | | | | | |
|---|---|---|---|---|---|---|---|---|---|---|---|
| NRC-AA-3006 | *Apis* | *dorsata* | Male ( drone) | Nyaton Kitnya | 2021 | 4 | 18 | India | 13.07 | 77.575 | |
| NRC-AA-3007 | *Apis* | *dorsata* | Male ( drone) | Nyaton Kitnya | 2021 | 4 | 18 | India | 13.07 | 77.575 | |
| NRC-AA-3009 | *Apis* | *dorsata* | Male ( drone) | Nyaton Kitnya | 2018 | 5 | 1 | India | 25.643 | 93.753 | 460m |
| NRC-AA-3010 | *Apis* | *dorsata* | Male ( drone) | Nyaton Kitnya | 2018 | 5 | 1 | India | 25.643 | 93.753 | 460m |
| NRC-AA-3011 | *Apis* | *dorsata* | Male ( drone) | Nyaton Kitnya | 2018 | 5 | 1 | India | 25.643 | 93.753 | 460m |
| NRC-AA-3012 | *Apis* | *dorsata* | Male ( drone) | Nyaton Kitnya | 2018 | 5 | 1 | India | 25.643 | 93.753 | 460m |
| NRC-AA-3013 | *Apis* | *dorsata* | Male ( drone) | Thai & Se | 2001 | 8 | 8 | Vietnam | 9.377 | 104.98 | |
| NRC-AA-3014 | *Apis* | *dorsata* | Male ( drone) | Nyaton Kitnya | 2019 | 5 | 16 | India | 28.456 | 94.684 | 356m |
| NRC-AA-3015 | *Apis* | *dorsata* | Male ( drone) | Nyaton Kitnya | 2019 | 5 | 16 | India | 28.456 | 94.684 | 356m |
| NRC-AA-3016 | *Apis* | *dorsata* | Male ( drone) | Nyaton Kitnya | 2019 | 5 | 16 | India | 28.456 | 94.684 | 356m |
| NRC-AA-3017 | *Apis* | *dorsata* | Male ( drone) | Nyaton Kitnya | 2019 | 5 | 16 | India | 28.456 | 94.684 | 356m |
| NRC-AA-3018 | *Apis* | *dorsata* | Male ( drone) | Nyaton Kitnya | 2019 | 5 | 16 | India | 28.456 | 94.684 | 356m |
| NRC-AA-3019 | *Apis* | *dorsata* | Male ( drone) | Nyaton Kitnya | 2019 | 5 | 16 | India | 28.456 | 94.684 | 356m |
| NRC-AA-3071 | *Apis* | *dorsata* | Female(worker) | Nyaton Kitnya | 2019 | 5 | 16 | India | 28.456 | 94.684 | 356m |
| NRC-AA-3072 | *Apis* | *dorsata* | Female(worker) | Nyaton Kitnya | 2019 | 5 | 16 | India | 28.456 | 94.684 | 356m |
| NRC-AA-3073 | *Apis* | *dorsata* | Female(worker) | Nyaton Kitnya | 2019 | 5 | 16 | India | 28.456 | 94.684 | 356m |
| NRC-AA-3074 | *Apis* | *dorsata* | Female(worker) | Nyaton Kitnya | 2019 | 5 | 16 | India | 28.456 | 94.684 | 356m |
| NRC-AA-3075 | *Apis* | *dorsata* | Female(worker) | Nyaton Kitnya | 2019 | 5 | 16 | India | 28.456 | 94.684 | 356m |
| NRC-AA-3076 | *Apis* | *dorsata* | Female(worker) | | 1999 | 2 | 6 | Thailand | | | |
| NRC-AA-3077 | *Apis* | *dorsata* | Female(worker) | | 1999 | 2 | 6 | Thailand | | | |
| NRC-AA-3078 | *Apis* | *dorsata* | Female(worker) | | 1999 | 2 | 6 | Thailand | | | |
| NRC-AA-3087 | *Apis* | *dorsata* | Female(worker) | Nyaton Kitnya; | 2019 | 3 | 28 | India | 28.497 | 95.088 | 558m |
| NRC-AA-3839 | *Apis* | *dorsata* | Female(worker) | Nyaton Kitnya | 2017 | 10 | 13 | India | 26.934 N | 95.576 E | 1470m |
| NRC-AA-3840 | *Apis* | *dorsata* | Female(worker) | Nyaton Kitnya | 2017 | 10 | 13 | India | 26.934 N | 95.576 E | 1470m |
| NRC-AA-3841 | *Apis* | *dorsata* | Female(worker) | Nyaton Kitnya | 2019 | 3 | 28 | India | 28.487 N | 95.087 E | 534m |
| NRC-AA-3842 | *Apis* | *dorsata* | Female(worker) | Nyaton Kitnya | 2019 | 3 | 28 | India | 28.487 N | 95.087 E | 534m |
| NRC-AA-3843 | *Apis* | *dorsata* | Female(worker) | Nyaton Kitnya | 2019 | 3 | 28 | India | 28.487 N | 95.087 E | 534m |
| NRC-AA-3844 | *Apis* | *dorsata* | Female(worker) | Nyaton Kitnya | 2019 | 3 | 28 | India | 28.487 N | 95.087 E | 534m |
| NRC-AA-3845 | *Apis* | *dorsata* | Female(worker) | Nyaton Kitnya | 2019 | 3 | 28 | India | 28.487 N | 95.087 E | 534m |
| NRC-AA-3846 | *Apis* | *dorsata* | Female(worker) | Nyaton Kitnya | 2017 | 10 | 28 | India | 27.206 N | 92.561 E | 1164m |
| NRC-AA-3847 | *Apis* | *dorsata* | Female(worker) | Nyaton Kitnya | 2017 | 10 | 28 | India | 27.206 N | 92.561 E | 1164m |
| NRC-AA-3848 | *Apis* | *dorsata* | Female(worker) | Nyaton Kitnya | 2017 | 10 | 28 | India | 27.206 N | 92.561 E | 1164m |
| NRC-AA-3849 | *Apis* | *dorsata* | Female(worker) | Nyaton Kitnya | 2017 | 10 | 28 | India | 27.206 N | 92.561 E | 1164m |
| NRC-AA-3850 | *Apis* | *dorsata* | Female(worker) | Nyaton Kitnya | 2017 | 10 | 28 | India | 27.206 N | 92.561 E | 1164m |
| NRC-AA-3851 | *Apis* | *dorsata* | Female(worker) | Nyaton Kitnya | 2017 | 10 | 28 | India | 27.206 N | 92.561 E | 1164m |
| NRC-AA-3852 | *Apis* | *dorsata* | Female(worker) | Nyaton Kitnya | 2017 | 10 | 28 | India | 27.206 N | 92.561 E | 1164m |
| NRC-AA-3853 | *Apis* | *dorsata* | Female(worker) | Nyaton Kitnya | 2019 | 5 | 16 | India | 28.456 N | 94.684 E | 356m |
| NRC-AA-3854 | *Apis* | *dorsata* | Female(worker) | Nyaton Kitnya | 2019 | 5 | 16 | India | 28.456 N | 94.684 E | 356m |
| NRC-AA-3855 | *Apis* | *dorsata* | Female(worker) | Nyaton Kitnya | 2019 | 5 | 16 | India | 28.456 N | 94.684 E | 356m |
| NRC-AA-3856 | *Apis* | *dorsata* | Female(worker) | Nyaton Kitnya | 2019 | 5 | 16 | India | 28.456 N | 94.684 E | 356m |
| NRC-AA-3857 | *Apis* | *dorsata* | Female(worker) | Nyaton Kitnya | 2019 | 5 | 16 | India | 28.456 N | 94.684 E | 356m |
| NRC-AA-3858 | *Apis* | *dorsata* | Female(worker) | Nyaton Kitnya | 2019 | 5 | 16 | India | 28.456 N | 94.684 E | 356m |
| NRC-AA-3859 | *Apis* | *dorsata* | Female(worker) | Nyaton Kitnya | 2019 | 5 | 16 | India | 28.456 N | 94.684 E | 356m |
| NRC-AA-3860 | *Apis* | *dorsata* | Female(worker) | Nyaton Kitnya | 2019 | 5 | 16 | India | 28.456 N | 94.684 E | 356m |
| NRC-AA-3861 | *Apis* | *dorsata* | Female(worker) | Nyaton Kitnya | 2019 | 5 | 16 | India | 28.456 N | 94.684 E | 356m |
| NRC-AA-3862 | *Apis* | *dorsata* | Female(worker) | Nyaton Kitnya | 2019 | 5 | 16 | India | 28.456 N | 94.684 E | 356m |
| NRC-AA-3863 | *Apis* | *dorsata* | Female(worker) | Nyaton Kitnya | 2018 | 4 | 28 | India | 26.962 N | 95.631 E | 1060m |
| NRC-AA-3864 | *Apis* | *dorsata* | Female(worker) | Nyaton Kitnya | 2018 | 4 | 28 | India | 26.962 N | 95.631 E | 1060m |
| NRC-AA-3865 | *Apis* | *dorsata* | Female(worker) | Nyaton Kitnya | 2018 | 4 | 28 | India | 26.962 N | 95.631 E | 1060m |
| NRC-AA-3866 | *Apis* | *dorsata* | Female(worker) | Nyaton Kitnya | 2018 | 4 | 28 | India | 26.962 N | 95.631 E | 1060m |
| NRC-AA-3867 | *Apis* | *dorsata* | Female(worker) | Nyaton Kitnya | 2018 | 4 | 28 | India | 26.962 N | 95.631 E | 1060m |
| NRC-AA-3868 | *Apis* | *dorsata* | Female(worker) | Nyaton Kitnya | 2018 | 4 | 28 | India | 26.962 N | 95.631 E | 1060m |
| NRC-AA-3869 | *Apis* | *dorsata* | Female(worker) | Nyaton Kitnya | 2018 | 4 | 28 | India | 26.962 N | 95.631 E | 1060m |
| NRC-AA-3870 | *Apis* | *dorsata* | Female(worker) | Nyaton Kitnya | 2018 | 4 | 28 | India | 26.962 N | 95.631 E | 1060m |
| NRC-AA-3871 | *Apis* | *dorsata* | Female(worker) | Nyaton Kitnya | 2018 | 4 | 28 | India | 26.962 N | 95.631 E | 1060m |
| NRC-AA-3872 | *Apis* | *dorsata* | Female(worker) | Nyaton Kitnya | 2018 | 4 | 28 | India | 26.962 N | 95.631 E | 1060m |
| NRC-AA-3873 | *Apis* | *dorsata* | Female(worker) | Nyaton Kitnya | 2019 | 5 | 18 | India | 28.020 N | 95.340 E | 152m |
| NRC-AA-3874 | *Apis* | *dorsata* | Female(worker) | Nyaton Kitnya | 2019 | 5 | 18 | India | 28.020 N | 95.340 E | 152m |
| NRC-AA-3875 | *Apis* | *dorsata* | Female(worker) | Nyaton Kitnya | 2019 | 5 | 18 | India | 28.020 N | 95.340 E | 152m |
| NRC-AA-3876 | *Apis* | *dorsata* | Female(worker) | Nyaton Kitnya | 2019 | 5 | 18 | India | 28.020 N | 95.340 E | 152m |
| NRC-AA-3877 | *Apis* | *dorsata* | Female(worker) | Nyaton Kitnya | 2019 | 5 | 18 | India | 28.020 N | 95.340 E | 152m |

| | | | | | | | | | | | |
|---|---|---|---|---|---|---|---|---|---|---|---|
| NRC-AA-3878 | *Apis* | *dorsata* | Female(worker) | Nyaton Kitnya | 2019 | 5 | 18 | India | 28.020 N | 95.340 E | 152m |
| NRC-AA-3879 | *Apis* | *dorsata* | Female(worker) | Nyaton Kitnya | 2019 | 5 | 18 | India | 28.020 N | 95.340 E | 152m |
| NRC-AA-3880 | *Apis* | *dorsata* | Female(worker) | Nyaton Kitnya | 2019 | 5 | 18 | India | 28.020 N | 95.340 E | 152m |
| NRC-AA-3881 | *Apis* | *dorsata* | Female(worker) | Nyaton Kitnya | 2019 | 5 | 18 | India | 28.020 N | 95.340 E | 152m |
| NRC-AA-3882 | *Apis* | *dorsata* | Female(worker) | Nyaton Kitnya | 2019 | 5 | 18 | India | 28.020 N | 95.340 E | 152m |
| NRC-AA-3883 | *Apis* | *dorsata* | Female(worker) | Nyaton Kitnya | 2019 | 5 | 18 | India | 28.020 N | 95.280 E | 245 m |
| NRC-AA-3884 | *Apis* | *dorsata* | Female(worker) | Nyaton Kitnya | 2019 | 5 | 18 | India | 28.020 N | 95.280 E | 245 m |
| NRC-AA-3885 | *Apis* | *dorsata* | Female(worker) | Nyaton Kitnya | 2019 | 5 | 18 | India | 28.020 N | 95.280 E | 245 m |
| NRC-AA-3886 | *Apis* | *dorsata* | Female(worker) | Nyaton Kitnya | 2019 | 5 | 18 | India | 28.020 N | 95.280 E | 245 m |
| NRC-AA-3887 | *Apis* | *dorsata* | Female(worker) | Nyaton Kitnya | 2019 | 5 | 18 | India | 28.020 N | 95.280 E | 245 m |
| NRC-AA-3888 | *Apis* | *dorsata* | Female(worker) | Nyaton Kitnya | 2019 | 5 | 18 | India | 28.020 N | 95.280 E | 245 m |
| NRC-AA-3889 | *Apis* | *dorsata* | Female(worker) | Nyaton Kitnya | 2019 | 5 | 18 | India | 28.020 N | 95.280 E | 245 m |
| NRC-AA-3890 | *Apis* | *dorsata* | Female(worker) | Nyaton Kitnya | 2019 | 5 | 18 | India | 28.020 N | 95.280 E | 245 m |
| NRC-AA-3891 | *Apis* | *dorsata* | Female(worker) | Nyaton Kitnya | 2019 | 5 | 18 | India | 28.020 N | 95.280 E | 245 m |
| NRC-AA-3892 | *Apis* | *dorsata* | Female(worker) | Nyaton Kitnya | 2019 | 5 | 18 | India | 28.020 N | 95.280 E | 245 m |
| NRC-AA-3893 | *Apis* | *dorsata* | Female(worker) | Nyaton Kitnya | 2018 | 2 | 27 | India | 13.150 N | 77.060 E | 887m |
| NRC-AA-3894 | *Apis* | *dorsata* | Female(worker) | Nyaton Kitnya | 2018 | 2 | 27 | India | 13.150 N | 77.060 E | 887m |
| NRC-AA-3895 | *Apis* | *dorsata* | Female(worker) | Nyaton Kitnya | 2018 | 2 | 27 | India | 13.150 N | 77.060 E | 887m |
| NRC-AA-3896 | *Apis* | *dorsata* | Female(worker) | Nyaton Kitnya | 2018 | 2 | 27 | India | 13.030 N | 77.520 E | 918m |
| NRC-AA-3897 | *Apis* | *dorsata* | Female(worker) | Nyaton Kitnya | 2018 | 2 | 27 | India | 13.030 N | 77.520 E | 918m |
| NRC-AA-3898 | *Apis* | *dorsata* | Female(worker) | Nyaton Kitnya | 2018 | 2 | 27 | India | 13.030 N | 77.520 E | 918m |
| NRC-AA-3899 | *Apis* | *dorsata* | Female(worker) | Nyaton Kitnya | 2018 | 1 | 13 | India | 13.07 N | 77.580 E | 930m |
| NRC-AA-3900 | *Apis* | *dorsata* | Female(worker) | Nyaton Kitnya | 2018 | 1 | 13 | India | 13.07 N | 77.580 E | 930m |
| NRC-AA-3901 | *Apis* | *dorsata* | Female(worker) | Nyaton Kitnya | 2018 | 1 | 13 | India | 13.07 N | 77.580 E | 930m |
| NRC-AA-3902 | *Apis* | *dorsata* | Female(worker) | Nyaton Kitnya | 2018 | 1 | 13 | India | 13.07 N | 77.580 E | 930m |
| NRC-AA-7073 | *Apis* | *dorsata* | Female(worker) | itnya, Anne-Sop | 2023 | 6 | 14 | India | º"15.822" | 35'12.44 | 49m |
| NRC-AA-7112 | *Apis* | *dorsata* | Female(worker) | Sivaraju C | 2023 | 4 | 17 | India | 13.028 N | 77.585 E | |
| NRC-AA-7113 | *Apis* | *dorsata* | Female(worker) | Sivaraju C | 2023 | 4 | 17 | India | 13.028 N | 77.585 E | |
| NRC-AA-7114 | *Apis* | *dorsata* | Female(worker) | Sivaraju C | 2023 | 4 | 17 | India | 13.028 N | 77.585 E | |
| NRC-AA-7115 | *Apis* | *dorsata* | Female(worker) | Sivaraju C | 2023 | 4 | 17 | India | 13.028 N | 77.585 E | |
| NRC-AA-7116 | *Apis* | *dorsata* | Female(worker) | Sivaraju C | 2023 | 4 | 17 | India | 13.028 N | 77.585 E | |
| NRC-AA-7117 | *Apis* | *dorsata* | Female(worker) | Sivaraju C | 2023 | 4 | 17 | India | 13.028 N | 77.585 E | |
| NRC-AA-7118 | *Apis* | *dorsata* | Female(worker) | Sivaraju C | 2023 | 4 | 17 | India | 13.028 N | 77.585 E | |
| NRC-AA-7119 | *Apis* | *dorsata* | Female(worker) | Sivaraju C | 2023 | 4 | 17 | India | 13.028 N | 77.585 E | |
| NRC-AA-7120 | *Apis* | *dorsata* | Female(worker) | Sivaraju C | 2023 | 4 | 17 | India | 13.028 N | 77.585 E | |
| NRC-AA-7121 | *Apis* | *dorsata* | Female(worker) | Sivaraju C | 2023 | 4 | 17 | India | 13.028 N | 77.585 E | |
| NRC-AA-7122 | *Apis* | *dorsata* | Female(worker) | Sivaraju C | 2023 | 4 | 17 | India | 13.028 N | 77.585 E | |
| NRC-AA-7123 | *Apis* | *dorsata* | Female(worker) | Sivaraju C | 2023 | 4 | 17 | India | 13.028 N | 77.585 E | |
| NRC-AA-7124 | *Apis* | *dorsata* | Female(worker) | Sivaraju C | 2023 | 4 | 17 | India | 13.028 N | 77.585 E | |
| NRC-AA-7125 | *Apis* | *dorsata* | Female(worker) | Sivaraju C | 2023 | 4 | 17 | India | 13.028 N | 77.585 E | |
| NRC-AA-7126 | *Apis* | *dorsata* | Female(worker) | Sivaraju C | 2023 | 4 | 17 | India | 13.028 N | 77.585 E | |
| NRC-AA-7127 | *Apis* | *dorsata* | Female(worker) | Sivaraju C | 2023 | 4 | 17 | India | 13.028 N | 77.585 E | |
| NRC-AA-7128 | *Apis* | *dorsata* | Female(worker) | Sivaraju C | 2023 | 4 | 17 | India | 13.028 N | 77.585 E | |
| NRC-AA-7129 | *Apis* | *dorsata* | Female(worker) | Sivaraju C | 2023 | 4 | 17 | India | 13.028 N | 77.585 E | |
| NRC-AA-7130 | *Apis* | *dorsata* | Female(worker) | Sivaraju C | 2023 | 4 | 17 | India | 13.028 N | 77.585 E | |
| NRC-AA-7131 | *Apis* | *dorsata* | Female(worker) | Sivaraju C | 2023 | 4 | 17 | India | 13.028 N | 77.585 E | |
| NRC-AA-7132 | *Apis* | *dorsata* | Female(worker) | Krishnaswamy A | 2023 | 3 | 5 | India | 11.034 | 76.874 | |
| NRC-AA-7133 | *Apis* | *dorsata* | Female(worker) | Krishnaswamy A | 2023 | 3 | 5 | India | 11.034 | 76.874 | |
| NRC-AA-7134 | *Apis* | *dorsata* | Female(worker) | Krishnaswamy A | 2023 | 3 | 5 | India | 11.034 | 76.874 | |
| NRC-AA-7135 | *Apis* | *dorsata* | Female(worker) | Krishnaswamy A | 2023 | 3 | 5 | India | 11.034 | 76.874 | |
| NRC-AA-7136 | *Apis* | *dorsata* | Female(worker) | Krishnaswamy A | 2023 | 3 | 5 | India | 11.034 | 76.874 | |
| NRC-AA-7137 | *Apis* | *dorsata* | Female(worker) | Krishnaswamy A | 2023 | 3 | 5 | India | 11.034 | 76.874 | |
| NRC-AA-7138 | *Apis* | *dorsata* | Female(worker) | Krishnaswamy A | 2023 | 3 | 5 | India | 11.034 | 76.874 | |
| NRC-AA-7139 | *Apis* | *dorsata* | Female(worker) | Krishnaswamy A | 2023 | 3 | 5 | India | 11.034 | 76.874 | |
| NRC-AA-7140 | *Apis* | *dorsata* | Female(worker) | Krishnaswamy A | 2023 | 3 | 5 | India | 11.034 | 76.874 | |
| NRC-AA-7141 | *Apis* | *dorsata* | Female(worker) | Krishnaswamy A | 2023 | 3 | 5 | India | 11.034 | 76.874 | |

| | | | | | | | | | | | |
|---|---|---|---|---|---|---|---|---|---|---|---|
| NRC-AA-7142 | *Apis* | *dorsata* | Female(worker) | Savita Chib | 2023 | 3 | 15 | India | 22.461 N | 78.419 E | 980m |
| NRC-AA-7143 | *Apis* | *dorsata* | Female(worker) | Savita Chib | 2023 | 3 | 15 | India | 22.461 N | 78.419 E | 980m |
| NRC-AA-7144 | *Apis* | *dorsata* | Female(worker) | Savita Chib | 2023 | 3 | 15 | India | 22.461 N | 78.419 E | 980m |
| NRC-AA-7145 | *Apis* | *dorsata* | Female(worker) | Savita Chib | 2023 | 3 | 15 | India | 22.461 N | 78.419 E | 980m |
| NRC-AA-7146 | *Apis* | *dorsata* | Male (drone) | Nyaton Kitnya | 2023 | 2 | 28 | India | 13.094 N | 77.578 E | |
| NRC-AA-7147 | *Apis* | *dorsata* | Male (drone) | Nyaton Kitnya | 2023 | 2 | 28 | India | 13.094 N | 77.578 E | |
| NRC-AA-7148 | *Apis* | *dorsata* | Male (drone) | Nyaton Kitnya | 2023 | 2 | 28 | India | 13.094 N | 77.578 E | |
| NRC-AA-7149 | *Apis* | *dorsata* | Male (drone) | Nyaton Kitnya | 2023 | 2 | 28 | India | 13.094 N | 77.578 E | |
| NRC-AA-7150 | *Apis* | *dorsata* | Male (drone) | Nyaton Kitnya | 2023 | 2 | 28 | India | 13.094 N | 77.578 E | |
| NRC-AA-7151 | *Apis* | *dorsata* | Male (drone) | Nyaton Kitnya | 2023 | 2 | 28 | India | 13.094 N | 77.578 E | |
| NRC-AA-7152 | *Apis* | *dorsata* | Male (drone) | Nyaton Kitnya | 2023 | 2 | 28 | India | 13.094 N | 77.578 E | |
| NRC-AA-7153 | *Apis* | *dorsata* | Female(worker) | Savita Chib | 2023 | 3 | 15 | India | 22.461 N | 78.419 E | 980m |
| NRC-AA-7154 | *Apis* | *dorsata* | Female(worker) | Savita Chib | 2023 | 3 | 15 | India | 22.461 N | 78.419 E | 980m |
| NRC-AA-7155 | *Apis* | *dorsata* | Female(worker) | Savita Chib | 2023 | 3 | 15 | India | 22.461 N | 78.419 E | 980m |
| NRC-AA-7156 | *Apis* | *dorsata* | Female(worker) | Savita Chib | 2023 | 3 | 15 | India | 22.461 N | 78.419 E | 980m |
| NRC-AA-7157 | *Apis* | *dorsata* | Female(worker) | Savita Chib | 2023 | 3 | 15 | India | 22.461 N | 78.419 E | 980m |
| NRC-AA-7158 | *Apis* | *dorsata* | Female(worker) | Krishnaswamy A | 2023 | 3 | 5 | India | 11.034 | 76.874 | |
| NRC-AA-7159 | *Apis* | *dorsata* | Female(worker) | Krishnaswamy A | 2023 | 3 | 5 | India | 11.034 | 76.874 | |
| NRC-AA-7160 | *Apis* | *dorsata* | Female(worker) | Krishnaswamy A | 2023 | 3 | 5 | India | 11.034 | 76.874 | |
| NRC-AA-7161 | *Apis* | *dorsata* | Female(worker) | Krishnaswamy A | 2023 | 3 | 5 | India | 11.034 | 76.874 | |
| NRC-AA-7167 | *Apis* | *dorsata* | Female(worker) | Prabhudev MV | 2018 | 1 | 19 | India | 1°40.599' | 2°43.785 | 1060m |
| NRC-AA-7168 | *Apis* | *dorsata* | Female(worker) | Prabhudev MV | 2018 | 1 | 19 | India | 11.67 | 92.739 | 7m |
| NRC-AA-7169 | *Apis* | *dorsata* | Female(worker) | Prabhudev MV | 2018 | 1 | 19 | India | 1°40.599' | 2°43.785 | 152m |
| NRC-AA-7170 | *Apis* | *dorsata* | Female(worker) | Prabhudev MV | 2018 | 1 | 19 | India | 11.67 | 92.739 | 7m |
| NRC-AA-7171 | *Apis* | *dorsata* | Female(worker) | Nyaton Kitnya | 2023 | 2 | 14 | India | 13.07 N | 77.580 E | 930m |
| NRC-AA-7172 | *Apis* | *dorsata* | Female(worker) | Nyaton Kitnya | 2023 | 2 | 14 | India | 13.07 N | 77.580 E | 930m |
| NRC-AA-7173 | *Apis* | *dorsata* | Female(worker) | Nyaton Kitnya | 2023 | 2 | 14 | India | 13.07 N | 77.580 E | 930m |
| NRC-AA-7174 | *Apis* | *dorsata* | Female(worker) | Nyaton Kitnya | 2023 | 2 | 14 | India | 13.07 N | 77.580 E | 930m |
| NRC-AA-7175 | *Apis* | *dorsata* | Female(worker) | Nyaton Kitnya | 2023 | 2 | 14 | India | 13.07 N | 77.580 E | 930m |
| NRC-AA-7176 | *Apis* | *dorsata* | Female(worker) | Nyaton Kitnya | 2023 | 2 | 14 | India | 13.07 N | 77.580 E | 930m |
| NRC-AA-7177 | *Apis* | *dorsata* | Female(worker) | Nyaton Kitnya | 2023 | 2 | 14 | India | 13.07 N | 77.580 E | 930m |
| NRC-AA-7178 | *Apis* | *dorsata* | Female(worker) | Nyaton Kitnya | 2023 | 2 | 14 | India | 13.07 N | 77.580 E | 930m |
| NRC-AA-7179 | *Apis* | *dorsata* | Female(worker) | Nyaton Kitnya | 2023 | 2 | 14 | India | 13.07 N | 77.580 E | 930m |
| NRC-AA-7180 | *Apis* | *dorsata* | Female(worker) | Nyaton Kitnya | 2023 | 2 | 14 | India | 13.07 N | 77.580 E | 930m |
| NRC-AA-7181 | *Apis* | *dorsata* | Female(worker) | Nyaton Kitnya | 2023 | 2 | 14 | India | 13.07 N | 77.580 E | 930m |
| NRC-AA-7182 | *Apis* | *dorsata* | Female(worker) | Nyaton Kitnya | 2023 | 2 | 14 | India | 13.07 N | 77.580 E | 930m |
| NRC-AA-7183 | *Apis* | *dorsata* | Female(worker) | Nyaton Kitnya | 2023 | 2 | 14 | India | 13.07 N | 77.580 E | 930m |
| NRC-AA-7184 | *Apis* | *dorsata* | Female(worker) | Kumar K and F | 2023 | 2 | 15 | India | 13.07 | 77.575 | 930m |
| NRC-AA-7185 | *Apis* | *dorsata* | Female(worker) | Kumar K and F | 2023 | 2 | 15 | India | 13.07 | 77.575 | 930m |
| NRC-AA-7186 | *Apis* | *dorsata* | Female(worker) | Kumar K and F | 2023 | 2 | 15 | India | 13.07 | 77.575 | 930m |
| NRC-AA-7187 | *Apis* | *dorsata* | Female(worker) | Kumar K and F | 2023 | 2 | 15 | India | 13.07 | 77.575 | 930m |
| NRC-AA-7188 | *Apis* | *dorsata* | Female(worker) | Kumar K and F | 2023 | 2 | 15 | India | 13.07 | 77.575 | 930m |
| NRC-AA-7189 | *Apis* | *dorsata* | Female(worker) | Kumar K and F | 2023 | 2 | 15 | India | 13.07 | 77.575 | 930m |
| NRC-AA-7190 | *Apis* | *dorsata* | Female(worker) | Kumar K and F | 2023 | 2 | 15 | India | 13.07 | 77.575 | 930m |
| NRC-AA-7191 | *Apis* | *dorsata* | Female(worker) | Kumar K and F | 2023 | 2 | 15 | India | 13.07 | 77.575 | 930m |
| NRC-AA-7192 | *Apis* | *dorsata* | Female(worker) | Kumar K and F | 2023 | 2 | 15 | India | 13.07 | 77.575 | 930m |
| NRC-AA-7193 | *Apis* | *dorsata* | Female(worker) | Kumar K and F | 2023 | 2 | 15 | India | 13.07 | 77.575 | 930m |
| NRC-AA-7194 | *Apis* | *dorsata* | Female(worker) | Kumar K and F | 2023 | 2 | 15 | India | 13.07 | 77.575 | 930m |
| NRC-AA-7195 | *Apis* | *dorsata* | Female(worker) | Kumar K and F | 2023 | 2 | 15 | India | 13.07 | 77.575 | 930m |
| NRC-AA-7196 | *Apis* | *dorsata* | Female(worker) | Jaya Narah | 2018 | 4 | 24 | India | 24°48'46 | 93°56'08 | 775m |
| NRC-AA-7197 | *Apis* | *dorsata* | Female(worker) | Jaya Narah | 2018 | 4 | 24 | India | 24°48'46 | 93°56'08 | 775m |
| NRC-AA-7198 | *Apis* | *dorsata* | Female(worker) | Nyaton Kitnya | 2018 | 3 | 5 | India | 25.550 N | 93.445 E | 118 |

| | | | | | | | | | | | |
|---|---|---|---|---|---|---|---|---|---|---|---|
| NRC-AA-7199 | *Apis* | *dorsata* | Female(worker) | Nyaton Kitnya | 2018 | 3 | 5 | India | 25.550 N | 93.445 E | 118 |
| NRC-AA-7200 | *Apis* | *dorsata* | Female(worker) | Nyaton Kitnya | 2018 | 3 | 5 | India | 25.550 N | 93.445 E | 118 |
| NRC-AA-7201 | *Apis* | *dorsata* | Female(worker) | Nyaton Kitnya | 2018 | 3 | 5 | India | 25.550 N | 93.445 E | 118 |
| NRC-AA-7202 | *Apis* | *dorsata* | Female(worker) | Nyaton Kitnya | 2018 | 3 | 5 | India | 25.550 N | 93.445 E | 118 |
| NRC-AA-7203 | *Apis* | *dorsata* | Female(worker) | Nyaton Kitnya | 2018 | 3 | 5 | India | 25.550 N | 93.445 E | 118 |
| NRC-AA-7204 | *Apis* | *dorsata* | Female(worker) | Nyaton Kitnya | 2018 | 3 | 5 | India | 25.550 N | 93.445 E | 118 |
| NRC-AA-7205 | *Apis* | *dorsata* | Female(worker) | Nyaton Kitnya | 2019 | 6 | 15 | India | 25.404 N | 93.464 E | 496 |
| NRC-AA-7206 | *Apis* | *dorsata* | Female(worker) | Nyaton Kitnya | 2019 | 6 | 15 | India | 25.404 N | 93.464 E | 496 |
| NRC-AA-7207 | *Apis* | *dorsata* | Female(worker) | Nyaton Kitnya | 2019 | 6 | 15 | India | 25.404 N | 93.464 E | 496 |
| NRC-AA-7208 | *Apis* | *dorsata* | Female(worker) | Nyaton Kitnya | 2019 | 6 | 15 | India | 25.404 N | 93.464 E | 496 |
| NRC-AA-7209 | *Apis* | *dorsata* | Female(worker) | Nyaton Kitnya | 2018 | 5 | 1 | India | 25.643 | 93.753 | 460m |
| NRC-AA-7210 | *Apis* | *dorsata* | Female(worker) | Nyaton Kitnya | 2018 | 5 | 1 | India | 25.643 | 93.753 | 460m |
| NRC-AA-7211 | *Apis* | *dorsata* | Female(worker) | Nyaton Kitnya | 2018 | 5 | 1 | India | 25.643 | 93.753 | 460m |
| NRC-AA-7212 | *Apis* | *dorsata* | Female(worker) | Nyaton Kitnya | 2018 | 5 | 1 | India | 25.643 | 93.753 | 460m |
| NRC-AA-7213 | *Apis* | *dorsata* | Female(worker) | Nyaton Kitnya | 2018 | 5 | 1 | India | 25.643 | 93.753 | 460m |
| NRC-AA-7214 | *Apis* | *dorsata* | Female(worker) | Nyaton Kitnya | 2018 | 5 | 1 | India | 25.643 | 93.753 | 460m |
| NRC-AA-7215 | *Apis* | *dorsata* | Female(worker) | Nyaton Kitnya | 2018 | 5 | 1 | India | 25.643 | 93.753 | 460m |
| NRC-AA-7216 | *Apis* | *dorsata* | Female(worker) | Nyaton Kitnya | 2019 | 5 | 18 | India | 28.020 N | 95.340 E | 152m |
| NRC-AA-7217 | *Apis* | *dorsata* | Female(worker) | Nyaton Kitnya | 2019 | 5 | 18 | India | 28.020 N | 95.280 E | 245 m |
| NRC-AA-7218 | *Apis* | *dorsata* | Female(worker) | Nyaton Kitnya | 2019 | 5 | 18 | India | 28.020 N | 95.340 E | 152m |
| NRC-AA-7219 | *Apis* | *dorsata* | Female(worker) | Nyaton Kitnya | 2019 | 5 | 18 | India | 28.020 N | 95.340 E | 152m |
| NRC-AA-7220 | *Apis* | *dorsata* | Female(worker) | Nyaton Kitnya | 2019 | 5 | 18 | India | 28.020 N | 95.340 E | 152m |
| NRC-AA-7221 | *Apis* | *dorsata* | Female(worker) | Nyaton Kitnya | 2019 | 5 | 18 | India | 28.020 N | 95.340 E | 152m |
| NRC-AA-7222 | *Apis* | *dorsata* | Female(worker) | Nyaton Kitnya | 2019 | 5 | 9 | India | 28.215 N | 94.832 E | 240m |
| NRC-AA-7223 | *Apis* | *dorsata* | Female(worker) | Nyaton Kitnya | 2019 | 5 | 9 | India | 28.215 N | 94.832 E | 240m |
| NRC-AA-7224 | *Apis* | *dorsata* | Female(worker) | Nyaton Kitnya | 2019 | 5 | 16 | India | 28.456 N | 94.684 E | 356m |
| NRC-AA-7225 | *Apis* | *dorsata* | Female(worker) | Nyaton Kitnya | 2019 | 5 | 16 | India | 28.456 N | 94.684 E | 356m |
| NRC-AA-7226 | *Apis* | *dorsata* | Female(worker) | Nyaton Kitnya | 2019 | 5 | 16 | India | 28.456 N | 94.684 E | 356m |
| NRC-AA-7227 | *Apis* | *dorsata* | Female(worker) | Nyaton Kitnya | 2019 | 5 | 16 | India | 28.456 N | 94.684 E | 356m |
| NRC-AA-7228 | *Apis* | *dorsata* | Female(worker) | a; Gard W Otis; | 2019 | 3 | 19 | India | 28.497 | 95.088 | 558m |
| NRC-AA-7229 | *Apis* | *dorsata* | Female(worker) | a; Gard W Otis; | 2019 | 3 | 19 | India | 28.497 | 95.088 | 558m |
| NRC-AA-7230 | *Apis* | *dorsata* | Female(worker) | a; Gard W Otis; | 2019 | 3 | 19 | India | 28.497 | 95.088 | 558m |
| NRC-AA-7231 | *Apis* | *dorsata* | Female(worker) | a; Gard W Otis; | 2019 | 3 | 19 | India | 28.497 | 95.088 | 558m |
| NRC-AA-7232 | *Apis* | *dorsata* | Female(worker) | a; Gard W Otis; | 2019 | 3 | 19 | India | 28.497 | 95.088 | 558m |
| NRC-AA-7233 | *Apis* | *dorsata* | Female(worker) | a; Gard W Otis; | 2019 | 3 | 19 | India | 28.497 | 95.088 | 558m |
| NRC-AA-7234 | *Apis* | *dorsata* | Female(worker) | a; Gard W Otis; | 2019 | 3 | 19 | India | 28.497 | 95.088 | 558m |
| NRC-AA-7235 | *Apis* | *dorsata* | Female(worker) | a; Gard W Otis; | 2019 | 3 | 19 | India | 28.497 | 95.088 | 558m |
| NRC-AA-7236 | *Apis* | *dorsata* | Female(worker) | a; Gard W Otis; | 2019 | 3 | 19 | India | 28.497 | 95.088 | 558m |
| NRC-AA-7237 | *Apis* | *dorsata* | Female(worker) | Axel Brockmann | 2019 | 3 | 28 | India | 30.359 | 76.45 | 250m |
| NRC-AA-7238 | *Apis* | *dorsata* | Female(worker) | Axel Brockmann | 2019 | 3 | 28 | India | 30.359 | 76.45 | 250m |
| NRC-AA-7239 | *Apis* | *dorsata* | Female(worker) | Axel Brockmann | 2019 | 3 | 28 | India | 30.359 | 76.45 | 250m |
| NRC-AA-7240 | *Apis* | *dorsata* | Female(worker) | Axel Brockmann | 2019 | 3 | 28 | India | 30.359 | 76.45 | 250m |
| NRC-AA-7241 | *Apis* | *dorsata* | Female(worker) | Axel Brockmann | 2019 | 3 | 28 | India | 30.359 | 76.45 | 250m |
| NRC-AA-7242 | *Apis* | *dorsata* | Female(worker) | Nyaton Kitnya | 2018 | 5 | 1 | India | 25.643 | 93.753 | 460m |
| NRC-AA-7243 | *Apis* | *dorsata* | Female(worker) | Nyaton Kitnya | 2018 | 5 | 1 | India | 25.643 | 93.753 | 460m |
| NRC-AA-7244 | *Apis* | *dorsata* | Female(worker) | Nyaton Kitnya | 2018 | 5 | 1 | India | 25.643 | 93.753 | 460m |
| NRC-AA-7245 | *Apis* | *dorsata* | Female(worker) | Nyaton Kitnya | 2018 | 5 | 1 | India | 25.643 | 93.753 | 460m |
| NRC-AA-7246 | *Apis* | *dorsata* | Female(worker) | Nyaton Kitnya | 2018 | 5 | 1 | India | 25.643 | 93.753 | 460m |
| NRC-AA-7247 | *Apis* | *dorsata* | Female(worker) | Nyaton Kitnya | 2018 | 5 | 1 | India | 25.643 | 93.753 | 460m |
| NRC-AA-7248 | *Apis* | *dorsata* | Female(worker) | Nyaton Kitnya | 2018 | 5 | 1 | India | 25.643 | 93.753 | 460m |
| NRC-AA-7249 | *Apis* | *dorsata* | Female(worker) | Nyaton Kitnya | 2018 | 5 | 1 | India | 25.643 | 93.753 | 460m |
| NRC-AA-7250 | *Apis* | *dorsata* | Female(worker) | Nyaton Kitnya | 2023 | 3 | 29 | India | 13.07 N | 77.580 E | 930m |

| | | | | | | | | | | | |
|---|---|---|---|---|---|---|---|---|---|---|---|
| NRC-AA-7251 | *Apis* | *dorsata* | Female(worker) | Nyaton Kitnya | 2023 | 3 | 29 | India | 13.07 N | 77.580 E | 930m |
| NRC-AA-7252 | *Apis* | *dorsata* | Female(worker) | Nyaton Kitnya | 2023 | 3 | 29 | India | 13.07 N | 77.580 E | 930m |
| NRC-AA-7253 | *Apis* | *dorsata* | Female(worker) | Nyaton Kitnya | 2023 | 3 | 29 | India | 13.07 N | 77.580 E | 930m |
| NRC-AA-7254 | *Apis* | *dorsata* | Female(worker) | Nyaton Kitnya | 2023 | 3 | 29 | India | 13.07 N | 77.580 E | 930m |
| NRC-AA-7255 | *Apis* | *dorsata* | Female(worker) | Nyaton Kitnya | 2023 | 3 | 29 | India | 13.07 N | 77.580 E | 930m |
| NRC-AA-7256 | *Apis* | *dorsata* | Female(worker) | Nyaton Kitnya | 2023 | 3 | 29 | India | 13.07 N | 77.580 E | 930m |
| NRC-AA-7257 | *Apis* | *dorsata* | Female(worker) | Bharat Kumar K | 2022 | 4 | 15 | India | 13.07 | 77.58 | 930m |
| NRC-AA-7258 | *Apis* | *dorsata* | Female(worker) | Nyaton Kitnya | 2023 | 3 | 27 | India | 13.07 N | 77.580 E | 930m |
| NRC-AA-7259 | *Apis* | *dorsata* | Female(worker) | Nyaton Kitnya | 2023 | 3 | 27 | India | 13.07 N | 77.580 E | 930m |
| NRC-AA-7260 | *Apis* | *dorsata* | Female(worker) | Nyaton Kitnya | 2023 | 3 | 27 | India | 13.07 N | 77.580 E | 930m |
| NRC-AA-7261 | *Apis* | *dorsata* | Female(worker) | Nyaton Kitnya | 2023 | 3 | 28 | India | 13.094 N | 77.578 E | |
| NRC-AA-7262 | *Apis* | *dorsata* | Female(worker) | Nyaton Kitnya | 2023 | 3 | 28 | India | 13.094 N | 77.578 E | |
| NRC-AA-7263 | *Apis* | *dorsata* | Female(worker) | Nyaton Kitnya | 2023 | 3 | 28 | India | 13.094 N | 77.578 E | |
| NRC-AA-7264 | *Apis* | *dorsata* | Female(worker) | Nyaton Kitnya | 2023 | 3 | 28 | India | 13.094 N | 77.578 E | |
| NRC-AA-7287 | *Apis* | *dorsata* | Female(worker) | Hedge; Prabhud | *2018* | *1* | | India | 11.67 | 92.739 | 7m |
| NRC-AA-7288 | *Apis* | *dorsata* | Female(worker) | Hedge; Prabhud | *2018* | *1* | | India | 11.67 | 92.739 | 7m |
| NRC-AA-7289 | *Apis* | *dorsata* | Female(worker) | Hedge; Prabhud | *2018* | *1* | | India | 11.67 | 92.739 | 7m |
| NRC-AA-7290 | *Apis* | *dorsata* | Female(worker) | Hedge; Prabhud | *2018* | *1* | | India | 11.67 | 92.739 | 7m |
| NRC-AA-7291 | *Apis* | *dorsata* | Female(worker) | Hedge; Prabhud | *2018* | *1* | | India | 11.67 | 92.739 | 7m |
| NRC-AA-7292 | *Apis* | *dorsata* | Female(worker) | Hedge; Prabhud | *2018* | *1* | | India | 11.67 | 92.739 | 7m |
| NRC-AA-7293 | *Apis* | *dorsata* | Female(worker) | Hedge; Prabhud | *2018* | *1* | | India | 11.67 | 92.739 | 7m |
| NRC-AA-7294 | *Apis* | *dorsata* | Female(worker) | Hedge; Prabhud | *2018* | *1* | | India | 11.67 | 92.739 | 7m |
| NRC-AA-7295 | *Apis* | *dorsata* | Female(worker) | Hedge; Prabhud | *2018* | *1* | | India | 11.67 | 92.739 | 7m |
| NRC-AA-7296 | *Apis* | *dorsata* | Female(worker) | Hedge; Prabhud | *2018* | *1* | | India | 11.67 | 92.739 | 7m |
| NRC-AA-7297 | *Apis* | *dorsata* | Female(worker) | Hedge; Prabhud | *2018* | *1* | | India | l°40.599' | 2°43.78! | 152m |
| NRC-AA-7298 | *Apis* | *dorsata* | Female(worker) | Hedge; Prabhud | *2018* | *1* | | India | l°40.599' | 2°43.78! | 152m |
| NRC-AA-7299 | *Apis* | *dorsata* | Female(worker) | Hedge; Prabhud | *2018* | *1* | | India | l°40.599' | 2°43.78! | 152m |
| NRC-AA-7300 | *Apis* | *dorsata* | Female(worker) | Hedge; Prabhud | *2018* | *1* | | India | l°40.599' | 2°43.78! | 152m |
| NRC-AA-7301 | *Apis* | *dorsata* | Female(worker) | Hedge; Prabhud | *2018* | *1* | | India | l°40.599' | 2°43.78! | 152m |
| NRC-AA-7302 | *Apis* | *dorsata* | Female(worker) | Hedge; Prabhud | *2018* | *1* | | India | l°40.599' | 2°43.78! | 152m |
| NRC-AA-7303 | *Apis* | *dorsata* | Female(worker) | Hedge; Prabhud | *2018* | *1* | | India | l°40.599' | 2°43.78! | 152m |
| NRC-AA-7304 | *Apis* | *dorsata* | Female(worker) | Hedge; Prabhud | *2018* | *1* | | India | l°40.599' | 2°43.78! | 152m |
| NRC-AA-7305 | *Apis* | *dorsata* | Female(worker) | Hedge; Prabhud | *2018* | *1* | | India | l°40.599' | 2°43.78! | 152m |
| NRC-AA-7306 | *Apis* | *dorsata* | Female(worker) | Hedge; Prabhud | *2018* | *1* | | India | l°40.599' | 2°43.78! | 152m |
| NRC-AA-7307 | *Apis* | *dorsata* | Female(worker) | Hedge; Prabhud | 2018 | 1 | 19 | India | 12.506 | 92.914 | 22m |
| NRC-AA-7308 | *Apis* | *dorsata* | Female(worker) | Hedge; Prabhud | 2018 | 1 | 19 | India | 12.506 | 92.914 | 22m |
| NRC-AA-7309 | *Apis* | *dorsata* | Female(worker) | Hedge; Prabhud | 2018 | 1 | 19 | India | 12.506 | 92.914 | 22m |
| NRC-AA-7310 | *Apis* | *dorsata* | Female(worker) | Hedge; Prabhud | 2018 | 1 | 19 | India | 12.506 | 92.914 | 22m |
| NRC-AA-7311 | *Apis* | *dorsata* | Female(worker) | Hedge; Prabhud | 2018 | 1 | 19 | India | 12.506 | 92.914 | 22m |
| NRC-AA-7312 | *Apis* | *dorsata* | Female(worker) | Hedge; Prabhud | 2018 | 1 | 19 | India | 13°13.59 | 93°02.70 | 5m |
| NRC-AA-7313 | *Apis* | *dorsata* | Female(worker) | Hedge; Prabhud | 2018 | 1 | 19 | India | 13°13.59 | 93°02.70 | 5m |
| NRC-AA-7314 | *Apis* | *dorsata* | Female(worker) | Hedge; Prabhud | 2018 | 1 | 19 | India | 13°13.59 | 93°02.70 | 5m |
| NRC-AA-7315 | *Apis* | *dorsata* | Female(worker) | Hedge; Prabhud | 2018 | 1 | 19 | India | 13°13.59 | 93°02.70 | 5m |
| NRC-AA-7316 | *Apis* | *dorsata* | Female(worker) | Hedge; Prabhud | 2018 | 1 | 19 | India | 13°13.59 | 93°02.70 | 5m |
| NRC-AA-7317 | *Apis* | *dorsata* | Female(worker) | Sachin Sahu | 2019 | 4 | 15 | India | 22.965 N | 88.525 E | |
| NRC-AA-7318 | *Apis* | *dorsata* | Female(worker) | Sachin Sahu | 2019 | 4 | 15 | India | 22.965 N | 88.525 E | |
| NRC-AA-7319 | *Apis* | *dorsata* | Female(worker) | Sachin Sahu | 2019 | 4 | 15 | India | 22.965 N | 88.525 E | |
| NRC-AA-7320 | *Apis* | *dorsata* | Female(worker) | Sachin Sahu | 2019 | 4 | 15 | India | 22.965 N | 88.525 E | |
| NRC-AA-7321 | *Apis* | *dorsata* | Female(worker) | Sachin Sahu | 2019 | 4 | 15 | India | 22.965 N | 88.525 E | |
| NRC-AA-7322 | *Apis* | *dorsata* | Female(worker) | Sachin Sahu | 2019 | 4 | 15 | India | 22.965 N | 88.525 E | |
| NRC-AA-7323 | *Apis* | *dorsata* | Female(worker) | Sachin Sahu | 2019 | 4 | 15 | India | 22.965 N | 88.525 E | |
| NRC-AA-7324 | *Apis* | *dorsata* | Female(worker) | Sachin Sahu | 2019 | 4 | 15 | India | 22.965 N | 88.525 E | |

| | | | | | | | | | | | |
|---|---|---|---|---|---|---|---|---|---|---|---|
| NRC-AA-7325 | *Apis* | *dorsata* | Female(worker) | Sachin Sahu | 2019 | 4 | 15 | India | 22.965 N | 88.525 E | |
| NRC-AA-7326 | *Apis* | *dorsata* | Female(worker) | Sachin Sahu | 2019 | 4 | 15 | India | 22.965 N | 88.525 E | |
| NRC-AA-7327 | *Apis* | *dorsata* | Female(worker) | Sachin Sahu | 2019 | 4 | 15 | India | 22.965 N | 88.525 E | |
| NRC-AA-7328 | *Apis* | *dorsata* | Female(worker) | Sachin Sahu | 2019 | 4 | 15 | India | 22.965 N | 88.525 E | |
| NRC-AA-7329 | *Apis* | *dorsata* | Female(worker) | Prabhudev MV | 2019 | 4 | 28 | India | 29.751 | 80.379 | 642m |
| NRC-AA-7330 | *Apis* | *dorsata* | Female(worker) | Prabhudev MV | 2019 | 4 | 28 | India | 29.751 | 80.379 | 642m |
| NRC-AA-7331 | *Apis* | *dorsata* | Female(worker) | Prabhudev MV | 2019 | 4 | 28 | India | 29.751 | 80.379 | 642m |
| NRC-AA-7332 | *Apis* | *dorsata* | Female(worker) | Prabhudev MV | 2019 | 4 | 28 | India | 29.751 | 80.379 | 642m |
| NRC-AA-7333 | *Apis* | *dorsata* | Female(worker) | Prabhudev MV | 2019 | 4 | 28 | India | 29.751 | 80.379 | 642m |
| NRC-AA-7334 | *Apis* | *dorsata* | Female(worker) | Prabhudev MV | 2019 | 4 | 28 | India | 29.751 | 80.379 | 642m |
| NRC-AA-7335 | *Apis* | *dorsata* | Female(worker) | Prabhudev MV | 2019 | 4 | 28 | India | 29.751 | 80.379 | 642m |
| NRC-AA-7336 | *Apis* | *dorsata* | Female(worker) | Prabhudev MV | 2019 | 4 | 28 | India | 29.751 | 80.379 | 642m |
| NRC-AA-7337 | *Apis* | *dorsata* | Female(worker) | Prabhudev MV | 2019 | 4 | 28 | India | 29.751 | 80.379 | 642m |
| NRC-AA-7338 | *Apis* | *dorsata* | Female(worker) | Prabhudev MV | 2019 | 4 | 28 | India | 29.751 | 80.379 | 642m |
| NRC-AA-7339 | *Apis* | *dorsata* | Female(worker) | Axel Brockman | 2019 | 3 | 28 | India | 30.359 | 76.45 | 250m |
| NRC-AA-7340 | *Apis* | *dorsata* | Female(worker) | Axel Brockman | 2019 | 3 | 28 | India | 30.359 | 76.45 | 250m |
| NRC-AA-7341 | *Apis* | *dorsata* | Female(worker) | Axel Brockman | 2019 | 3 | 28 | India | 30.359 | 76.45 | 250m |
| NRC-AA-7342 | *Apis* | *dorsata* | Female(worker) | Axel Brockman | 2019 | 3 | 28 | India | 30.359 | 76.45 | 250m |
| NRC-AA-7343 | *Apis* | *dorsata* | Female(worker) | Axel Brockman | 2019 | 3 | 28 | India | 30.359 | 76.45 | 250m |
| NRC-AA-7344 | *Apis* | *dorsata* | Female(worker) | Axel Brockman | 2019 | 3 | 28 | India | 30.359 | 76.45 | 250m |
| NRC-AA-7345 | *Apis* | *dorsata* | Female(worker) | Axel Brockman | 2019 | 3 | 28 | India | 30.359 | 76.45 | 250m |
| NRC-AA-7346 | *Apis* | *dorsata* | Female(worker) | Axel Brockman | 2019 | 3 | 28 | India | 30.359 | 76.45 | 250m |
| NRC-AA-7347 | *Apis* | *dorsata* | Female(worker) | Axel Brockman | 2019 | 3 | 28 | India | 30.359 | 76.45 | 250m |
| NRC-AA-7348 | *Apis* | *dorsata* | Female(worker) | Axel Brockman | 2019 | 3 | 28 | India | 30.359 | 76.45 | 250m |
| NRC-AA-7349 | *Apis* | *dorsata* | Female(worker) | Geetha T | 2018 | 2 | 27 | India | 13.150 N | 77.060 E | 887m |
| NRC-AA-7350 | *Apis* | *dorsata* | Female(worker) | Geetha T | 2018 | 2 | 27 | India | 13.150 N | 77.060 E | 887m |
| NRC-AA-7351 | *Apis* | *dorsata* | Female(worker) | Geetha T | 2018 | 2 | 27 | India | 13.030 N | 77.520 E | 918m |
| NRC-AA-7352 | *Apis* | *dorsata* | Female(worker) | Nyaton Kitnya | 2018 | 1 | 13 | India | 13.07 N | 77.580 E | 930m |
| NRC-AA-7353 | *Apis* | *dorsata* | Female(worker) | Nyaton Kitnya | 2018 | 1 | 13 | India | 13.07 N | 77.580 E | 930m |
| NRC-AA-7354 | *Apis* | *dorsata* | Female(worker) | Nyaton Kitnya | 2018 | 1 | 13 | India | 13.07 N | 77.580 E | 930m |
| NRC-AA-7355 | *Apis* | *dorsata* | Female(worker) | Nyaton Kitnya | 2018 | 1 | 13 | India | 13.07 N | 77.580 E | 930m |
| NRC-AA-7356 | *Apis* | *dorsata* | Female(worker) | Nyaton Kitnya | 2018 | 1 | 13 | India | 13.07 N | 77.580 E | 930m |
| NRC-AA-7357 | *Apis* | *dorsata* | Female(worker) | Nyaton Kitnya | 2018 | 1 | 13 | India | 13.07 N | 77.580 E | 930m |
| NRC-AA-7358 | *Apis* | *dorsata* | Female(worker) | Jaya Narah | 2018 | 4 | 24 | India | 24°49'10 | 93°56'6 | 779m |
| NRC-AA-7359 | *Apis* | *dorsata* | Female(worker) | Jaya Narah | 2018 | 4 | 24 | India | 24°49'10 | 93°56'6 | 779m |
| NRC-AA-7360 | *Apis* | *dorsata* | Female(worker) | Jaya Narah | 2018 | 4 | 24 | India | 24°49'10 | 93°56'6 | 779m |
| NRC-AA-7361 | *Apis* | *dorsata* | Female(worker) | Jaya Narah | 2018 | 4 | 24 | India | 24°48'46 | 93°56'0 | 775m |
| NRC-AA-7362 | *Apis* | *dorsata* | Female(worker) | Jaya Narah | 2018 | 4 | 24 | India | 24°48'46 | 93°56'0 | 775m |
| NRC-AA-7363 | *Apis* | *dorsata* | Female(worker) | Jaya Narah | 2018 | 4 | 24 | India | 24°48'46 | 93°56'0 | 775m |
| NRC-AA-7364 | *Apis* | *dorsata* | Female(worker) | Jaya Narah | 2018 | 4 | 24 | India | 24°48'46 | 93°56'0 | 775m |
| NRC-AA-7365 | *Apis* | *dorsata* | Female(worker) | Jaya Narah | 2018 | 4 | 24 | India | 24°48'46 | 93°56'0 | 775m |
| NRC-AA-7366 | *Apis* | *dorsata* | Female(worker) | Jaya Narah | 2018 | 4 | 24 | India | 24°48'46 | 93°56'0 | 775m |
| NRC-AA-7367 | *Apis* | *dorsata* | Female(worker) | Jaya Narah | 2018 | 4 | 24 | India | 24°48'46 | 93°56'0 | 775m |
| NRC-AA-7368 | *Apis* | *dorsata* | Female(worker) | Jaya Narah | 2018 | 4 | 24 | India | 24°48'46 | 93°56'0 | 775m |
| NRC-AA-7369 | *Apis* | *dorsata* | Female(worker) | Jaya Narah | 2018 | 4 | 24 | India | 24°48'46 | 93°56'0 | 775m |
| NRC-AA-7370 | *Apis* | *dorsata* | Female(worker) | Jaya Narah | 2018 | 4 | 24 | India | 24°48'46 | 93°56'0 | 775m |
| NRC-AA-7371 | *Apis* | *dorsata* | Female(worker) | Nyaton Kitnya | 2018 | 3 | 5 | India | 25.550 N | 93.445 E | 118 |
| NRC-AA-7372 | *Apis* | *dorsata* | Female(worker) | Nyaton Kitnya | 2018 | 3 | 5 | India | 25.550 N | 93.445 E | 118 |
| NRC-AA-7373 | *Apis* | *dorsata* | Female(worker) | Nyaton Kitnya | 2018 | 3 | 5 | India | 25.550 N | 93.445 E | 118 |
| NRC-AA-7374 | *Apis* | *dorsata* | Female(worker) | Nyaton Kitnya | 2018 | 3 | 5 | India | 25.550 N | 93.445 E | 118 |
| NRC-AA-7375 | *Apis* | *dorsata* | Female(worker) | Nyaton Kitnya | 2018 | 3 | 5 | India | 25.550 N | 93.445 E | 118 |
| NRC-AA-7376 | *Apis* | *dorsata* | Female(worker) | Nyaton Kitnya | 2018 | 3 | 5 | India | 25.550 N | 93.445 E | 118 |

| | | | | | | | | | | | |
|---|---|---|---|---|---|---|---|---|---|---|---|
| NRC-AA-7377 | *Apis* | *dorsata* | Female(worker) | Nyaton Kitnya | 2019 | 6 | 15 | India | 25.404 N | 93.464 E | 496 |
| NRC-AA-7378 | *Apis* | *dorsata* | Female(worker) | Nyaton Kitnya | 2019 | 6 | 15 | India | 25.404 N | 93.464 E | 496 |
| NRC-AA-7379 | *Apis* | *dorsata* | Female(worker) | Nyaton Kitnya | 2019 | 6 | 15 | India | 25.404 N | 93.464 E | 496 |
| NRC-AA-7380 | *Apis* | *dorsata* | Female(worker) | Nyaton Kitnya | 2019 | 6 | 15 | India | 25.404 N | 93.464 E | 496 |
| NRC-AA-7381 | *Apis* | *dorsata* | Female(worker) | Nyaton Kitnya | 2019 | 6 | 15 | India | 25.404 N | 93.464 E | 496 |
| NRC-AA-7382 | *Apis* | *dorsata* | Female(worker) | Nyaton Kitnya | 2019 | 6 | 15 | India | 25.404 N | 93.464 E | 496 |
| NRC-AA-7383 | *Apis* | *dorsata* | Female(worker) | Nyaton Kitnya | 2019 | 6 | 15 | India | 25.404 N | 93.464 E | 496 |
| NRC-AA-7384 | *Apis* | *dorsata* | Female(worker) | Nyaton Kitnya | 2019 | 6 | 15 | India | 25.404 N | 93.464 E | 496 |
| NRC-AA-7385 | *Apis* | *dorsata* | Female(worker) | Nyaton Kitnya | 2019 | 6 | 15 | India | 25.404 N | 93.464 E | 496 |
| NRC-AA-7386 | *Apis* | *dorsata* | Female(worker) | Nyaton Kitnya | 2019 | 6 | 15 | India | 25.404 N | 93.464 E | 496 |
| NRC-AA-7387 | *Apis* | *dorsata* | Female(worker) | Nyaton Kitnya | 2018 | 5 | 1 | India | 25.643 | 93.753 | 460m |
| NRC-AA-7388 | *Apis* | *dorsata* | Female(worker) | Nyaton Kitnya | 2018 | 5 | 1 | India | 25.643 | 93.753 | 460m |
| NRC-AA-7389 | *Apis* | *dorsata* | Female(worker) | Nyaton Kitnya | 2018 | 5 | 1 | India | 25.643 | 93.753 | 460m |
| NRC-AA-7390 | *Apis* | *dorsata* | Female(worker) | Nyaton Kitnya | 2018 | 5 | 1 | India | 25.643 | 93.753 | 460m |
| NRC-AA-7391 | *Apis* | *dorsata* | Female(worker) | Nyaton Kitnya | 2018 | 5 | 1 | India | 25.643 | 93.753 | 460m |
| NRC-AA-7392 | *Apis* | *dorsata* | Female(worker) | Nyaton Kitnya | 2018 | 5 | 1 | India | 25.643 | 93.753 | 460m |
| NRC-AA-7393 | *Apis* | *dorsata* | Female(worker) | Nyaton Kitnya | 2018 | 5 | 1 | India | 25.643 | 93.753 | 460m |
| NRC-AA-7394 | *Apis* | *dorsata* | Female(worker) | Nyaton Kitnya | 2018 | 5 | 1 | India | 25.643 | 93.753 | 460m |
| NRC-AA-7395 | *Apis* | *dorsata* | Female(worker) | Nyaton Kitnya | 2018 | 5 | 1 | India | 25.643 | 93.753 | 460m |
| NRC-AA-7396 | *Apis* | *dorsata* | Female(worker) | Nyaton Kitnya | 2018 | 5 | 1 | India | 25.643 | 93.753 | 460m |
| NRC-AA-7397 | *Apis* | *dorsata* | Female(worker) | Nyaton Kitnya | 2018 | 3 | 5 | India | 25.550 N | 93.445 E | 118 |
| NRC-AA-7398 | *Apis* | *dorsata* | Female(worker) | Nyaton Kitnya | 2018 | 3 | 5 | India | 25.550 N | 93.445 E | 118 |
| NRC-AA-7399 | *Apis* | *dorsata* | Female(worker) | Nyaton Kitnya | 2018 | 3 | 5 | India | 25.550 N | 93.445 E | 118 |
| NRC-AA-7400 | *Apis* | *dorsata* | Female(worker) | Nyaton Kitnya | 2018 | 3 | 5 | India | 25.550 N | 93.445 E | 118 |
| NRC-AA-7401 | *Apis* | *dorsata* | Female(worker) | Nyaton Kitnya | 2019 | 5 | 24 | India | 5'38.369 | 3'45.133 | 532 |
| NRC-AA-7402 | *Apis* | *dorsata* | Female(worker) | Nyaton Kitnya | 2019 | 5 | 24 | India | 5'38.369 | 3'45.133 | 532 |
| NRC-AA-7403 | *Apis* | *dorsata* | Female(worker) | Nyaton Kitnya | 2019 | 5 | 24 | India | 5'38.369 | 3'45.133 | 532 |
| NRC-AA-7404 | *Apis* | *dorsata* | Female(worker) | Nyaton Kitnya | 2019 | 5 | 24 | India | 5'38.369 | 3'45.133 | 532 |
| NRC-AA-7405 | *Apis* | *dorsata* | Female(worker) | Nyaton Kitnya | 2019 | 5 | 24 | India | 5'38.369 | 3'45.133 | 532 |
| NRC-AA-7406 | *Apis* | *dorsata* | Female(worker) | Nyaton Kitnya | 2019 | 5 | 24 | India | 5'38.369 | 3'45.133 | 532 |
| NRC-AA-7407 | *Apis* | *dorsata* | Female(worker) | Nyaton Kitnya | 2019 | 5 | 24 | India | 5'38.369 | 3'45.133 | 532 |
| NRC-AA-7408 | *Apis* | *dorsata* | Female(worker) | Nyaton Kitnya | 2019 | 5 | 24 | India | 5'38.369 | 3'45.133 | 532 |
| NRC-AA-7409 | *Apis* | *dorsata* | Female(worker) | Nyaton Kitnya | 2019 | 5 | 24 | India | 5'38.369 | 3'45.133 | 532 |
| NRC-AA-7410 | *Apis* | *dorsata* | Female(worker) | Nyaton Kitnya | 2019 | 5 | 24 | India | 5'38.369 | 3'45.133 | 532 |
| NRC-AA-7411 | *Apis* | *dorsata* | Female(worker) | Nyaton Kitnya | 2019 | 3 | 29 | India | 7'08.942 | 3'45.807 | 249 |
| NRC-AA-7412 | *Apis* | *dorsata* | Female(worker) | Nyaton Kitnya | 2019 | 3 | 29 | India | 7'08.942 | 3'45.807 | 249 |
| NRC-AA-7413 | *Apis* | *dorsata* | Female(worker) | Nyaton Kitnya | 2018 | 4 | 21 | India | 28.281 | 94.851 | 500m |
| NRC-AA-7414 | *Apis* | *dorsata* | Female(worker) | Nyaton Kitnya | 2018 | 4 | 21 | India | 28.281 | 94.851 | 500m |
| NRC-AA-7415 | *Apis* | *dorsata* | Female(worker) | Nyaton Kitnya | 2018 | 4 | 21 | India | 28.281 | 94.851 | 500m |
| NRC-AA-7416 | *Apis* | *dorsata* | Female(worker) | Nyaton Kitnya | 2018 | 4 | 21 | India | 28.281 | 94.851 | 500m |
| NRC-AA-7417 | *Apis* | *dorsata* | Female(worker) | Nyaton Kitnya | 2018 | 4 | 21 | India | 28.281 | 94.851 | 500m |
| NRC-AA-7418 | *Apis* | *dorsata* | Female(worker) | Nyaton Kitnya | 2018 | 4 | 21 | India | 28.281 | 94.851 | 500m |
| NRC-AA-7419 | *Apis* | *dorsata* | Female(worker) | Nyaton Kitnya | 2018 | 4 | 21 | India | 28.281 | 94.851 | 500m |
| NRC-AA-7420 | *Apis* | *dorsata* | Female(worker) | Nyaton Kitnya | 2018 | 4 | 21 | India | 28.281 | 94.851 | 500m |
| NRC-AA-7421 | *Apis* | *dorsata* | Female(worker) | Nyaton Kitnya | 2018 | 4 | 21 | India | 28.281 | 94.851 | 500m |
| NRC-AA-7422 | *Apis* | *dorsata* | Female(worker) | Nyaton Kitnya | 2019 | 5 | 15 | India | 28.438 | 94.67 | 480m |
| NRC-AA-7423 | *Apis* | *dorsata* | Female(worker) | Nyaton Kitnya | 2019 | 5 | 15 | India | 28.438 | 94.67 | 480m |
| NRC-AA-7424 | *Apis* | *dorsata* | Female(worker) | Nyaton Kitnya | 2019 | 5 | 15 | India | 28.438 | 94.67 | 480m |
| NRC-AA-7425 | *Apis* | *dorsata* | Female(worker) | Nyaton Kitnya | 2019 | 5 | 15 | India | 28.438 | 94.67 | 480m |
| NRC-AA-7426 | *Apis* | *dorsata* | Female(worker) | Nyaton Kitnya | 2019 | 5 | 15 | India | 28.438 | 94.67 | 480m |
| NRC-AA-7427 | *Apis* | *dorsata* | Female(worker) | Nyaton Kitnya | 2019 | 5 | 15 | India | 28.438 | 94.67 | 480m |
| NRC-AA-7428 | *Apis* | *dorsata* | Female(worker) | Nyaton Kitnya | 2019 | 5 | 15 | India | 28.438 | 94.67 | 480m |

| | | | | | | | | | | | |
|---|---|---|---|---|---|---|---|---|---|---|---|
| NRC-AA-7429 | *Apis* | *dorsata* | Female(worker) | Nyaton Kitnya | 2019 | 5 | 15 | India | 28.438 | 94.67 | 480m |
| NRC-AA-7430 | *Apis* | *dorsata* | Female(worker) | Nyaton Kitnya | 2019 | 5 | 15 | India | 28.438 | 94.67 | 480m |
| NRC-AA-7431 | *Apis* | *dorsata* | Female(worker) | Nyaton Kitnya | 2019 | 5 | 15 | India | 28.438 | 94.67 | 480m |
| NRC-AA-7432 | *Apis* | *dorsata* | Female(worker) | Nyaton Kitnya | 2019 | 3 | 28 | India | 28.497 | 95.088 | 558m |
| NRC-AA-7433 | *Apis* | *dorsata* | Female(worker) | Nyaton Kitnya | 2019 | 3 | 28 | India | 28.497 | 95.088 | 558m |
| NRC-AA-7434 | *Apis* | *dorsata* | Female(worker) | Nyaton Kitnya | 2019 | 3 | 28 | India | 28.497 | 95.088 | 558m |
| NRC-AA-7435 | *Apis* | *dorsata* | Female(worker) | Nyaton Kitnya | 2019 | 3 | 28 | India | 28.497 | 95.088 | 558m |
| NRC-AA-7436 | *Apis* | *dorsata* | Female(worker) | Nyaton Kitnya | 2019 | 3 | 28 | India | 28.497 | 95.088 | 558m |
| NRC-AA-7437 | *Apis* | *dorsata* | Female(worker) | Nyaton Kitnya | 2019 | 3 | 28 | India | 28.497 | 95.088 | 558m |
| NRC-AA-7438 | *Apis* | *dorsata* | Female(worker) | Nyaton Kitnya | 2019 | 3 | 28 | India | 28.497 | 95.088 | 558m |
| NRC-AA-7439 | *Apis* | *dorsata* | Female(worker) | Nyaton Kitnya | 2019 | 3 | 28 | India | 28.497 | 95.088 | 558m |
| NRC-AA-7440 | *Apis* | *dorsata* | Female(worker) | Nyaton Kitnya | 2019 | 3 | 28 | India | 28.497 | 95.088 | 558m |
| NRC-AA-7441 | *Apis* | *dorsata* | Female(worker) | Nyaton Kitnya | 2019 | 3 | 28 | India | 28.497 | 95.088 | 558m |
| NRC-AA-7442 | *Apis* | *dorsata* | Female(worker) | Nyaton Kitnya | 2019 | 3 | 27 | India | 28.215 N | 94.832 E | 240m |
| NRC-AA-7443 | *Apis* | *dorsata* | Female(worker) | Nyaton Kitnya | 2019 | 3 | 27 | India | 28.215 N | 94.832 E | 240m |
| NRC-AA-7444 | *Apis* | *dorsata* | Female(worker) | Nyaton Kitnya | 2019 | 3 | 29 | India | 28.215 N | 94.832 E | 240m |
| NRC-AA-7445 | *Apis* | *dorsata* | Female(worker) | Nyaton Kitnya | 2019 | 5 | 9 | India | 28.215 N | 94.832 E | 240m |
| NRC-AA-7446 | *Apis* | *dorsata* | Female(worker) | Nyaton Kitnya | 2019 | 5 | 9 | India | 28.215 N | 94.832 E | 240m |
| NRC-AA-7447 | *Apis* | *dorsata* | Female(worker) | Nyaton Kitnya | 2019 | 5 | 9 | India | 28.215 N | 94.832 E | 240m |
| NRC-AA-7448 | *Apis* | *dorsata* | Female(worker) | Nyaton Kitnya | 2019 | 5 | 9 | India | 28.215 N | 94.832 E | 240m |
| NRC-AA-7449 | *Apis* | *dorsata* | Female(worker) | Nyaton Kitnya | 2019 | 5 | 9 | India | 28.215 N | 94.832 E | 240m |
| NRC-AA-7450 | *Apis* | *dorsata* | Female(worker) | Nyaton Kitnya | 2019 | 5 | 9 | India | 28.215 N | 94.832 E | 240m |
| NRC-AA-7451 | *Apis* | *dorsata* | Female(worker) | Nyaton Kitnya | 2019 | 5 | 9 | India | 28.215 N | 94.832 E | 240m |
| NRC-AA-7452 | *Apis* | *dorsata* | Female(worker) | Nyaton Kitnya | 2019 | 5 | 9 | India | 28.215 N | 94.832 E | 240m |
| NRC-AA-7453 | *Apis* | *dorsata* | Female(worker) | Nyaton Kitnya | 2019 | 5 | 9 | India | 28.215 N | 94.832 E | 240m |
| NRC-AA-7454 | *Apis* | *dorsata* | Female(worker) | Nyaton Kitnya | 2019 | 5 | 9 | India | 28.215 N | 94.832 E | 240m |
| NRC-AA-7455 | *Apis* | *dorsata* | Female(worker) | ıjamin Rutschm | 2018 | 2 | 21 | Vietnam | 10.189 | 103.97 | 0m |
| NRC-AA-7456 | *Apis* | *dorsata* | Female(worker) | ıjamin Rutschm | 2018 | 2 | 21 | Vietnam | 10.189 | 103.97 | 0m |
| NRC-AA-7457 | *Apis* | *dorsata* | Female(worker) | ıjamin Rutschm | 2018 | 2 | 21 | Vietnam | 10.189 | 103.97 | 0m |
| NRC-AA-7458 | *Apis* | *dorsata* | Female(worker) | ıjamin Rutschm | 2018 | 2 | 21 | Vietnam | 10.189 | 103.97 | 0m |
| NRC-AA-7459 | *Apis* | *dorsata* | Female(worker) | ıjamin Rutschm | 2018 | 2 | 21 | Vietnam | 10.189 | 103.97 | 0m |
| NRC-AA-7460 | *Apis* | *dorsata* | Female(worker) | ıjamin Rutschm | 2018 | 2 | 21 | Vietnam | 10.189 | 103.97 | 0m |
| NRC-AA-7461 | *Apis* | *dorsata* | Female(worker) | ıjamin Rutschm | 2018 | 2 | 21 | Vietnam | 10.189 | 103.97 | 0m |
| NRC-AA-7462 | *Apis* | *dorsata* | Female(worker) | ıjamin Rutschm | 2018 | 2 | 21 | Vietnam | 10.189 | 103.97 | 0m |
| NRC-AA-7463 | *Apis* | *dorsata* | Female(worker) | ıjamin Rutschm | 2018 | 2 | 21 | Vietnam | 10.189 | 103.97 | 0m |
| NRC-AA-7464 | *Apis* | *dorsata* | Female(worker) | ıjamin Rutschm | 2018 | 2 | 21 | Vietnam | 10.189 | 103.97 | 0m |
| NRC-AA-7465 | *Apis* | *dorsata* | Female(worker) | | 1994 | | | Vietnam | | | |
| NRC-AA-7466 | *Apis* | *dorsata* | Female(worker) | | 1994 | | | Vietnam | | | |
| NRC-AA-7467 | *Apis* | *dorsata* | Female(worker) | | 1994 | | | Vietnam | | | |
| NRC-AA-7468 | *Apis* | *dorsata* | Female(worker) | | 1994 | | | Vietnam | | | |
| NRC-AA-7469 | *Apis* | *dorsata* | Female(worker) | | 1994 | | | Vietnam | | | |
| NRC-AA-7470 | *Apis* | *dorsata* | Female(worker) | Prasad Bhatta | (CPB) | | | Nepal | | | |
| NRC-AA-7471 | *Apis* | *dorsata* | Female(worker) | Prasad Bhatta | (CPB) | | | Nepal | | | |
| NRC-AA-7472 | *Apis* | *dorsata* | Female(worker) | Prasad Bhatta | (CPB) | | | Nepal | | | |
| NRC-AA-7473 | *Apis* | *dorsata* | Female(worker) | Prasad Bhatta | (CPB) | | | Nepal | | | |
| NRC-AA-7474 | *Apis* | *dorsata* | Female(worker) | Prasad Bhatta | (CPB) | | | Nepal | | | |
| NRC-AA-7475 | *Apis* | *dorsata* | Female(worker) | Prasad Bhatta | (CPB) | | | Nepal | | | |
| NRC-AA-7670 | *Apis* | *dorsata* | Female(worker) | Nyaton Kitnya | 2022 | 1 | 20 | India | 13.07 | 77.575 | 930m |
| NRC-AA-7671 | *Apis* | *dorsata* | Female(worker) | Nyaton Kitnya | 2022 | 1 | 18 | India | 13.07 | 77.575 | 930m |
| NRC-AA-7672 | *Apis* | *dorsata* | Female(worker) | Prabhudev MV | 2018 | 1 | 19 | India | 12.506 | 92.914 | 22m |
| NRC-AA-7673 | *Apis* | *dorsata* | Female(worker) | Prabhudev MV | 2018 | 1 | 19 | India | 12.506 | 92.914 | 22m |
| NRC-AA-7674 | *Apis* | *dorsata* | Female(worker) | Prabhudev MV | 2018 | 1 | 19 | India | 12.506 | 92.914 | 22m |

| | | | | | | | | | | | |
|---|---|---|---|---|---|---|---|---|---|---|---|
| NRC-AA-7675 | *Apis* | *dorsata* | Female(worker) | Prabhudev MV | 2018 | 1 | 19 | India | °40.599' | 2°43.78 | 152m |
| NRC-AA-7676 | *Apis* | *dorsata* | Female(worker) | Prabhudev MV | *2018* | *1* | | India | °40.599' | 2°43.78 | 152m |
| NRC-AA-7677 | *Apis* | *dorsata* | Female(worker) | Prabhudev MV | *2018* | *1* | | India | °40.599' | 2°43.78 | 152m |
| NRC-AA-7678 | *Apis* | *dorsata* | Female(worker) | Prabhudev MV | *2018* | *1* | | India | °40.599' | 2°43.78 | 152m |
| NRC-AA-7679 | *Apis* | *dorsata* | Female(worker) | Prabhudev MV | *2018* | *1* | | India | °40.599' | 2°43.78 | 152m |
| NRC-AA-7680 | *Apis* | *dorsata* | Female(worker) | Prabhudev MV | *2018* | *1* | | India | °40.599' | 2°43.78 | 152m |
| NRC-AA-7681 | *Apis* | *dorsata* | Female(worker) | Prabhudev MV | *2018* | *1* | | India | °40.599' | 2°43.78 | 152m |
| NRC-AA-7682 | *Apis* | *dorsata* | Female(worker) | Prabhudev MV | *2018* | *1* | | India | °40.599' | 2°43.78 | 152m |
| NRC-AA-7683 | *Apis* | *dorsata* | Female(worker) | Prabhudev MV | 2018 | 1 | | India | 11.67 | 92.739 | 7m |
| NRC-AA-7684 | *Apis* | *dorsata* | Female(worker) | Prabhudev MV | 2018 | 1 | | India | 11.67 | 92.739 | 7m |
| NRC-AA-7685 | *Apis* | *dorsata* | Female(worker) | Prabhudev MV | 2018 | 1 | | India | 11.67 | 92.739 | 7m |
| NRC-AA-7686 | *Apis* | *dorsata* | Female(worker) | Prabhudev MV | 2018 | 1 | | India | 11.67 | 92.739 | 7m |
| NRC-AA-8016 | *Apis* | *dorsata* | Female(worker) | n Kitnya, Jaya | 2018 | 6 | 20 | India | 27.107 | 92.525 | |
| NRC-AA-8017 | *Apis* | *dorsata* | Female(worker) | n Kitnya, Jaya | 2018 | 6 | 20 | India | 27.107 | 92.525 | |
| NRC-AA-8018 | *Apis* | *dorsata* | Male (drone) | CPB | | | | Nepal | | | |
| NRC-AA-8068 | *Apis* | *dorsata* | Male (drone) | Nyaton Kitnya | 2019 | 5 | 16 | India | 28.456 | 94.684 | 356m |
| NRC-AA-8069 | *Apis* | *dorsata* | Male (drone) | Nyaton Kitnya | 2019 | 5 | 16 | India | 28.456 | 94.684 | 356m |
| NRC-AA-8070 | *Apis* | *dorsata* | Male (drone) | Nyaton Kitnya | 2019 | 5 | 16 | India | 28.456 | 94.684 | 356m |
| NRC-AA-8071 | *Apis* | *dorsata* | Male (drone) | Nyaton Kitnya | 2019 | 5 | 16 | India | 28.456 | 94.684 | 356m |
| NRC-AA-8072 | *Apis* | *dorsata* | Male (drone) | Nyaton Kitnya | 2019 | 5 | 16 | India | 28.456 | 94.684 | 356m |
| NRC-AA-8073 | *Apis* | *dorsata* | Male (drone) | Nyaton Kitnya | 2019 | 5 | 16 | India | 28.456 | 94.684 | 356m |
| NRC-AA-8074 | *Apis* | *dorsata* | Male (drone) | Nyaton Kitnya | 2019 | 5 | 16 | India | 28.456 | 94.684 | 356m |
| NRC-AA-8075 | *Apis* | *dorsata* | Male (drone) | Nyaton Kitnya | 2019 | 5 | 16 | India | 28.456 | 94.684 | 356m |
| NRC-AA-8076 | *Apis* | *dorsata* | Male (drone) | Nyaton Kitnya | 2019 | 5 | 16 | India | 28.456 | 94.684 | 356m |
| NRC-AA-8077 | *Apis* | *dorsata* | Male (drone) | Nyaton Kitnya | 2019 | 5 | 16 | India | 28.456 | 94.684 | 356m |
| NRC-AA-8078 | *Apis* | *dorsata* | Male (drone) | Benjamin | 2018 | 2 | 21 | Vietnam | | | |
| NRC-AA-8079 | *Apis* | *dorsata* | Male (drone) | Benjamin | 2018 | 2 | 21 | Vietnam | | | |
| NCBS-BF225 | *Apis* | *dorsata* | Female(worker) | Nyaton Kitnya | 2017 | 11 | 12 | India | °57'16.3 | °41'19.3 | 1200m |
| NCBS-BF226 | *Apis* | *dorsata* | Female(worker) | Nyaton Kitnya | 2017 | 11 | 12 | India | °57'16.3 | °41'19.3 | 1200m |
| NCBS-BF227 | *Apis* | *dorsata* | Female(worker) | Nyaton Kitnya | 2017 | 11 | 12 | India | °57'16.3 | °41'19.3 | 1200m |
| NCBS-BF228 | *Apis* | *dorsata* | Female(worker) | Nyaton Kitnya | 2017 | 11 | 12 | India | °57'16.3 | °41'19.3 | 1200m |
| NCBS-BF229 | *Apis* | *dorsata* | Female(worker) | Nyaton Kitnya | 2017 | 11 | 12 | India | °57'16.3 | °41'19.3 | 1200m |
| NCBS-BF230 | *Apis* | *dorsata* | Female(worker) | Nyaton Kitnya | 2017 | 11 | 12 | India | °57'16.3 | °41'19.3 | 1200m |
| NCBS-BF231 | *Apis* | *dorsata* | Female(worker) | Nyaton Kitnya | 2017 | 11 | 12 | India | °57'16.3 | °41'19.3 | 1200m |
| NCBS-BF232 | *Apis* | *dorsata* | Female(worker) | Nyaton Kitnya | 2017 | 10 | 13 | India | °57'13.8 | °39'06.3 | 1000m |
| NCBS-BF233 | *Apis* | *dorsata* | Female(worker) | Nyaton Kitnya | 2017 | 10 | 13 | India | °57'13.8 | °39'06.3 | 1000m |
| NCBS-BF234 | *Apis* | *dorsata* | Female(worker) | Nyaton Kitnya | 2017 | 10 | 13 | India | °57'13.8 | °39'06.3 | 1000m |
| NCBS-BF235 | *Apis* | *dorsata* | Female(worker) | Nyaton Kitnya | 2017 | 10 | 13 | India | °57'13.8 | °39'06.3 | 1000m |
| NCBS-BF236 | *Apis* | *dorsata* | Female(worker) | Nyaton Kitnya | 2017 | 10 | 13 | India | °57'13.8 | °39'06.3 | 1000m |
| NCBS-BF237 | *Apis* | *dorsata* | Female(worker) | Nyaton Kitnya | 2017 | 10 | 13 | India | °57'13.8 | °39'06.3 | 1000m |
| NCBS-BF238 | *Apis* | *dorsata* | Female(worker) | Nyaton Kitnya | 2017 | 10 | 13 | India | °57'13.8 | °39'06.3 | 1000m |
| NCBS-BF239 | *Apis* | *dorsata* | Female(worker) | Prabhudev M.V | 2018 | 1 | 19 | India | °42'15.0' | °44'15.7 | 149m |
| NCBS-BF240 | *Apis* | *dorsata* | Female(worker) | Prabhudev M.V | 2018 | 1 | 19 | India | °42'15.0' | °44'15.7 | 149m |
| NCBS-BF241 | *Apis* | *dorsata* | Female(worker) | Nyaton Kitnya | 2017 | 12 | 4 | India | °08'56.8 | °45'57.5 | 300m |
| NCBS-BF242 | *Apis* | *dorsata* | Female(worker) | Prabhudev M.V | 2017 | 10 | 28 | India | °12'22.8' | °33'40.4 | 1162m |
| NCBS-BF243 | *Apis* | *dorsata* | Female(worker) | Prabhudev M.V | 2017 | 10 | 28 | India | °12'22.8' | °33'40.4 | 1162m |
| NCBS-BF246 | *Apis* | *dorsata* | Female(worker) | Geetha G.T. | 2018 | 2 | 16 | India | °08'46.80 | 31'51.8 | 887m |
| NCBS-BF247 | *Apis* | *dorsata* | Female(worker) | Geetha G.T. | 2018 | 2 | 16 | India | °08'46.80 | 31'51.8 | 887m |
| NCBS-BF118 | *Apis* | *dorsata* | Female(worker) | GU/Uni Würzbu | 2017 | 8 | 28 | India | ###### | 94.991 | 856 |
| NCBS-BF119 | *Apis* | *dorsata* | Female(worker) | GU/Uni Würzbu | 2017 | 8 | 28 | India | ###### | 94.991 | 856 |
| NCBS-BF120 | *Apis* | *dorsata* | Female(worker) | GU/Uni Würzbu | 2016 | 6 | 5 | India | ###### | 94.447 | 1099 |
| NCBS-BF121 | *Apis* | *dorsata* | Female(worker) | GU/Uni Würzbu | 2016 | 6 | 7 | India | ###### | 94.109 | 1940 |
| NCBS-BF122 | *Apis* | *dorsata* | Female(worker) | GU/Uni Würzbu | 2016 | 6 | 7 | India | ###### | 94.109 | 1940 |

| | | | | | | | | | | | |
|---|---|---|---|---|---|---|---|---|---|---|---|
| NCBS-BF123 | *Apis* | *dorsata* | Female(worker) | GU/Uni Würzbu | 2016 | 6 | 8 | India | ###### | 94.229 | 1755 |
| NCBS-BF124 | *Apis* | *dorsata* | Female(worker) | GU/Uni Würzbu | 2016 | 6 | 8 | India | ###### | 94.229 | 1755 |
| NCBS-BF125 | *Apis* | *dorsata* | Female(worker) | GU/Uni Würzbu | 2016 | 6 | 12 | India | ###### | 94.391 | 99 |
| NCBS-BF126 | *Apis* | *dorsata* | Female(worker) | GU/Uni Würzbu | 2016 | 6 | 12 | India | ###### | 94.391 | 99 |
| NCBS-BF127 | *Apis* | *dorsata* | Female(worker) | GU/Uni Würzbu | 2016 | 6 | 10 | India | ###### | 94.63 | 623 |
| NCBS-BF128 | *Apis* | *dorsata* | Female(worker) | GU/Uni Würzbu | 2016 | 6 | 10 | India | ###### | 94.63 | 623 |
| NRC-AA-2861 | *Apis* | *laboriosa* | Female(worker) | Qiu Lifei | 2019 | 4 | 29 | China | 24.158 | 101.43 | 1824m |
| NRC-AA-2862 | *Apis* | *laboriosa* | Female(worker) | Qiu Lifei | 2019 | 4 | 29 | China | 24.158 | 101.43 | 1824m |
| NRC-AA-2863 | *Apis* | *laboriosa* | Female(worker) | Qiu Lifei | 2019 | 4 | 29 | China | 24.158 | 101.43 | 1824m |
| NRC-AA-2864 | *Apis* | *laboriosa* | Female(worker) | Qiu Lifei | 2019 | 4 | 29 | China | 24.158 | 101.43 | 1824m |
| NRC-AA-2865 | *Apis* | *laboriosa* | Female(worker) | Qiu Lifei | 2019 | 4 | 29 | China | 24.158 | 101.43 | 1824m |
| NRC-AA-2866 | *Apis* | *laboriosa* | Female(worker) | Qiu Lifei | 2019 | 4 | 14 | China | 23.925 | 101.59 | 1463m |
| NRC-AA-2867 | *Apis* | *laboriosa* | Female(worker) | Qiu Lifei | 2019 | 4 | 14 | China | 23.925 | 101.59 | 1463m |
| NRC-AA-2868 | *Apis* | *laboriosa* | Female(worker) | Qiu Lifei | 2019 | 4 | 14 | China | 23.925 | 101.59 | 1463m |
| NRC-AA-2869 | *Apis* | *laboriosa* | Female(worker) | Qiu Lifei | 2019 | 4 | 14 | China | 23.925 | 101.59 | 1463m |
| NRC-AA-2870 | *Apis* | *laboriosa* | Female(worker) | Qiu Lifei | 2019 | 4 | 14 | China | 23.925 | 101.59 | 1463m |
| NRC-AA-2871 | *Apis* | *laboriosa* | Female(worker) | Nyaton Kitnya; | 2019 | 3 | 26 | India | 28.155 | 95.182 | 233m |
| NRC-AA-2872 | *Apis* | *laboriosa* | Female(worker) | Nyaton Kitnya; | 2019 | 3 | 26 | India | 28.155 | 95.182 | 233m |
| NRC-AA-2873 | *Apis* | *laboriosa* | Female(worker) | Nyaton Kitnya; | 2019 | 3 | 28 | India | 28.487 | 95.087 | 534m |
| NRC-AA-2874 | *Apis* | *laboriosa* | Female(worker) | Nyaton Kitnya; | 2019 | 3 | 28 | India | 28.487 | 95.087 | 534m |
| NRC-AA-2875 | *Apis* | *laboriosa* | Female(worker) | Nyaton Kitnya | 2018 | 4 | 28 | India | 26.962 | 95.631 | 1060m |
| NRC-AA-2876 | *Apis* | *laboriosa* | Female(worker) | Nyaton Kitnya | 2018 | 4 | 28 | India | 26.962 | 95.631 | 1060m |
| NRC-AA-2877 | *Apis* | *laboriosa* | Female(worker) | Nyaton Kitnya | 2018 | 4 | 28 | India | 26.962 | 95.631 | 1060m |
| NRC-AA-2878 | *Apis* | *laboriosa* | Female(worker) | Nyaton Kitnya | 2018 | 4 | 28 | India | 26.962 | 95.631 | 1060m |
| NRC-AA-2879 | *Apis* | *laboriosa* | Female(worker) | Nyaton Kitnya | 2018 | 4 | 28 | India | 26.962 | 95.631 | 1060m |
| NRC-AA-2880 | *Apis* | *laboriosa* | Female(worker) | Nyaton Kitnya; | 2019 | 4 | 4 | India | 27.207 | 92.482 | 1286m |
| NRC-AA-2881 | *Apis* | *laboriosa* | Female(worker) | Nyaton Kitnya | 2017 | 10 | 28 | India | 27.206 | 92.561 | 1164m |
| NRC-AA-2882 | *Apis* | *laboriosa* | Female(worker) | Nyaton Kitnya | 2017 | 10 | 28 | India | 27.206 | 92.561 | 1164m |
| NRC-AA-2883 | *Apis* | *laboriosa* | Female(worker) | Nyaton Kitnya | 2017 | 10 | 28 | India | 27.206 | 92.561 | 1164m |
| NRC-AA-2884 | *Apis* | *laboriosa* | Female(worker) | Nyaton Kitnya | 2017 | 10 | 28 | India | 27.206 | 92.561 | 1164m |
| NRC-AA-2885 | *Apis* | *laboriosa* | Female(worker) | Nyaton Kitnya | 2017 | 10 | 28 | India | 27.206 | 92.561 | 1164m |
| NRC-AA-2886 | *Apis* | *laboriosa* | Female(worker) | Nyaton Kitnya | 2017 | 10 | 28 | India | 27.206 | 92.561 | 1164m |
| NRC-AA-2887 | *Apis* | *laboriosa* | Female(worker) | Nyaton Kitnya | 2017 | 10 | 28 | India | 27.206 | 92.561 | 1164m |
| NRC-AA-2888 | *Apis* | *laboriosa* | Female(worker) | Nyaton Kitnya | 2021 | 10 | 28 | India | 27.206 | 92.561 | 1164m |
| NRC-AA-2889 | *Apis* | *laboriosa* | Female(worker) | Nyaton Kitnya | 2018 | 4 | 6 | India | 25.579 | 93.868 | 1837m |
| NRC-AA-2890 | *Apis* | *laboriosa* | Female(worker) | Nyaton Kitnya | 2018 | 4 | 6 | India | 25.579 | 93.868 | 1837m |
| NRC-AA-2891 | *Apis* | *laboriosa* | Female(worker) | Nyaton Kitnya | 2018 | 4 | 6 | India | 25.579 | 93.868 | 1837m |
| NRC-AA-2892 | *Apis* | *laboriosa* | Female(worker) | Prabhudev MV | 2019 | 5 | 2 | India | 30.491 | 79.087 | 1008m |
| NRC-AA-2893 | *Apis* | *laboriosa* | Female(worker) | Prabhudev MV | 2019 | 5 | 2 | India | 30.491 | 79.087 | 1008m |
| NRC-AA-2894 | *Apis* | *laboriosa* | Female(worker) | Prabhudev MV | 2019 | 5 | 2 | India | 30.491 | 79.087 | 1008m |
| NRC-AA-2895 | *Apis* | *laboriosa* | Female(worker) | Prabhudev MV | | | | India | 30.018 | 80.57 | 1540m |
| NRC-AA-2896 | *Apis* | *laboriosa* | Female(worker) | Prabhudev MV | | | | India | 30.018 | 80.57 | 1540m |
| NRC-AA-2897 | *Apis* | *laboriosa* | Female(worker) | Prabhudev MV | | | | India | 30.018 | 80.57 | 1540m |
| NRC-AA-2898 | *Apis* | *laboriosa* | Female(worker) | Prabhudev MV | | | | India | 30.018 | 80.57 | 1540m |
| NRC-AA-2899 | *Apis* | *laboriosa* | Female(worker) | Prabhudev MV | | | | India | 30.018 | 80.57 | 1540m |
| NRC-AA-2900 | *Apis* | *laboriosa* | Female(worker) | Prabhudev MV | | | | India | 30.018 | 80.57 | 1540m |
| NRC-AA-2901 | *Apis* | *laboriosa* | Female(worker) | Prabhudev MV | | | | India | 30.018 | 80.57 | 1540m |
| NRC-AA-2902 | *Apis* | *laboriosa* | Female(worker) | Prabhudev MV | | | | India | 30.018 | 80.57 | 1540m |
| NRC-AA-2903 | *Apis* | *laboriosa* | Female(worker) | Thai Hong Pham | 2019 | 4 | 12 | Vietnam | 22.346 | 103.85 | |
| NRC-AA-2904 | *Apis* | *laboriosa* | Female(worker) | Thai Hong Pham | 2019 | 4 | 12 | Vietnam | 22.346 | 103.85 | |
| NRC-AA-2905 | *Apis* | *laboriosa* | Female(worker) | Thai Hong Pham | 2019 | 4 | 12 | Vietnam | 22.346 | 103.85 | |
| NRC-AA-2906 | *Apis* | *laboriosa* | Female(worker) | Thai Hong Pham | 2007 | 11 | 30 | Vietnam | 21.544 | 104.02 | |
| NRC-AA-2907 | *Apis* | *laboriosa* | Female(worker) | Thai & Se | 2001 | 7 | | Vietnam | 20.995 | 104.74 | |
| NRC-AA-2908 | *Apis* | *laboriosa* | Female(worker) | Thai & Se | 2001 | 7 | | Vietnam | 20.995 | 104.74 | |
| NRC-AA-2909 | *Apis* | *laboriosa* | Female(worker) | Thai & Se | 2001 | 7 | | Vietnam | 20.995 | 104.74 | |
| NRC-AA-2910 | *Apis* | *laboriosa* | Male ( drone) | Qiu Lifei | 2019 | 4 | 14 | China | 23.925 | 101.59 | 1463m |
| NRC-AA-2911 | *Apis* | *laboriosa* | Male ( drone) | Qiu Lifei | 2019 | 4 | 14 | China | 23.925 | 101.59 | 1463m |
| NRC-AA-2912 | *Apis* | *laboriosa* | Male ( drone) | Qiu Lifei | 2019 | 4 | 14 | China | 23.925 | 101.59 | 1463m |
| NRC-AA-2913 | *Apis* | *laboriosa* | Male ( drone) | Qiu Lifei | 2019 | 4 | 14 | China | 23.925 | 101.59 | 1463m |
| NRC-AA-2914 | *Apis* | *laboriosa* | Male ( drone) | Nyaton Kitnya | 2018 | 4 | 28 | India | 26.954 | 95.652 | 1060m |

| | | | | | | | | | | |
|---|---|---|---|---|---|---|---|---|---|---|
| NRC-AA-2915 | *Apis laboriosa* | Male ( drone) | Nyaton Kitnya | 2018 | 4 | 28 | India | 26.954 | 95.652 | 1060m |
| NRC-AA-2916 | *Apis laboriosa* | Male ( drone) | Nyaton Kitnya | 2018 | 4 | 28 | India | 26.954 | 95.652 | 1060m |
| NRC-AA-2917 | *Apis laboriosa* | Male ( drone) | Nyaton Kitnya | 2018 | 4 | 28 | India | 26.954 | 95.652 | 1060m |
| NRC-AA-2918 | *Apis laboriosa* | Male ( drone) | B.A. Underwod | 1984 | 5 | 8 | Nepal | 28.267 | 84.017 | 6100 ft |
| NRC-AA-2919 | *Apis laboriosa* | Male ( drone) | Thai Hong Pham | 2019 | 4 | 12 | Vietnam | 22.346 | 103.85 | |
| NRC-AA-2920 | *Apis laboriosa* | Male ( drone) | Thai & Se | 2001 | 7 | | Vietnam | 20.995 | 104.74 | |
| NRC-AA-2921 | *Apis laboriosa* | Male ( drone) | Thai & Se | 2001 | 7 | | Vietnam | 20.995 | 104.74 | |
| NRC-AA-2928 | *Apis laboriosa* | Male ( drone) | Qiu Lifei | 2019 | 4 | 14 | China | 23.925 | 101.59 | 1463m |
| NRC-AA-2958 | *Apis laboriosa* | Male ( drone) | Qiu Lifei | 2019 | 4 | 14 | China | 23.925 | 101.59 | 1463m |
| NRC-AA-3079 | *Apis laboriosa* | Female(worker) | Nyaton Kitnya | 2018 | 4 | 28 | India | 26.962 | 95.631 | 1060m |
| NRC-AA-3080 | *Apis laboriosa* | Female(worker) | Nyaton Kitnya | 2018 | 4 | 28 | India | 26.962 | 95.631 | 1060m |
| NRC-AA-3081 | *Apis laboriosa* | Female(worker) | Nyaton Kitnya | 2018 | 4 | 28 | India | 26.962 | 95.631 | 1060m |
| NRC-AA-3082 | *Apis laboriosa* | Female(worker) | Nyaton Kitnya | 2018 | 4 | 28 | India | 26.962 | 95.631 | 1060m |
| NRC-AA-3083 | *Apis laboriosa* | Female(worker) | Nyaton Kitnya | 2018 | 4 | 28 | India | 26.962 | 95.631 | 1060m |
| NRC-AA-3903 | *Apis laboriosa* | Female(worker) | Nyaton Kitnya | 2017 | 10 | 13 | India | 26.934 N | 95.576 E | 1470m |
| NRC-AA-3904 | *Apis laboriosa* | Female(worker) | Nyaton Kitnya | 2017 | 10 | 13 | India | 26.934 N | 95.576 E | 1470m |
| NRC-AA-3905 | *Apis laboriosa* | Female(worker) | Nyaton Kitnya | 2017 | 10 | 13 | India | 26.934 N | 95.576 E | 1470m |
| NRC-AA-3906 | *Apis laboriosa* | Female(worker) | Nyaton Kitnya | 2017 | 10 | 13 | India | 26.934 N | 95.576 E | 1470m |
| NRC-AA-3907 | *Apis laboriosa* | Female(worker) | Nyaton Kitnya | 2017 | 10 | 13 | India | 26.934 N | 95.576 E | 1470m |
| NRC-AA-3908 | *Apis laboriosa* | Female(worker) | Nyaton Kitnya | 2019 | 3 | 28 | India | 28.487 N | 95.087 E | 534m |
| NRC-AA-3909 | *Apis laboriosa* | Female(worker) | Nyaton Kitnya | 2019 | 3 | 28 | India | 28.487 N | 95.087 E | 534m |
| NRC-AA-3910 | *Apis laboriosa* | Female(worker) | Nyaton Kitnya | 2019 | 3 | 28 | India | 28.487 N | 95.087 E | 534m |
| NRC-AA-3911 | *Apis laboriosa* | Female(worker) | Nyaton Kitnya | 2019 | 3 | 28 | India | 28.487 N | 95.087 E | 534m |
| NRC-AA-3912 | *Apis laboriosa* | Female(worker) | Nyaton Kitnya | 2019 | 3 | 28 | India | 28.487 N | 95.087 E | 534m |
| NRC-AA-3913 | *Apis laboriosa* | Female(worker) | Nyaton Kitnya | 2017 | 10 | 28 | India | 27.206 N | 92.561 E | 1164m |
| NRC-AA-3914 | *Apis laboriosa* | Female(worker) | Nyaton Kitnya | 2017 | 10 | 28 | India | 27.206 N | 92.561 E | 1164m |
| NRC-AA-3915 | *Apis laboriosa* | Female(worker) | Nyaton Kitnya | 2017 | 10 | 28 | India | 27.206 N | 92.561 E | 1164m |
| NRC-AA-3916 | *Apis laboriosa* | Female(worker) | Nyaton Kitnya | 2017 | 10 | 28 | India | 27.206 N | 92.561 E | 1164m |
| NRC-AA-3917 | *Apis laboriosa* | Female(worker) | Nyaton Kitnya | 2017 | 10 | 28 | India | 27.206 N | 92.561 E | 1164m |
| NRC-AA-3918 | *Apis laboriosa* | Female(worker) | Nyaton Kitnya | 2017 | 10 | 28 | India | 27.206 N | 92.561 E | 1164m |
| NRC-AA-3919 | *Apis laboriosa* | Female(worker) | Nyaton Kitnya | 2017 | 10 | 28 | India | 27.206 N | 92.561 E | 1164m |
| NRC-AA-3920 | *Apis laboriosa* | Female(worker) | Nyaton Kitnya | 2017 | 10 | 28 | India | 27.206 N | 92.561 E | 1164m |
| NRC-AA-3921 | *Apis laboriosa* | Female(worker) | Nyaton Kitnya | 2017 | 10 | 28 | India | 27.206 N | 92.561 E | 1164m |
| NRC-AA-3922 | *Apis laboriosa* | Female(worker) | Nyaton Kitnya | 2019 | 5 | 16 | India | 28.456 N | 94.684 E | 356m |
| NRC-AA-3923 | *Apis laboriosa* | Female(worker) | Nyaton Kitnya | 2019 | 5 | 16 | India | 28.456 N | 94.684 E | 356m |
| NRC-AA-3924 | *Apis laboriosa* | Female(worker) | Nyaton Kitnya | 2019 | 5 | 16 | India | 28.456 N | 94.684 E | 356m |
| NRC-AA-3925 | *Apis laboriosa* | Female(worker) | Nyaton Kitnya | 2019 | 5 | 16 | India | 28.456 N | 94.684 E | 356m |
| NRC-AA-3926 | *Apis laboriosa* | Female(worker) | Nyaton Kitnya | 2019 | 5 | 16 | India | 28.456 N | 94.684 E | 356m |
| NRC-AA-3927 | *Apis laboriosa* | Female(worker) | Nyaton Kitnya | 2019 | 5 | 16 | India | 28.456 N | 94.684 E | 356m |
| NRC-AA-3928 | *Apis laboriosa* | Female(worker) | Nyaton Kitnya | 2019 | 5 | 16 | India | 28.456 N | 94.684 E | 356m |
| NRC-AA-3929 | *Apis laboriosa* | Female(worker) | Nyaton Kitnya | 2019 | 5 | 16 | India | 28.456 N | 94.684 E | 356m |
| NRC-AA-3930 | *Apis laboriosa* | Female(worker) | Nyaton Kitnya | 2019 | 5 | 16 | India | 28.456 N | 94.684 E | 356m |
| NRC-AA-3931 | *Apis laboriosa* | Female(worker) | Nyaton Kitnya | 2019 | 5 | 16 | India | 28.456 N | 94.684 E | 356m |
| NRC-AA-3932 | *Apis laboriosa* | Female(worker) | Nyaton Kitnya | 2018 | 4 | 28 | India | 26.962 N | 95.631 E | 1060m |
| NRC-AA-3933 | *Apis laboriosa* | Female(worker) | Nyaton Kitnya | 2018 | 4 | 28 | India | 26.962 N | 95.631 E | 1060m |
| NRC-AA-3934 | *Apis laboriosa* | Female(worker) | Nyaton Kitnya | 2018 | 4 | 28 | India | 26.962 N | 95.631 E | 1060m |
| NRC-AA-3935 | *Apis laboriosa* | Female(worker) | Nyaton Kitnya | 2018 | 4 | 28 | India | 26.962 N | 95.631 E | 1060m |
| NRC-AA-3936 | *Apis laboriosa* | Female(worker) | Nyaton Kitnya | 2018 | 4 | 28 | India | 26.962 N | 95.631 E | 1060m |
| NRC-AA-3937 | *Apis laboriosa* | Female(worker) | Nyaton Kitnya | 2018 | 4 | 28 | India | 26.962 N | 95.631 E | 1060m |
| NRC-AA-3938 | *Apis laboriosa* | Female(worker) | Nyaton Kitnya | 2018 | 4 | 28 | India | 26.962 N | 95.631 E | 1060m |
| NRC-AA-3939 | *Apis laboriosa* | Female(worker) | Nyaton Kitnya | 2018 | 4 | 28 | India | 26.962 N | 95.631 E | 1060m |
| NRC-AA-3940 | *Apis laboriosa* | Female(worker) | Nyaton Kitnya | 2018 | 4 | 28 | India | 26.962 N | 95.631 E | 1060m |
| NRC-AA-3941 | *Apis laboriosa* | Female(worker) | Nyaton Kitnya | 2018 | 4 | 28 | India | 26.962 N | 95.631 E | 1060m |
| NRC-AA-3942 | *Apis laboriosa* | Female(worker) | Nyaton Kitnya | 2018 | 7 | 5 | India | 27.580 N | 91.870 E | 2940m |
| NRC-AA-3943 | *Apis laboriosa* | Female(worker) | Nyaton Kitnya | 2018 | 7 | 5 | India | 27.580 N | 91.870 E | 2940m |
| NRC-AA-3944 | *Apis laboriosa* | Female(worker) | Nyaton Kitnya | 2018 | 7 | 5 | India | 27.580 N | 91.870 E | 2940m |
| NRC-AA-3945 | *Apis laboriosa* | Female(worker) | Nyaton Kitnya | 2018 | 7 | 5 | India | 27.580 N | 91.870 E | 2940m |
| NRC-AA-3946 | *Apis laboriosa* | Female(worker) | Nyaton Kitnya | 2018 | 7 | 5 | India | 27.580 N | 91.870 E | 2940m |
| NRC-AA-3947 | *Apis laboriosa* | Female(worker) | Nyaton Kitnya | 2018 | 7 | 5 | India | 27.580 N | 91.870 E | 2940m |
| NRC-AA-3948 | *Apis laboriosa* | Female(worker) | Nyaton Kitnya | 2018 | 7 | 5 | India | 27.580 N | 91.870 E | 2940m |
| NRC-AA-3949 | *Apis laboriosa* | Female(worker) | Nyaton Kitnya | 2018 | 7 | 5 | India | 27.580 N | 91.870 E | 2940m |

| | | | | | | | | | | |
|---|---|---|---|---|---|---|---|---|---|---|
| NRC-AA-3950 | *Apis laboriosa* | Female(worker) | Nyaton Kitnya | 2018 | 7 | 5 | India | 27.580 N | 91.870 E | 2940m |
| NRC-AA-3951 | *Apis laboriosa* | Female(worker) | Nyaton Kitnya | 2018 | 7 | 5 | India | 27.580 N | 91.870 E | 2940m |
| NRC-AA-3952 | *Apis laboriosa* | Female(worker) | Nyaton Kitnya | 2018 | 7 | 5 | India | 27.570 N | 91.850 E | 2320m |
| NRC-AA-3953 | *Apis laboriosa* | Female(worker) | Nyaton Kitnya | 2018 | 7 | 5 | India | 27.570 N | 91.850 E | 2320m |
| NRC-AA-3954 | *Apis laboriosa* | Female(worker) | Nyaton Kitnya | 2018 | 7 | 5 | India | 27.570 N | 91.850 E | 2320m |
| NRC-AA-3955 | *Apis laboriosa* | Female(worker) | Nyaton Kitnya | 2018 | 7 | 5 | India | 27.570 N | 91.850 E | 2320m |
| NRC-AA-3956 | *Apis laboriosa* | Female(worker) | Nyaton Kitnya | 2018 | 7 | 5 | India | 27.570 N | 91.850 E | 2320m |
| NRC-AA-3957 | *Apis laboriosa* | Female(worker) | Nyaton Kitnya | 2018 | 7 | 5 | India | 27.570 N | 91.850 E | 2320m |
| NRC-AA-3958 | *Apis laboriosa* | Female(worker) | Nyaton Kitnya | 2018 | 7 | 5 | India | 27.570 N | 91.850 E | 2320m |
| NRC-AA-3959 | *Apis laboriosa* | Female(worker) | Nyaton Kitnya | 2018 | 7 | 5 | India | 27.570 N | 91.850 E | 2320m |
| NRC-AA-7057 | *Apis laboriosa* | Male (drone) | itnya, Anne-Sop | 2023 | 6 | 10 | India | 26.954 | 95.652 | 1060m |
| NRC-AA-7058 | *Apis laboriosa* | Male (drone) | itnya, Anne-Sop | 2023 | 6 | 10 | India | 26.954 | 95.652 | 1060m |
| NRC-AA-7059 | *Apis laboriosa* | Male (drone) | itnya, Anne-Sop | 2023 | 6 | 10 | India | 26.954 | 95.652 | 1060m |
| NRC-AA-7060 | *Apis laboriosa* | Male (drone) | itnya, Anne-Sop | 2023 | 6 | 10 | India | 26.954 | 95.652 | 1060m |
| NRC-AA-7061 | *Apis laboriosa* | Male (drone) | itnya, Anne-Sop | 2023 | 6 | 10 | India | 26.954 | 95.652 | 1060m |
| NRC-AA-7062 | *Apis laboriosa* | Male (drone) | itnya, Anne-Sop | 2023 | 6 | 10 | India | 26.954 | 95.652 | 1060m |
| NRC-AA-7063 | *Apis laboriosa* | Male (drone) | itnya, Anne-Sop | 2023 | 6 | 10 | India | 26.954 | 95.652 | 1060m |
| NRC-AA-7064 | *Apis laboriosa* | Female(worker) | itnya, Anne-Sop | 2023 | 6 | 10 | India | 26.954 | 95.652 | 1060m |
| NRC-AA-7065 | *Apis laboriosa* | Female(worker) | itnya, Anne-Sop | 2023 | 6 | 10 | India | 26.954 | 95.652 | 1060m |
| NRC-AA-7066 | *Apis laboriosa* | Female(worker) | itnya, Anne-Sop | 2023 | 6 | 10 | India | 26.954 | 95.652 | 1060m |
| NRC-AA-7067 | *Apis laboriosa* | Female(worker) | itnya, Anne-Sop | 2023 | 6 | 10 | India | 26.954 | 95.652 | 1060m |
| NRC-AA-7068 | *Apis laboriosa* | Female(worker) | itnya, Anne-Sop | 2023 | 6 | 10 | India | 26.954 | 95.652 | 1060m |
| NRC-AA-7069 | *Apis laboriosa* | Female(worker) | itnya, Anne-Sop | 2023 | 6 | 10 | India | 26.954 | 95.652 | 1060m |
| NRC-AA-7070 | *Apis laboriosa* | Female(worker) | itnya, Anne-Sop | 2023 | 6 | 10 | India | 26.954 | 95.652 | 1060m |
| NRC-AA-7071 | *Apis laboriosa* | Female(worker) | itnya, Anne-Sop | 2023 | 6 | 10 | India | 26.954 | 95.652 | 1060m |
| NRC-AA-7072 | *Apis laboriosa* | Female(worker) | itnya, Anne-Sop | 2023 | 6 | 10 | India | 26.954 | 95.652 | 1060m |
| NRC-AA-7162 | *Apis laboriosa* | Female(worker) | Nyaton Kitnya | 2018 | 4 | 28 | India | 26.962 | 95.631 | 1060m |
| NRC-AA-7163 | *Apis laboriosa* | Female(worker) | Nyaton Kitnya | 2018 | 4 | 28 | India | 26.962 | 95.631 | 1060m |
| NRC-AA-7164 | *Apis laboriosa* | Female(worker) | Nyaton Kitnya | 2018 | 4 | 28 | India | 26.962 | 95.631 | 1060m |
| NRC-AA-7165 | *Apis laboriosa* | Female(worker) | Nyaton Kitnya | 2018 | 4 | 28 | India | 26.962 | 95.631 | 1060m |
| NRC-AA-7166 | *Apis laboriosa* | Female(worker) | Nyaton Kitnya | 2018 | 4 | 28 | India | 26.962 | 95.631 | 1060m |
| NRC-AA-7265 | *Apis laboriosa* | Female(worker) | Nyaton Kitnya | 2018 | 4 | 7 | India | 27.577 | 91.875 | 2585m |
| NRC-AA-7266 | *Apis laboriosa* | Female(worker) | Nyaton Kitnya | 2018 | 4 | 7 | India | 27.577 | 91.875 | 2585m |
| NRC-AA-7267 | *Apis laboriosa* | Female(worker) | Nyaton Kitnya | 2018 | 4 | 7 | India | 27.577 | 91.875 | 2585m |
| NRC-AA-7268 | *Apis laboriosa* | Female(worker) | Nyaton Kitnya | 2018 | 4 | 7 | India | 27.577 | 91.875 | 2585m |
| NRC-AA-7269 | *Apis laboriosa* | Female(worker) | Nyaton Kitnya | 2018 | 4 | 7 | India | 27.577 | 91.875 | 2585m |
| NRC-AA-7270 | *Apis laboriosa* | Female(worker) | Nyaton Kitnya | 2018 | 4 | 7 | India | 27.577 | 91.875 | 2585m |
| NRC-AA-7271 | *Apis laboriosa* | Female(worker) | Nyaton Kitnya | 2018 | 4 | 7 | India | 27.577 | 91.875 | 2585m |
| NRC-AA-7272 | *Apis laboriosa* | Female(worker) | Nyaton Kitnya | 2018 | 4 | 7 | India | 27.577 | 91.875 | 2585m |
| NRC-AA-7273 | *Apis laboriosa* | Female(worker) | Nyaton Kitnya | 2018 | 4 | 7 | India | 27.577 | 91.875 | 2585m |
| NRC-AA-7274 | *Apis laboriosa* | Female(worker) | Nyaton Kitnya | 2018 | 4 | 7 | India | 27.577 | 91.875 | 2585m |
| NRC-AA-7275 | *Apis laboriosa* | Female(worker) | Nyaton Kitnya | 2018 | 4 | 7 | India | 27.577 | 91.875 | 2585m |
| NRC-AA-7276 | *Apis laboriosa* | Female(worker) | Nyaton Kitnya | 2018 | 4 | 7 | India | 27.577 | 91.875 | 2585m |
| NRC-AA-7277 | *Apis laboriosa* | Female(worker) | Prabhudev MV | 2019 | 5 | 2 | India | 30.458 N | 79.275 E | 1550m |
| NRC-AA-7278 | *Apis laboriosa* | Female(worker) | Prabhudev MV | 2019 | 5 | 2 | India | 30.458 N | 79.275 E | 1550m |
| NRC-AA-7279 | *Apis laboriosa* | Female(worker) | Prabhudev MV | 2019 | 5 | 2 | India | 30.458 N | 79.275 E | 1550m |
| NRC-AA-7280 | *Apis laboriosa* | Female(worker) | Prabhudev MV | 2019 | 5 | 2 | India | 30.458 N | 79.275 E | 1550m |
| NRC-AA-7281 | *Apis laboriosa* | Female(worker) | Prabhudev MV | 2019 | 5 | 2 | India | 30.458 N | 79.275 E | 1550m |
| NRC-AA-7282 | *Apis laboriosa* | Female(worker) | Prabhudev MV | 2019 | 5 | 2 | India | 30.458 N | 79.275 E | 1550m |
| NRC-AA-7283 | *Apis laboriosa* | Female(worker) | Prabhudev MV | 2019 | 5 | 2 | India | 30.458 N | 79.275 E | 1550m |
| NRC-AA-7284 | *Apis laboriosa* | Female(worker) | Prabhudev MV | 2019 | 5 | 2 | India | 30.458 N | 79.275 E | 1550m |
| NRC-AA-7285 | *Apis laboriosa* | Female(worker) | Prabhudev MV | 2019 | 5 | 2 | India | 30.458 N | 79.275 E | 1550m |
| NRC-AA-7286 | *Apis laboriosa* | Female(worker) | Nyaton Kitnya | 2019 | 5 | 16 | India | 28.456 N | 94.684 E | 356m |
| NRC-AA-7476 | *Apis laboriosa* | Female(worker) | Prabhudev MV | 2018 | 5 | 31 | India | 30.086 | 80.259 | 1500m |

| | | | | | | | | | | |
|---|---|---|---|---|---|---|---|---|---|---|
| NRC-AA-7477 | *Apis laboriosa* | Female(worker) | Prabhudev MV | 2018 | 5 | 31 | India | 30.086 | 80.259 | 1500m |
| NRC-AA-7478 | *Apis laboriosa* | Female(worker) | Prabhudev MV | 2018 | 5 | 31 | India | 30.086 | 80.259 | 1500m |
| NRC-AA-7479 | *Apis laboriosa* | Female(worker) | Prabhudev MV | 2018 | 5 | 31 | India | 30.086 | 80.259 | 1500m |
| NRC-AA-7480 | *Apis laboriosa* | Female(worker) | Prabhudev MV | 2018 | 5 | 31 | India | 30.086 | 80.259 | 1500m |
| NRC-AA-7481 | *Apis laboriosa* | Female(worker) | Prabhudev MV | 2018 | 5 | 31 | India | 30.086 | 80.259 | 1500m |
| NRC-AA-7482 | *Apis laboriosa* | Female(worker) | Prabhudev MV | 2018 | 5 | 31 | India | 30.086 | 80.259 | 1500m |
| NRC-AA-7483 | *Apis laboriosa* | Female(worker) | Prabhudev MV | 2018 | 5 | 31 | India | 30.086 | 80.259 | 1500m |
| NRC-AA-7484 | *Apis laboriosa* | Female(worker) | Prabhudev MV | 2018 | 5 | 31 | India | 30.086 | 80.259 | 1500m |
| NRC-AA-7485 | *Apis laboriosa* | Female(worker) | Prabhudev MV | 2018 | 5 | 31 | India | 30.086 | 80.259 | 1500m |
| NRC-AA-7486 | *Apis laboriosa* | Female(worker) | Prabhudev MV | 2019 | 5 | 2 | India | °29'8.13" | 17'31.15 | 2250m |
| NRC-AA-7487 | *Apis laboriosa* | Female(worker) | Prabhudev MV | 2019 | 5 | 2 | India | °29'8.13" | 17'31.15 | 2250m |
| NRC-AA-7488 | *Apis laboriosa* | Female(worker) | Prabhudev MV | 2019 | 5 | 2 | India | °29'8.13" | 17'31.15 | 2250m |
| NRC-AA-7489 | *Apis laboriosa* | Female(worker) | Prabhudev MV | 2019 | 5 | 2 | India | °29'8.13" | 17'31.15 | 2250m |
| NRC-AA-7490 | *Apis laboriosa* | Female(worker) | Prabhudev MV | 2019 | 5 | 2 | India | °29'8.13" | 17'31.15 | 2250m |
| NRC-AA-7491 | *Apis laboriosa* | Female(worker) | Prabhudev MV | 2019 | 5 | 2 | India | °29'8.13" | 17'31.15 | 2250m |
| NRC-AA-7492 | *Apis laboriosa* | Female(worker) | Prabhudev MV | 2019 | 5 | 2 | India | °29'8.13" | 17'31.15 | 2250m |
| NRC-AA-7493 | *Apis laboriosa* | Female(worker) | Prabhudev MV | 2019 | 5 | 2 | India | °29'8.13" | 17'31.15 | 2250m |
| NRC-AA-7494 | *Apis laboriosa* | Female(worker) | Prabhudev MV | 2019 | 5 | 2 | India | °29'8.13" | 17'31.15 | 2250m |
| NRC-AA-7495 | *Apis laboriosa* | Female(worker) | Prabhudev MV | 2019 | 5 | 2 | India | °29'8.13" | 17'31.15 | 2250m |
| NRC-AA-7496 | *Apis laboriosa* | Female(worker) | Prabhudev MV | 2019 | 5 | 2 | India | 30.491 | 79.087 | 1008m |
| NRC-AA-7497 | *Apis laboriosa* | Female(worker) | Prabhudev MV | 2019 | 5 | 2 | India | 30.491 | 79.087 | 1008m |
| NRC-AA-7498 | *Apis laboriosa* | Female(worker) | Prabhudev MV | 2019 | 5 | 2 | India | 30.491 | 79.087 | 1008m |
| NRC-AA-7499 | *Apis laboriosa* | Female(worker) | Prabhudev MV | 2019 | 5 | 2 | India | 30.491 | 79.087 | 1008m |
| NRC-AA-7500 | *Apis laboriosa* | Female(worker) | Prabhudev MV | 2019 | 5 | 2 | India | 30.491 | 79.087 | 1008m |
| NRC-AA-7501 | *Apis laboriosa* | Female(worker) | Prabhudev MV | 2019 | 5 | 2 | India | 30.491 | 79.087 | 1008m |
| NRC-AA-7502 | *Apis laboriosa* | Female(worker) | Prabhudev MV | 2019 | 5 | 2 | India | 30.491 | 79.087 | 1008m |
| NRC-AA-7503 | *Apis laboriosa* | Female(worker) | Prabhudev MV | 2019 | 5 | 2 | India | 30.491 | 79.087 | 1008m |
| NRC-AA-7504 | *Apis laboriosa* | Female(worker) | Prabhudev MV | 2019 | 5 | 2 | India | 30.491 | 79.087 | 1008m |
| NRC-AA-7505 | *Apis laboriosa* | Female(worker) | Prabhudev MV | 2019 | 5 | 2 | India | 30.491 | 79.087 | 1008m |
| NRC-AA-7506 | *Apis laboriosa* | Female(worker) | Prabhudev MV | 2019 | 5 | 2 | India | 30.458 N | 79.275 E | 1550m |
| NRC-AA-7507 | *Apis laboriosa* | Female(worker) | Prabhudev MV | 2019 | 5 | 2 | India | 30.458 N | 79.275 E | 1550m |
| NRC-AA-7508 | *Apis laboriosa* | Female(worker) | Prabhudev MV | 2019 | 5 | 2 | India | 30.458 N | 79.275 E | 1550m |
| NRC-AA-7509 | *Apis laboriosa* | Female(worker) | Prabhudev MV | 2019 | 5 | 2 | India | 30.458 N | 79.275 E | 1550m |
| NRC-AA-7510 | *Apis laboriosa* | Female(worker) | Prabhudev MV | 2019 | 5 | 2 | India | 30.458 N | 79.275 E | 1550m |
| NRC-AA-7511 | *Apis laboriosa* | Female(worker) | Prabhudev MV | 2019 | 5 | 2 | India | 30.458 N | 79.275 E | 1550m |
| NRC-AA-7512 | *Apis laboriosa* | Female(worker) | Prabhudev MV | 2019 | 5 | 2 | India | 30.458 N | 79.275 E | 1550m |
| NRC-AA-7513 | *Apis laboriosa* | Female(worker) | Prabhudev MV | 2019 | 5 | 2 | India | 30.458 N | 79.275 E | 1550m |
| NRC-AA-7514 | *Apis laboriosa* | Female(worker) | Prabhudev MV | 2019 | 5 | 2 | India | 30.458 N | 79.275 E | 1550m |
| NRC-AA-7515 | *Apis laboriosa* | Female(worker) | Prabhudev MV | 2019 | 5 | 2 | India | 30.458 N | 79.275 E | 1550m |
| NRC-AA-7516 | *Apis laboriosa* | Female(worker) | Prabhudev MV | 2018 | 6 | 30 | India | 30.018 N | 30.018 E | 1536m |
| NRC-AA-7517 | *Apis laboriosa* | Female(worker) | Prabhudev MV | 2018 | 6 | 30 | India | 30.018 N | 30.018 E | 1536m |
| NRC-AA-7518 | *Apis laboriosa* | Female(worker) | Prabhudev MV | 2018 | 6 | 30 | India | 30.018 N | 30.018 E | 1536m |
| NRC-AA-7519 | *Apis laboriosa* | Female(worker) | Prabhudev MV | 2018 | 6 | 30 | India | 30.018 N | 30.018 E | 1536m |
| NRC-AA-7520 | *Apis laboriosa* | Female(worker) | Prabhudev MV | 2018 | 6 | 30 | India | 30.018 N | 30.018 E | 1536m |
| NRC-AA-7521 | *Apis laboriosa* | Female(worker) | Prabhudev MV | 2018 | 6 | 30 | India | 30.018 N | 30.018 E | 1536m |
| NRC-AA-7522 | *Apis laboriosa* | Female(worker) | Nyaton Kitnya | 2018 | 4 | 6 | India | 25.579 | 93.868 | 1837m |
| NRC-AA-7523 | *Apis laboriosa* | Female(worker) | Nyaton Kitnya | 2018 | 4 | 6 | India | 25.579 | 93.868 | 1837m |
| NRC-AA-7524 | *Apis laboriosa* | Female(worker) | Nyaton Kitnya | 2018 | 4 | 6 | India | 25.579 | 93.868 | 1837m |
| NRC-AA-7525 | *Apis laboriosa* | Female(worker) | Nyaton Kitnya | 2018 | 4 | 6 | India | 25.579 | 93.868 | 1837m |
| NRC-AA-7526 | *Apis laboriosa* | Female(worker) | Nyaton Kitnya | 2018 | 4 | 6 | India | 25.579 | 93.868 | 1837m |
| NRC-AA-7527 | *Apis laboriosa* | Female(worker) | Nyaton Kitnya | 2018 | 4 | 6 | India | 25.579 | 93.868 | 1837m |
| NRC-AA-7528 | *Apis laboriosa* | Female(worker) | Nyaton Kitnya | 2018 | 4 | 6 | India | 25.579 | 93.868 | 1837m |

| | | | | | | | | | | |
|---|---|---|---|---|---|---|---|---|---|---|
| NRC-AA-7529 | *Apis laboriosa* | Female(worker) | Nyaton Kitnya | 2018 | 4 | 6 | India | 25.579 | 93.868 | 1837m |
| NRC-AA-7530 | *Apis laboriosa* | Female(worker) | Nyaton Kitnya | 2018 | 4 | 6 | India | 25.579 | 93.868 | 1837m |
| NRC-AA-7531 | *Apis laboriosa* | Female(worker) | Nyaton Kitnya | 2018 | 4 | 6 | India | 25.579 | 93.868 | 1837m |
| NRC-AA-7532 | *Apis laboriosa* | Female(worker) | Nyaton Kitnya | 2018 | 4 | 6 | India | 25.564 N | 93.861 E | 1674m |
| NRC-AA-7533 | *Apis laboriosa* | Female(worker) | Nyaton Kitnya | 2018 | 4 | 6 | India | 25.564 N | 93.861 E | 1674m |
| NRC-AA-7534 | *Apis laboriosa* | Female(worker) | Nyaton Kitnya | 2018 | 4 | 6 | India | 25.564 N | 93.861 E | 1674m |
| NRC-AA-7535 | *Apis laboriosa* | Female(worker) | Nyaton Kitnya | 2018 | 4 | 6 | India | 25.564 N | 93.861 E | 1674m |
| NRC-AA-7536 | *Apis laboriosa* | Female(worker) | Nyaton Kitnya | 2018 | 4 | 6 | India | 25.564 N | 93.861 E | 1674m |
| NRC-AA-7537 | *Apis laboriosa* | Female(worker) | Nyaton Kitnya | 2018 | 4 | 6 | India | 25.564 N | 93.861 E | 1674m |
| NRC-AA-7538 | *Apis laboriosa* | Female(worker) | Nyaton Kitnya | 2018 | 4 | 6 | India | 25.564 N | 93.861 E | 1674m |
| NRC-AA-7539 | *Apis laboriosa* | Female(worker) | Nyaton Kitnya | 2018 | 4 | 6 | India | 25.564 N | 93.861 E | 1674m |
| NRC-AA-7540 | *Apis laboriosa* | Female(worker) | Nyaton Kitnya | 2018 | 4 | 6 | India | 25.564 N | 93.861 E | 1674m |
| NRC-AA-7541 | *Apis laboriosa* | Female(worker) | Nyaton Kitnya | 2018 | 4 | 6 | India | 25.564 N | 93.861 E | 1674m |
| NRC-AA-7542 | *Apis laboriosa* | Female(worker) | Nyaton Kitnya | 2017 | 10 | 13 | India | 26.934 N | 95.576 E | 1470m |
| NRC-AA-7543 | *Apis laboriosa* | Female(worker) | Nyaton Kitnya | 2017 | 10 | 13 | India | 26.934 N | 95.576 E | 1470m |
| NRC-AA-7544 | *Apis laboriosa* | Female(worker) | Nyaton Kitnya | 2017 | 10 | 13 | India | 26.934 N | 95.576 E | 1470m |
| NRC-AA-7545 | *Apis laboriosa* | Female(worker) | Nyaton Kitnya | 2017 | 10 | 13 | India | 26.934 N | 95.576 E | 1470m |
| NRC-AA-7546 | *Apis laboriosa* | Female(worker) | Nyaton Kitnya | 2017 | 10 | 13 | India | 26.934 N | 95.576 E | 1470m |
| NRC-AA-7547 | *Apis laboriosa* | Female(worker) | KarsingMegu | 2018 | 5 | 18 | India | 27.885 N | 6.81783 | 1242m |
| NRC-AA-7548 | *Apis laboriosa* | Female(worker) | KarsingMegu | 2018 | 5 | 18 | India | 7.978576 | 96.395 | 485m |
| NRC-AA-7549 | *Apis laboriosa* | Female(worker) | KarsingMegu | 2018 | 5 | 18 | India | 7.978576 | 96.395 | 485m |
| NRC-AA-7550 | *Apis laboriosa* | Female(worker) | KarsingMegu | 2018 | 5 | 18 | India | 7.978576 | 96.395 | 485m |
| NRC-AA-7551 | *Apis laboriosa* | Female(worker) | KarsingMegu | 2018 | 5 | 18 | India | 7.978576 | 96.395 | 485m |
| NRC-AA-7552 | *Apis laboriosa* | Female(worker) | Nyaton Kitnya | 2018 | 7 | 6 | India | 27'05.54 | 92'10.90 | 1978M |
| NRC-AA-7553 | *Apis laboriosa* | Female(worker) | Nyaton Kitnya | 2018 | 7 | 6 | India | 27'05.54 | 92'10.90 | 1978M |
| NRC-AA-7554 | *Apis laboriosa* | Female(worker) | Nyaton Kitnya | 2018 | 7 | 6 | India | 27'05.54 | 92'10.90 | 1978M |
| NRC-AA-7555 | *Apis laboriosa* | Female(worker) | Nyaton Kitnya | 2018 | 7 | 6 | India | 27'05.54 | 92'10.90 | 1978M |
| NRC-AA-7556 | *Apis laboriosa* | Female(worker) | Nyaton Kitnya | 2018 | 7 | 6 | India | 27'05.54 | 92'10.90 | 1978M |
| NRC-AA-7557 | *Apis laboriosa* | Female(worker) | Nyaton Kitnya | 2019 | 4 | 4 | India | 27.207 | 92.482 | 1286m |
| NRC-AA-7558 | *Apis laboriosa* | Female(worker) | Nyaton Kitnya | 2019 | 4 | 4 | India | 27.207 | 92.482 | 1286m |
| NRC-AA-7559 | *Apis laboriosa* | Female(worker) | Nyaton Kitnya | 2019 | 4 | 4 | India | 27.207 | 92.482 | 1286m |
| NRC-AA-7560 | *Apis laboriosa* | Female(worker) | Nyaton Kitnya | 2019 | 4 | 4 | India | 27.207 | 92.482 | 1286m |
| NRC-AA-7561 | *Apis laboriosa* | Female(worker) | Nyaton Kitnya | 2019 | 4 | 4 | India | 27.207 | 92.482 | 1286m |
| NRC-AA-7562 | *Apis laboriosa* | Female(worker) | Nyaton Kitnya | 2019 | 4 | 4 | India | 27.207 | 92.482 | 1286m |
| NRC-AA-7563 | *Apis laboriosa* | Female(worker) | Nyaton Kitnya | 2019 | 4 | 4 | India | 27.207 | 92.482 | 1286m |
| NRC-AA-7564 | *Apis laboriosa* | Female(worker) | Nyaton Kitnya | 2019 | 4 | 4 | India | 27.207 | 92.482 | 1286m |
| NRC-AA-7565 | *Apis laboriosa* | Female(worker) | Nyaton Kitnya | 2019 | 4 | 4 | India | 27.207 | 92.482 | 1286m |
| NRC-AA-7566 | *Apis laboriosa* | Female(worker) | Nyaton Kitnya | 2019 | 4 | 4 | India | 27.207 | 92.482 | 1286m |
| NRC-AA-7567 | *Apis laboriosa* | Female(worker) | Nyaton Kitnya | 2017 | 10 | 6 | India | 27.342 N | 92.273 E | 1500m |
| NRC-AA-7568 | *Apis laboriosa* | Female(worker) | Nyaton Kitnya | 2017 | 10 | 6 | India | 27.342 N | 92.273 E | 1500m |
| NRC-AA-7569 | *Apis laboriosa* | Female(worker) | Nyaton Kitnya | 2017 | 10 | 6 | India | 27.342 N | 92.273 E | 1500m |
| NRC-AA-7570 | *Apis laboriosa* | Female(worker) | Nyaton Kitnya | 2017 | 10 | 6 | India | 27.342 N | 92.273 E | 1500m |
| NRC-AA-7571 | *Apis laboriosa* | Female(worker) | Nyaton Kitnya | 2018 | 7 | 6 | India | 27.405 N | 92.134 E | 1986m |
| NRC-AA-7572 | *Apis laboriosa* | Female(worker) | Nyaton Kitnya | 2018 | 7 | 6 | India | 27.405 N | 92.134 E | 1986m |
| NRC-AA-7573 | *Apis laboriosa* | Female(worker) | Nyaton Kitnya | 2018 | 7 | 6 | India | 27.405 N | 92.134 E | 1986m |
| NRC-AA-7574 | *Apis laboriosa* | Female(worker) | Nyaton Kitnya | 2018 | 4 | 7 | India | 27.577 | 91.875 | 2585m |
| NRC-AA-7575 | *Apis laboriosa* | Female(worker) | Nyaton Kitnya | 2018 | 4 | 7 | India | 27.577 | 91.875 | 2585m |
| NRC-AA-7576 | *Apis laboriosa* | Female(worker) | Nyaton Kitnya | 2018 | 4 | 7 | India | 27.577 | 91.875 | 2585m |
| NRC-AA-7577 | *Apis laboriosa* | Female(worker) | Nyaton Kitnya | 2018 | 4 | 7 | India | 27.577 | 91.875 | 2585m |
| NRC-AA-7578 | *Apis laboriosa* | Female(worker) | Nyaton Kitnya | 2018 | 4 | 7 | India | 27.577 | 91.875 | 2585m |
| NRC-AA-7579 | *Apis laboriosa* | Female(worker) | Nyaton Kitnya | 2018 | 4 | 7 | India | 27.577 | 91.875 | 2585m |
| NRC-AA-7580 | *Apis laboriosa* | Female(worker) | Nyaton Kitnya | 2018 | 4 | 7 | India | 27.577 | 91.875 | 2585m |

| | | | | | | | | | | |
|---|---|---|---|---|---|---|---|---|---|---|
| NRC-AA-7581 | *Apis laboriosa* | Female(worker) | Nyaton Kitnya | 2018 | 4 | 7 | India | 27.577 | 91.875 | 2585m |
| NRC-AA-7582 | *Apis laboriosa* | Female(worker) | Nyaton Kitnya | 2018 | 4 | 7 | India | 27.577 | 91.875 | 2585m |
| NRC-AA-7583 | *Apis laboriosa* | Female(worker) | Nyaton Kitnya | 2018 | 4 | 7 | India | 27.577 | 91.875 | 2585m |
| NRC-AA-7584 | *Apis laboriosa* | Female(worker) | Nyaton Kitnya | 2018 | 4 | 7 | India | 27.577 | 91.875 | 2585m |
| NRC-AA-7585 | *Apis laboriosa* | Female(worker) | Nyaton Kitnya | 2018 | 4 | 7 | India | 27.577 | 91.875 | 2585m |
| NRC-AA-7586 | *Apis laboriosa* | Female(worker) | Nyaton Kitnya | 2018 | 4 | 7 | India | 27.577 | 91.875 | 2585m |
| NRC-AA-7587 | *Apis laboriosa* | Female(worker) | Nyaton Kitnya | 2018 | 4 | 7 | India | 27.577 | 91.875 | 2585m |
| NRC-AA-7588 | *Apis laboriosa* | Female(worker) | Nyaton Kitnya | 2018 | 4 | 7 | India | 27.577 | 91.875 | 2585m |
| NRC-AA-7589 | *Apis laboriosa* | Female(worker) | Nyaton Kitnya | 2018 | 7 | 3-5 | India | 27.591 N | 91.876 E | 2930m |
| NRC-AA-7590 | *Apis laboriosa* | Female(worker) | Nyaton Kitnya | 2018 | 7 | 3-5 | India | 27.591 N | 91.876 E | 2930m |
| NRC-AA-7591 | *Apis laboriosa* | Female(worker) | Nyaton Kitnya | 2018 | 7 | 3-5 | India | 27.591 N | 91.876 E | 2930m |
| NRC-AA-7592 | *Apis laboriosa* | Female(worker) | Nyaton Kitnya | 2018 | 7 | 3-5 | India | 27.591 N | 91.876 E | 2930m |
| NRC-AA-7593 | *Apis laboriosa* | Female(worker) | Nyaton Kitnya | 2018 | 7 | 3-5 | India | 27.591 N | 91.876 E | 2930m |
| NRC-AA-7594 | *Apis laboriosa* | Female(worker) | Nyaton Kitnya | 2018 | 7 | 3-5 | India | 27.591 N | 91.876 E | 2930m |
| NRC-AA-7595 | *Apis laboriosa* | Female(worker) | Nyaton Kitnya | 2018 | 7 | 3-5 | India | 27.591 N | 91.876 E | 2930m |
| NRC-AA-7596 | *Apis laboriosa* | Female(worker) | Nyaton Kitnya | 2018 | 7 | 3-5 | India | 27.591 N | 91.876 E | 2930m |
| NRC-AA-7597 | *Apis laboriosa* | Female(worker) | Nyaton Kitnya | 2018 | 7 | 3-5 | India | 27.591 N | 91.876 E | 2930m |
| NRC-AA-7598 | *Apis laboriosa* | Female(worker) | Nyaton Kitnya | 2018 | 7 | 3-5 | India | 27.591 N | 91.876 E | 2930m |
| NRC-AA-7599 | *Apis laboriosa* | Female(worker) | Nyaton Kitnya | 2018 | 7 | 4 | India | 27.582 N | 91.868 E | 2769m |
| NRC-AA-7600 | *Apis laboriosa* | Female(worker) | Nyaton Kitnya | 2018 | 7 | 4 | India | 27.582 N | 91.868 E | 2769m |
| NRC-AA-7601 | *Apis laboriosa* | Female(worker) | Nyaton Kitnya | 2018 | 7 | 4 | India | 27.582 N | 91.868 E | 2769m |
| NRC-AA-7602 | *Apis laboriosa* | Female(worker) | Nyaton Kitnya | 2018 | 7 | 4 | India | 27.582 N | 91.868 E | 2769m |
| NRC-AA-7603 | *Apis laboriosa* | Female(worker) | Nyaton Kitnya | 2018 | 7 | 4 | India | 27.582 N | 91.868 E | 2769m |
| NRC-AA-7604 | *Apis laboriosa* | Female(worker) | Nyaton Kitnya | 2018 | 7 | 4 | India | 27.582 N | 91.868 E | 2769m |
| NRC-AA-7605 | *Apis laboriosa* | Female(worker) | Nyaton Kitnya | 2018 | 7 | 4 | India | 27.582 N | 91.868 E | 2769m |
| NRC-AA-7606 | *Apis laboriosa* | Female(worker) | Nyaton Kitnya | 2018 | 7 | 4 | India | 27.582 N | 91.868 E | 2769m |
| NRC-AA-7607 | *Apis laboriosa* | Female(worker) | Nyaton Kitnya | 2018 | 7 | 4 | India | 27.582 N | 91.868 E | 2769m |
| NRC-AA-7608 | *Apis laboriosa* | Female(worker) | Nyaton Kitnya | 2018 | 7 | 4 | India | 27.582 N | 91.868 E | 2769m |
| NRC-AA-7609 | *Apis laboriosa* | Female(worker) | Nyaton Kitnya | 2018 | 7 | 5 | India | 27.621 N | 91.861 E | 3649m |
| NRC-AA-7610 | *Apis laboriosa* | Female(worker) | Nyaton Kitnya | 2018 | 7 | 5 | India | 27.621 N | 91.861 E | 3649m |
| NRC-AA-7611 | *Apis laboriosa* | Female(worker) | Nyaton Kitnya | 2018 | 7 | 5 | India | 27.621 N | 91.861 E | 3649m |
| NRC-AA-7612 | *Apis laboriosa* | Female(worker) | Nyaton Kitnya | 2018 | 7 | 5 | India | 27.621 N | 91.861 E | 3649m |
| NRC-AA-7613 | *Apis laboriosa* | Female(worker) | Nyaton Kitnya | 2018 | 7 | 5 | India | 27.621 N | 91.861 E | 3649m |
| NRC-AA-7614 | *Apis laboriosa* | Female(worker) | Nyaton Kitnya | 2018 | 7 | 5 | India | 27.621 N | 91.861 E | 3649m |
| NRC-AA-7615 | *Apis laboriosa* | Female(worker) | Nyaton Kitnya | 2018 | 7 | 5 | India | 27.621 N | 91.861 E | 3649m |
| NRC-AA-7616 | *Apis laboriosa* | Female(worker) | Nyaton Kitnya | 2018 | 7 | 5 | India | 27.621 N | 91.861 E | 3649m |
| NRC-AA-7617 | *Apis laboriosa* | Female(worker) | Nyaton Kitnya | 2018 | 7 | 5 | India | 27.621 N | 91.861 E | 3649m |
| NRC-AA-7618 | *Apis laboriosa* | Female(worker) | Nyaton Kitnya | 2018 | 7 | 5 | India | 27.623 N | 91.862 E | 2849m |
| NRC-AA-7619 | *Apis laboriosa* | Female(worker) | Nyaton Kitnya | 2018 | 7 | 5 | India | 27.623 N | 91.862 E | 2849m |
| NRC-AA-7620 | *Apis laboriosa* | Female(worker) | Nyaton Kitnya | 2018 | 7 | 5 | India | 27.623 N | 91.862 E | 2849m |
| NRC-AA-7621 | *Apis laboriosa* | Female(worker) | Nyaton Kitnya | 2018 | 7 | 5 | India | 27.623 N | 91.862 E | 2849m |
| NRC-AA-7622 | *Apis laboriosa* | Female(worker) | Nyaton Kitnya | 2018 | 7 | 5 | India | 27.623 N | 91.862 E | 2849m |
| NRC-AA-7623 | *Apis laboriosa* | Female(worker) | Nyaton Kitnya | 2018 | 7 | 5 | India | 27.623 N | 91.862 E | 2849m |
| NRC-AA-7624 | *Apis laboriosa* | Female(worker) | Nyaton Kitnya | 2018 | 7 | 5 | India | 27.623 N | 91.862 E | 2849m |
| NRC-AA-7625 | *Apis laboriosa* | Female(worker) | Nyaton Kitnya | 2018 | 7 | 5 | India | 27.623 N | 91.862 E | 2849m |
| NRC-AA-7626 | *Apis laboriosa* | Female(worker) | Nyaton Kitnya | 2018 | 7 | 4 | India | 27.575 N | 91.866 E | 2730m |
| NRC-AA-7627 | *Apis laboriosa* | Female(worker) | Nyaton Kitnya | 2018 | 7 | 4 | India | 27.575 N | 91.866 E | 2730m |
| NRC-AA-7628 | *Apis laboriosa* | Female(worker) | Nyaton Kitnya | 2018 | 7 | 4 | India | 27.575 N | 91.866 E | 2730m |
| NRC-AA-7629 | *Apis laboriosa* | Female(worker) | Nyaton Kitnya | 2018 | 7 | 4 | India | 27.575 N | 91.866 E | 2730m |
| NRC-AA-7630 | *Apis laboriosa* | Female(worker) | Prasad Bhatta (CPB) | | | | Nepal | | | |
| NRC-AA-7631 | *Apis laboriosa* | Female(worker) | Prasad Bhatta (CPB) | | | | Nepal | | | |
| NRC-AA-7632 | *Apis laboriosa* | Female(worker) | Prasad Bhatta (CPB) | | | | Nepal | | | |

| | | | | | | | | | | |
|---|---|---|---|---|---|---|---|---|---|---|
| NRC-AA-7633 | *Apis laboriosa* | Female(worker) | Prasad Bhatta | (CPB) | | | Nepal | | | |
| NRC-AA-7634 | *Apis laboriosa* | Female(worker) | Prasad Bhatta | (CPB) | | | Nepal | | | |
| NRC-AA-7635 | *Apis laboriosa* | Female(worker) | Prasad Bhatta | (CPB) | | | Nepal | | | |
| NRC-AA-7636 | *Apis laboriosa* | Female(worker) | Prasad Bhatta | (CPB) | | | Nepal | | | |
| NRC-AA-7637 | *Apis laboriosa* | Female(worker) | Prasad Bhatta | (CPB) | | | Nepal | | | |
| NRC-AA-7638 | *Apis laboriosa* | Female(worker) | Prasad Bhatta | (CPB) | | | Nepal | | | |
| NRC-AA-7639 | *Apis laboriosa* | Female(worker) | La Quang Trung | 1996 | 6 | 7 | Vietnam | | | |
| NRC-AA-7640 | *Apis laboriosa* | Female(worker) | La Quang Trung | 1996 | 6 | 7 | Vietnam | | | |
| NRC-AA-7641 | *Apis laboriosa* | Female(worker) | La Quang Trung | 1996 | 6 | 7 | Vietnam | | | |
| NRC-AA-7642 | *Apis laboriosa* | Female(worker) | La Quang Trung | 1996 | 6 | 7 | Vietnam | | | |
| NRC-AA-7643 | *Apis laboriosa* | Female(worker) | La Quang Trung | 1996 | 6 | 7 | Vietnam | | | |
| NRC-AA-7644 | *Apis laboriosa* | Female(worker) | La Quang Trung | 1996 | 6 | 7 | Vietnam | | | |
| NRC-AA-7645 | *Apis laboriosa* | Female(worker) | La Quang Trung | 1996 | 6 | 7 | Vietnam | | | |
| NRC-AA-7646 | *Apis laboriosa* | Female(worker) | La Quang Trung | 1996 | 6 | 7 | Vietnam | | | |
| NRC-AA-7647 | *Apis laboriosa* | Female(worker) | | 1996 | | | Vietnam | | | |
| NRC-AA-7648 | *Apis laboriosa* | Female(worker) | | 1996 | | | Vietnam | | | |
| NRC-AA-7649 | *Apis laboriosa* | Female(worker) | | 1996 | | | Vietnam | | | |
| NRC-AA-7650 | *Apis laboriosa* | Female(worker) | | 1996 | | | Vietnam | | | |
| NRC-AA-7651 | *Apis laboriosa* | Female(worker) | | 1996 | | | Vietnam | | | |
| NRC-AA-7652 | *Apis laboriosa* | Female(worker) | Savita | 2023 | 3 | 15 | India | 22.461 N | 78.419 E | 980m |
| NRC-AA-7653 | *Apis laboriosa* | Female(worker) | Savita | 2023 | 3 | 15 | India | 22.461 N | 78.419 E | 980m |
| NRC-AA-7654 | *Apis laboriosa* | Female(worker) | Savita | 2023 | 3 | 15 | India | 22.461 N | 78.419 E | 980m |
| NRC-AA-7655 | *Apis laboriosa* | Female(worker) | Savita | 2023 | 3 | 15 | India | 22.461 N | 78.419 E | 980m |
| NRC-AA-7656 | *Apis laboriosa* | Female(worker) | Savita | 2023 | 3 | 15 | India | 22.461 N | 78.419 E | 980m |
| NRC-AA-7657 | *Apis laboriosa* | Female(worker) | Savita | 2023 | 3 | 15 | India | 22.461 N | 78.419 E | 980m |
| NRC-AA-7658 | *Apis laboriosa* | Female(worker) | Nyaton Kitnya | 2023 | 3 | 16 | India | 13.07 N | 77.580 E | 930m |
| NRC-AA-7659 | *Apis laboriosa* | Female(worker) | Nyaton Kitnya | 2023 | 3 | 16 | India | 13.07 N | 77.580 E | 930m |
| NRC-AA-7660 | *Apis laboriosa* | Female(worker) | Nyaton Kitnya | 2023 | 3 | 16 | India | 13.07 N | 77.580 E | 930m |
| NRC-AA-7661 | *Apis laboriosa* | Female(worker) | Nyaton Kitnya | 2023 | 3 | 16 | India | 13.07 N | 77.580 E | 930m |
| NRC-AA-8003 | *Apis laboriosa* | Female(worker) | itnya, Anne-Sop | 2023 | 6 | 10 | India | 26.954 | 95.652 | 1060m |
| NRC-AA-8004 | *Apis laboriosa* | Female(worker) | itnya, Anne-Sop | 2023 | 6 | 10 | India | 26.954 | 95.652 | 1060m |
| NRC-AA-8005 | *Apis laboriosa* | Female(worker) | itnya, Anne-Sop | 2023 | 6 | 10 | India | 26.954 | 95.652 | 1060m |
| NRC-AA-8006 | *Apis laboriosa* | Female(worker) | itnya, Anne-Sop | 2023 | 6 | 10 | India | 26.954 | 95.652 | 1060m |
| NRC-AA-8007 | *Apis laboriosa* | Female(worker) | itnya, Anne-Sop | 2023 | 6 | 10 | India | 26.954 | 95.652 | 1060m |
| NRC-AA-8008 | *Apis laboriosa* | Female(worker) | itnya, Anne-Sop | 2023 | 6 | 10 | India | 26.954 | 95.652 | 1060m |
| NRC-AA-8009 | *Apis laboriosa* | Female(worker) | itnya, Anne-Sop | 2023 | 6 | 10 | India | 26.954 | 95.652 | 1060m |
| NRC-AA-8010 | *Apis laboriosa* | Female(worker) | itnya, Anne-Sop | 2023 | 6 | 10 | India | 26.954 | 95.652 | 1060m |
| NRC-AA-8011 | *Apis laboriosa* | Female(worker) | itnya, Anne-Sop | 2023 | 6 | 10 | India | 26.954 | 95.652 | 1060m |
| NRC-AA-8012 | *Apis laboriosa* | Female(worker) | itnya, Anne-Sop | 2023 | 6 | 10 | India | 26.954 | 95.652 | 1060m |
| NRC-AA-8058 | *Apis laboriosa* | Male (drone) | itnya, Anne-Sop | 2023 | 6 | 10 | India | 26.954 | 95.652 | 1060m |
| NRC-AA-8059 | *Apis laboriosa* | Male (drone) | itnya, Anne-Sop | 2023 | 6 | 10 | India | 26.954 | 95.652 | 1060m |
| NRC-AA-8060 | *Apis laboriosa* | Male (drone) | itnya, Anne-Sop | 2023 | 6 | 10 | India | 26.954 | 95.652 | 1060m |
| NRC-AA-8061 | *Apis laboriosa* | Male (drone) | itnya, Anne-Sop | 2023 | 6 | 10 | India | 26.954 | 95.652 | 1060m |
| NRC-AA-8062 | *Apis laboriosa* | Male (drone) | itnya, Anne-Sop | 2023 | 6 | 10 | India | 26.954 | 95.652 | 1060m |
| NRC-AA-8063 | *Apis laboriosa* | Male (drone) | itnya, Anne-Sop | 2023 | 6 | 10 | India | 26.954 | 95.652 | 1060m |
| NRC-AA-8064 | *Apis laboriosa* | Male (drone) | itnya, Anne-Sop | 2023 | 6 | 10 | India | 26.954 | 95.652 | 1060m |
| NRC-AA-8065 | *Apis laboriosa* | Male (drone) | itnya, Anne-Sop | 2023 | 6 | 10 | India | 26.954 | 95.652 | 1060m |
| NRC-AA-8066 | *Apis laboriosa* | Male (drone) | itnya, Anne-Sop | 2023 | 6 | 10 | India | 26.954 | 95.652 | 1060m |
| NRC-AA-8067 | *Apis laboriosa* | Male (drone) | itnya, Anne-Sop | 2023 | 6 | 10 | India | 26.954 | 95.652 | 1060m |
| NRC-AA-8099 | *Apis laboriosa* | Female(worker) | | | | | Nepal | | | |
| NRC-AA-8100 | *Apis laboriosa* | Female(worker) | | | | | Nepal | | | |
| NRC-AA-8101 | *Apis laboriosa* | Female(worker) | | | | | Nepal | | | |

| | | | | | | | | | | |
|---|---|---|---|---|---|---|---|---|---|---|
| NRC-AA-8102 | *Apis laboriosa* | Female(worker) | | | | | Nepal | | | |
| NRC-AA-8103 | *Apis laboriosa* | Female(worker) | | | | | Nepal | | | |
| NRC-AA-8104 | *Apis laboriosa* | Female(worker) | | | | | Nepal | | | |
| NRC-AA-8105 | *Apis laboriosa* | Female(worker) | | | | | Nepal | | | |
| NRC-AA-8106 | *Apis laboriosa* | Female(worker) | | | | | Nepal | | | |
| NRC-AA-8107 | *Apis laboriosa* | Female(worker) | | | | | Nepal | | | |
| NCBS-BF244 | *Apis laboriosa* | Female(worker) | itnya and Prabh | 2017 | 9 | 7 | India | ',12.498' | 2,33.229 | 1162m |
| NCBS-BF245 | *Apis laboriosa* | Female(worker) | Nyaton Kitnya | 2017 | 10 | 12 | India | ' 59,60.24 | ' 32,26.7 | 1500m |
| NCBS-BF150 | *Apis laboriosa* | Female(worker) | GU/Uni Würzbu | 2016 | 6 | 11 | India | ###### | 95.022 | 289 |
| NCBS-BF151 | *Apis laboriosa* | Female(worker) | GU/Uni Würzbu | 2016 | 6 | 11 | India | ###### | 95.022 | 289 |
| NCBS-BF152 | *Apis laboriosa* | Female(worker) | GU/Uni Würzbu | 2016 | 6 | 11 | India | ###### | 95.022 | 289 |
| NCBS-BF153 | *Apis laboriosa* | Female(worker) | GU/Uni Würzbu | 2016 | 6 | 11 | India | ###### | 95.022 | 289 |
| NCBS-BF154 | *Apis laboriosa* | Female(worker) | GU/Uni Würzbu | 2016 | 5 | 26 | India | ###### | 92.414 | 2176 |
| NCBS-BF155 | *Apis laboriosa* | Female(worker) | GU/Uni Würzbu | 2016 | 5 | 26 | India | ###### | 92.414 | 2176 |
| NCBS-BF156 | *Apis laboriosa* | Female(worker) | GU/Uni Würzbu | 2016 | 5 | 26 | India | ###### | 92.414 | 2176 |
| NCBS-BF157 | *Apis laboriosa* | Female(worker) | GU/Uni Würzbu | 2016 | 5 | 26 | India | ###### | 92.277 | 1507 |
| NCBS-BF158 | *Apis laboriosa* | Female(worker) | GU/Uni Würzbu | 2016 | 5 | 26 | India | ###### | 92.277 | 1507 |
| NCBS-BF159 | *Apis laboriosa* | Female(worker) | GU/Uni Würzbu | 2016 | 5 | 26 | India | ###### | 92.277 | 1507 |
| NCBS-BF160 | *Apis laboriosa* | Female(worker) | GU/Uni Würzbu | 2016 | 5 | 26 | India | ###### | 92.277 | 1507 |
| NCBS-BF161 | *Apis laboriosa* | Female(worker) | GU/Uni Würzbu | 2016 | 5 | 26 | India | ###### | 92.277 | 1507 |
| NCBS-BF162 | *Apis laboriosa* | Female(worker) | GU/Uni Würzbu | 2016 | 5 | 26 | India | ###### | 92.248 | 1680 |
| NCBS-BF163 | *Apis laboriosa* | Female(worker) | GU/Uni Würzbu | 2016 | 5 | 26 | India | ###### | 92.248 | 1680 |
| NCBS-BF164 | *Apis laboriosa* | Female(worker) | GU/Uni Würzbu | 2016 | 5 | 27 | India | ###### | 91.991 | 2650 |
| NCBS-BF165 | *Apis laboriosa* | Female(worker) | GU/Uni Würzbu | 2016 | 5 | 27 | India | ###### | 91.991 | 2650 |
| NCBS-BF166 | *Apis laboriosa* | Female(worker) | GU/Uni Würzbu | 2016 | 5 | 28 | India | ###### | 91.862 | 2800 |
| NCBS-BF167 | *Apis laboriosa* | Female(worker) | GU/Uni Würzbu | 2016 | 5 | 28 | India | ###### | 91.862 | 2800 |
| NCBS-BF168 | *Apis laboriosa* | Female(worker) | GU/Uni Würzbu | 2016 | 5 | 28 | India | ###### | 91.871 | 3030 |
| NCBS-BF169 | *Apis laboriosa* | Female(worker) | GU/Uni Würzbu | 2016 | 5 | 28 | India | ###### | 91.871 | 3030 |
| NCBS-BF170 | *Apis laboriosa* | Female(worker) | GU/Uni Würzbu | 2016 | 5 | 28 | India | ###### | 91.871 | 3030 |
| NCBS-BF171 | *Apis laboriosa* | Female(worker) | GU/Uni Würzbu | 2016 | 5 | 28 | India | ###### | 91.871 | 3030 |
| NCBS-BF172 | *Apis laboriosa* | Female(worker) | GU/Uni Würzbu | 2016 | 5 | 29 | India | ###### | 91.998 | 3328 |
| NCBS-BF173 | *Apis laboriosa* | Female(worker) | GU/Uni Würzbu | 2016 | 5 | 30 | India | ###### | 91.923 | 3133 |
| NCBS-BF174 | *Apis laboriosa* | Female(worker) | GU/Uni Würzbu | 2016 | 5 | 30 | India | ###### | 91.923 | 3133 |
| NCBS-BF175 | *Apis laboriosa* | Female(worker) | GU/Uni Würzbu | 2016 | 5 | 31 | India | ###### | 92.109 | 3250 |

| | | | |
|---|---|---|---|
| Nyaton Kitnya | 2018/02/27 | | |
| Nyaton Kitnya | 2018/02/27 | | |
| Nyaton Kitnya | 2018/02/27 | | |
| Nyaton Kitnya | 2018/02/27 | | |
| Nyaton Kitnya | 2018/02/27 | | |
| Nyaton Kitnya | 2018/02/27 | | |
| Nyaton Kitnya | 2018/02/27 | | |
| Nyaton Kitnya | 2018/02/27 | | |
| Nyaton Kitnya | 2018/02/27 | | |
| Nyaton Kitnya | 2018/02/27 | | |
| Nyaton Kitnya | 2018/02/27 | | |
| Nyaton Kitnya | 2018/02/27 | | |
| Nyaton Kitnya | 2018/02/27 | | |
| Nyaton Kitnya | 2018/02/27 | | |
| Nyaton Kitnya | 2018/02/27 | | |
| Nyaton Kitnya | 2018/02/27 | | |
| Nyaton Kitnya | 2018/02/27 | | |
| Nyaton Kitnya | 2018/02/27 | | |
| Nyaton Kitnya | 2018/02/27 | | |
| Nyaton Kitnya | 2018/02/27 | | |
| Nyaton Kitnya | 2018/02/27 | | |
| Streinzer Martin | 2018 | Streinzer Martin | 2018 |
| Streinzer Martin | 2018 | Streinzer Martin | 2018 |
| Streinzer Martin | 2018 | Streinzer Martin | 2018 |
| Streinzer Martin | 2018 | Streinzer Martin | 2018 |
| Streinzer Martin | 2018 | Streinzer Martin | 2018 |

| | | | |
|---|---|---|---|
| Streinzer Martin | 2018 | Streinzer Martin | 2018 |
| Streinzer Martin | 2018 | Streinzer Martin | 2018 |
| Streinzer Martin | 2018 | Streinzer Martin | 2018 |
| Streinzer Martin | 2018 | Streinzer Martin | 2018 |
| Streinzer Martin | 2018 | Streinzer Martin | 2018 |
| Streinzer Martin | 2018 | Streinzer Martin | 2018 |
