## Supplementary file2 for "Taxonomic Revision and Identification Keys for the Giant Honey Bees": Supplimentary file2_Taxonomic Revision and Identification Keys for the Giant Honey Bees_submitted.pdf

Supplementary file 2: Taxonomic Revision and Identification Keys for the Giant Honey Bees_submitted

S2: A: Variation of ocellocular distance in *Megapis* worker. Arrow indicates the ocellocular distance (OOD) which is wider in *laboriosa* as compared to other three taxa: A. *binghami*; B. *breviligula*; C. *dorsata*; D. *laboriosa*.

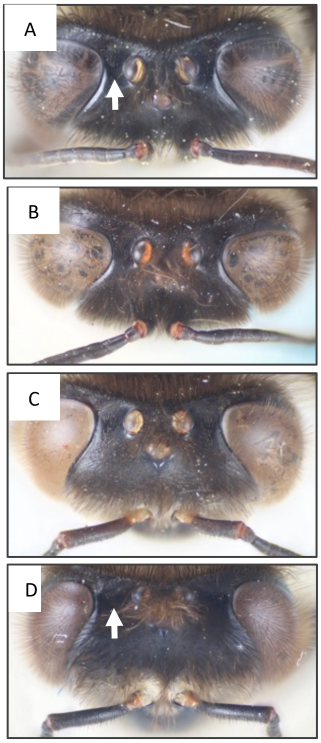

S2: B: Variation in projection of compound eyes toward centre of head in *Megapis* workers. A. *binghami*; B. *breviligula*; C. *dorsata*; D. *laboriosa*. Arrow indicates the extension of compound eyes towards the center of the head. Note that in *binghami* and *breviligula*, the compound eyes seem to be largest (with the greatest extension inwards). The compound eyes of *laboriosa* project the least towards toward the centre of the head.

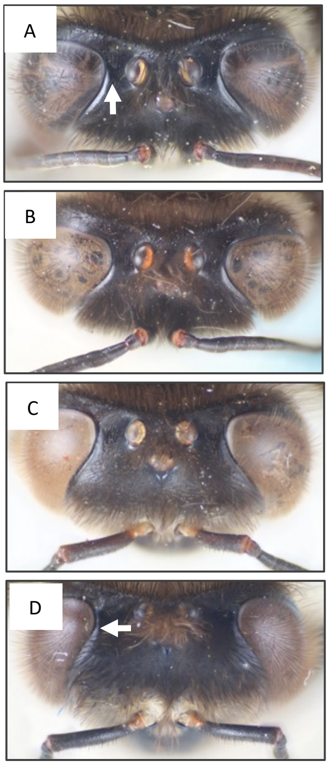

S2: C: Malar length in *Megapis* workers. A. *binghami*; B. *breviligula*; C. *dorsata*; D. *laboriosa*. The malar area is longer in *laboriosa* than in the other three taxa.

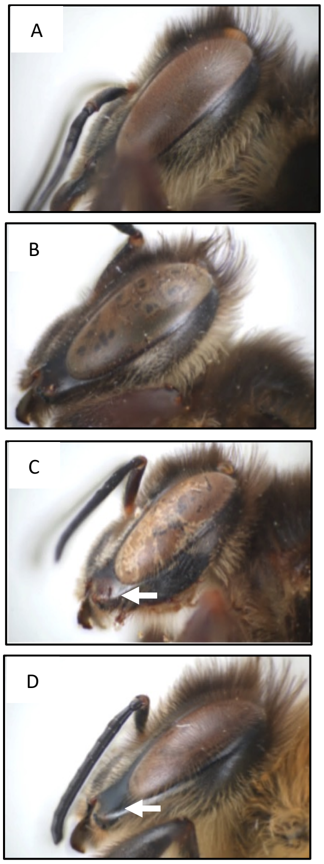
